# Expression of fatty acid synthase genes and their role in development and arboviral infection of *Aedes aegypti*

**DOI:** 10.1101/2022.02.14.480456

**Authors:** Nunya Chotiwan, Carlos A. Brito-Sierra, Gabriella Ramirez, Elena Lian, Catherine A. Hill, Rushika Perera

**Affiliations:** Department of Microbiology, Immunology and Pathology, Colorado State University, Fort Collins, Colorado, USA; Department of Entomology, Purdue University, West Lafayette, Illinois, USA; Purdue Institute of Inflammation, Immunology and Infectious Disease, Purdue University, West Lafayette, Indiana, United States

**Keywords:** *Aedes aegypti*, Aag2 cells, fatty acid synthase, FAS, lipid, lipid metabolism, dengue virus, AaegL5 genome assembly

## Abstract

**Background:** Fatty acids are the building blocks of complex lipids essential for living organisms. In mosquitoes, fatty acids are involved in cell membrane production, energy conservation and expenditure, innate immunity, development, and reproduction. Fatty acids are synthesized by a multifunctional enzyme complex called fatty acid synthase (FAS). Several paralogues of FAS were found in the *Aedes aegypti* (*Ae. aegypti*) mosquito. However, the molecular characteristics and the expression of some of these paralogues have not been investigated.

**Methods:** Genome assemblies of *Ae. aegypti* were analyzed and orthologues of human FAS were identified. Phylogenetic analysis and *in silico* molecular characterization were performed to identify the functional domains of the *Ae. aegypti* FAS (*Aa*FAS). Quantitative analysis and loss-of-function experiments were performed to determine the significance of different *Aa*FAS transcripts in various stages of development, expression following different diets and the impact of *Aa*FAS on dengue virus, serotype 2 (DENV2) infection and transmission.

**Results:** We identified seven putative FAS genes in the *Ae. aegypti* genome assembly, based on nucleotide similarity to the FAS proteins (tBLASTn) of humans, other mosquitoes and invertebrates. Bioinformatics and molecular analyses suggested that only five of the FAS genes produce mRNA and therefore represent complete gene models. Expression levels of *Aa*FAS varied among developmental stages and between male and female *Ae. aegypti*. Quantitative analyses revealed that expression of *Aa*FAS1, the putative orthologue of the human FAS, was highest in adult females. Transient knockdown (KD) of *Aa*FAS1 did not induce a complete compensation by other *Aa*FAS genes but limited DENV2 infection of Aag2 cells in culture and the midgut of the mosquito.

**Conclusion:** *Aa*FAS1 is the predominant *Aa*FAS in the adult mosquitoes. It has the highest amino acid similarity to human FAS and contains all enzymatic domains typical of human FAS. *Aa*FAS1 also facilitated DENV2 replication in both cell culture and in mosquito midguts. Our data suggest that *Aa*FAS1 may play a role in transmission of dengue viruses, and could represent a target for intervention strategies.

## Background

Fatty acid synthase (FAS) is a multifunctional enzyme catalyzing more than 40 steps in the *de novo* fatty acid biosynthesis pathway (1) (2). It contains seven catalytic and three non-catalytic domains which condense, reduce and dehydrate the three-carbon substrate, malonyl-CoA, into 16 to 18-carbon fatty acids. These fatty acids are essential building blocks of complex lipids, such as phosphoglycerolipids, glycerolipids and sphingolipids, which are components of cellular membranes and storage lipids, and function as signaling molecules, respectively.

In mosquitoes, fatty acids also play roles in innate immunity, reproduction, development, and flight (3–5). Fatty acids can be acquired or synthesized in both larvae and adult stages. Neonate larvae acquire lipids through the maternal deposition in eggs (6–8) and through consumption of aquatic diets such as diatoms and algae, which are the primary source of polyunsaturated fatty acids (9). Fatty acids from larval stages can be transferred to the adult stage and some can be deposited in eggs of the first gonotrophic cycle (10). Adult mosquitoes possess enzymes for *de novo* synthesis and modification of fatty acids from both sugar (carbohydrate-enriched) and blood (protein-enriched) meals (7, 9, 11). In the female, fatty acid synthesis is important for metabolism and production of eggs. Transient knockdown (KD) of acetyl-CoA carboxylase (ACC) and fatty acid synthase (FAS), two key enzymes in the *de novo* fatty acid biosynthesis pathway, led to significantly lower egg production in the first gonotrophic cycle (12). In addition, eggs produced by ACC-KD mosquitoes lacked eggshells and were nonviable (12).

Apart from its importance to mosquito biology, studies suggest FAS also plays a supportive role for several arboviral infections in both mammalian and mosquito cells (13–16). It is known that several RNA viruses induce expansion and rearrangement of host cell membranes to support viral genome replication and assembly (17–19). Studies have shown that FAS facilitates the production of dengue virus serotype 2 (DENV2) infection in both human and mosquito cells, potentially by providing the building blocks for this membrane expansion event (13, 15). Lastly, studies also reported the elevation of fatty acid abundance in C6/36 (*Aedes albopictus*) cells, and in the *Aedes aegypti* (*Ae. aegypti*) mosquito midgut during DENV2 infection (15, 20). These findings suggest that fatty acids are essential for the physiological function of mosquitoes, and support DENV2 infection of the mosquito.

Currently, understanding of FAS in mosquitoes and its role in pathogen transmission by the mosquito vector is limited. Here, we describe the molecular and functional characterization of the FAS gene family from *Ae. aegypti* (*Aa*FAS). We identified seven putative *Aa*FAS genes (*Aa*FAS 1-6 and *Aa*FAS-like) in the AaegL5 genome assembly, characterized the expression of these genes during mosquito development and following consumption of different diets. *Aa*FAS1 had the highest amino acid similarity to human FAS and was the predominant transcript. We investigated the role of *Aa*FAS1 in DENV2 infection in mosquito cells and live mosquitoes using gene KD. We observed a significant reduction of DENV2 replication following *Aa*FAS1-KD in *Ae. aegypti* cells and a transient reduction of infection in *Ae. aegypti* midguts at early time points post-infectious blood meal. These results provide insights to the molecular characteristic of *Aa*FASs and their role during *Ae. aegypti* development, food source acquisition and arbovirus infection.

## Methods

### Alignments, conserved motifs and phylogenetic tree

Putative *Aa*FAS sequences from the AaegL5 genome assembly were blasted against the AaegL3 genome assembly retrieved from VectorBase using tBLASTn (21–23). FAS sequences of *Anopheles gambiae, Drosophila melanogaster, Apis mellifera, Homo sapiens*, *Mus musculus* and *Saccharomyces cerevisiae* were aligned with putative *Aa*FAS sequences using ClustalW (24). mRNA sequences were retrieved from NCBI and manually curated to confirm the intron/exon boundaries. Conserved FAS motifs were identified by global alignment of vertebrate, invertebrate and yeast proteins using the Clustal Omega and Jalview 2.11.1.5 (accession numbers are shown in Table S1) (25) and conserved amino acids associated with catalytic domains of functional FAS were identified by comparison to sequences reported in published studies (26). Individual amino acid alignments were also performed between FAS-AaegL5 and FAS-AaegL3 using ClustalW to identify improvements in AaegL5 models.

A Bayesian inference of phylogeny was performed using the amino acid sequence of FAS from *Ae. aegypti, Anopheles gambiae, Drosophila melanogaster, Apis mellifera, Mus musculus* and *Homo sapiens.* Yeast Kexin was used as an outgroup. A sequence alignment with ClustalW was performed prior to tree construction in phylogeny.fr. The substitution model used for the Bayesian inference was Blosum62 and the Markov Chain Monte Carlo parameters included 100,000 generations with sampling every 10 generations, discarding the first 250 trees. The resulting tree was annotated and curated in iTOL.

### Annotation of protein domains in *Ae. aegypti* FAS genes

*Aa*FAS amino acid sequences were aligned against the human FAS (NP_004095.4, NCBI) using Clustal Omega (27) to identify the seven catalytic and three noncatalytic domains associated with mammalian FAS. The alignment results were viewed using MView tool (28). Motifs in the human FAS were identified based on Pfam 31.0 (29) and conserved domains in *Aa*FAS genes were identified by comparative analyses.

### Mosquito rearing

Larvae and adults of *Aedes aegypti* strain Chetumal, originally collected from Yucatan Peninsula in Mexico, were reared on fish food and on 10% sucrose solution, respectively and adults maintained under constant conditions of 28°C, 80% relative humidity (30).

### Blood feeding

Twenty-four hours prior to blood feeding, mosquitoes were starved for 4 hours by removal of sucrose solution. Defibrinated sheep blood (Colorado Veterinarian Product) was mixed with 1mM ATP and placed in an artificial membrane feeder warmed by a 37°C water jacket. Mosquitoes were allowed to feed for 45-60 minutes. Fully engorged mosquitoes were sorted and reared on 10% sucrose solution and water.

### Generating long double-stranded RNA

Long double-stranded RNA (dsRNA) was generated from *Ae. aegypti* mosquito total RNA. Primers were designed to amplify an ∼500 bp region of the gene of interest (Table S2). cDNA was generated by reverse transcription (RT) using specific reverse primers and SuperScript III Reverse Transcriptase (Invitrogen). Polymerase chain reaction (PCR) was performed using specific primers containing a 5’ T7 promotor sequence adapted to both forward and reverse primers and Taq polymerase (NEB). PCR products were purified using the GeneJET PCR Purification kit (Thermo Scientific) and in vitro transcription was performed using the MEGAscript T7 kit (Invitrogen) and incubation at 37°C for 12 hours. Following incubation, the product was heated to 75°C for 5 minutes and slowly cooled to room temperature for 4 hours to dsRNA annealing. Next, dsRNA was treated with DNase (NEB) and purified by phenol-chloroform extraction followed by ethanol precipitation and the purified dsRNA was stored at - 80°C.

### dsRNA knockdown of AaFAS1 in *Ae. aegypti* moquitoes

dsRNA was introduced via intrathoracic (IT) injection of adult females at 3 to 4 days post-eclosion (31). Mosquitoes were anesthetized at 4°C on a cold plate. Glass needles were prepared with a vertical pipette puller (P-30, Sutter Instrument Co., Novato, CA) and mosquitoes were IT injected with 3 µg/µl of dsRNA in an injection volume of 69 nl, twice (total of ∼400 ng of dsRNA) using a Nanojet II (Drummond Scientific Company, Broomall, PA). Injected mosquitoes were fed on sucrose solution or blood and reared at 28°C, 80% relative humidity for 17 days post-injection.

### dsRNA knockdown of *Aa*FAS1 gene and DENV2 infection of Aag2 cells

dsRNA KD was performed in RNA interference-competent *Ae. aegypti* (Aag2) cells. Aag2 cells were cultured in Schneider’s insect medium (Sigma-Aldrich) supplemented with 2 mM L-glutamine, 1% non-essential amino acids and 10% FBS. The cells were seeded in a 48-well plate at 50,000 cells/well for 24 hours, and subsequently transfected with 260 ng of dsRNA mixed with TransIT-2020 Reagent (Mirus) following the manufacturer’s protocol. New medium with 2% FBS was replaced at 6 hours post-transfection. Cell viability assays were performed at 2 days post-transfection using resazurin assay.

KD cells were infected with infectious DENV2 expressing a luciferase reporter (DEN-Luc) supplied by C. Rice, Rockefeller University. Cell culture medium was replaced with 300 µl of DEN-Luc supernatant at 48 hours post dsRNA transfection, and cells were incubated at 28°C without CO_2_. Virus supernatant was removed at 24 hours post-infection, and cells were lysed, and luciferase activity was read using the Luciferase Assay System (Promega) as per manufacturer protocol.

### Gene expression analyses

Total RNA was extracted from dissected midgut or whole mosquito by TRIzol (Life Tech) and cDNA was produced via reverse transcription using random primers (Life Tech) and SuperScript III Reverse Transcriptase (Invitrogen). Approximately 400 ng of total cDNA was employed for quantitative PCR (qPCR) analyses. Gene-specific primers are listed in Table S3. β-actin was used as a reference gene. Relative *Aa*FAS gene expression was assessed by normalization to the levels of the β-actin gene (2^-ΔCt^). The comparative Ct (2^-ΔΔCt^) method was used to calculate the relative expression of *Aa*FAS following treatment compared to the control (32).

For RT-PCR assay, total RNA was treated with DNase I, RNase-free (1 U/µl) kit (ThermoFisher) prior to reverse transcription reaction. Purified cDNA was then amplified using Q5® High-Fidelity DNA Polymerase kit (New England BioLabs) with following condition, 98°C for 30 seconds, 35 cycles of 98°C for 10 seconds, 68°C and 72°C for 2 minutes and 30 seconds. primers are listed in Table S4.

### Virus infection of *Ae. aegypti* by infectious blood meal

DENV2 serotype 2 strain Jamaica-1409 (33) was cultured in C6/36 cells. Cells were infected with DENV2 at a multiplicity of infection of 0.01 and incubation at room temperature for 1 hour. Virus supernatant was removed, and infected cells were cultured in 5 ml total volume of L15 medium supplemented with 3% fetal bovine serum (FBS), 50 μg/ml penicillin-streptomycin, and 2 mM L-glutamine. Media was replaced at 7 days post-infection (dpi) and virus supernatant was harvested on 12-14 dpi and immediately used to prepare the infectious blood meal.

### Midgut dissection and plaque titration

Mosquito tissues were collected at multiple days post-exposure to the virus indicated in the figure legends. Isolated midguts or the mosquito carcass (remainder of the body without midgut) were placed separately into 2 ml safe-lock Eppendorf tubes (Eppendorf) containing 250 µl of mosquito diluent (1 × PBS supplemented with 20% FBS, 50 µg/ml Penicillin/Streptomycin (Gibco), 50 µg/ml Gentamycin (Gibco), and 2.5 µg/ml Amphotericin B (Gibco)) and a stainless-steel bead (34). Tissue was homogenized using a Retsch Mixer Mill MM400 at 24 cycles per minute, centrifuged at 15,000g for 5LJminutes at 4°C and supernatant was transferred to a new tube for plaque titration.

Plaque assay was performed on BHK-15 cells. Ten-fold serially diluted viral supernatant was absorbed on the confluent cell layer. After 45 minutes of absorption, cells were overlaid with 1x Minimum Essential Media (MEM), 1X agar supplemented with 2.5 % FBS, 25 µg/ml Penicillin/Streptomycin, 25 µg/ml Gentamycin, and 1.25 µg/ml Amphotericin B and the cells were incubated at 37°C with 5% CO_2_. Cells were stained with 0.033% neutral red (Sigma) in 1x PBS on day 5 post-infection and plaques were counted at 24 hour post-staining.

## Results

### Molecular Characterization of *Ae. aegypti* FAS genes

Seven putative *Aa*FAS genes models were obtained via manual annotation using the AaegL5 assembly (35). Previously, five candidate FAS genes (*Aa*FAS1-5), were identified based on the AaegL3 assembly of Nene et al., 2007 (23, 36), and of these, only *Aa*FAS1 and 2 have undergone functional studies (12). The AaegL5 assembly enabled identification of two additional candidate FAS genes (*Aa*FAS6 and *Aa*FAS-like). The corresponding mRNA sequences showing predicted intron/exon structure and initiation and stop codons are shown in Supplemental File 1. The *Aa*FAS1 gene model revealed a gene structure comprising 11 exons, while *Aa*FAS2 had 5 exons and *Aa*FAS3*-*5 had 6 exons (Fig. 1). The incomplete *Aa*FAS-like and *Aa*FAS6 gene models comprised 2 and 3 exons, respectively.

**Fig. 1.**
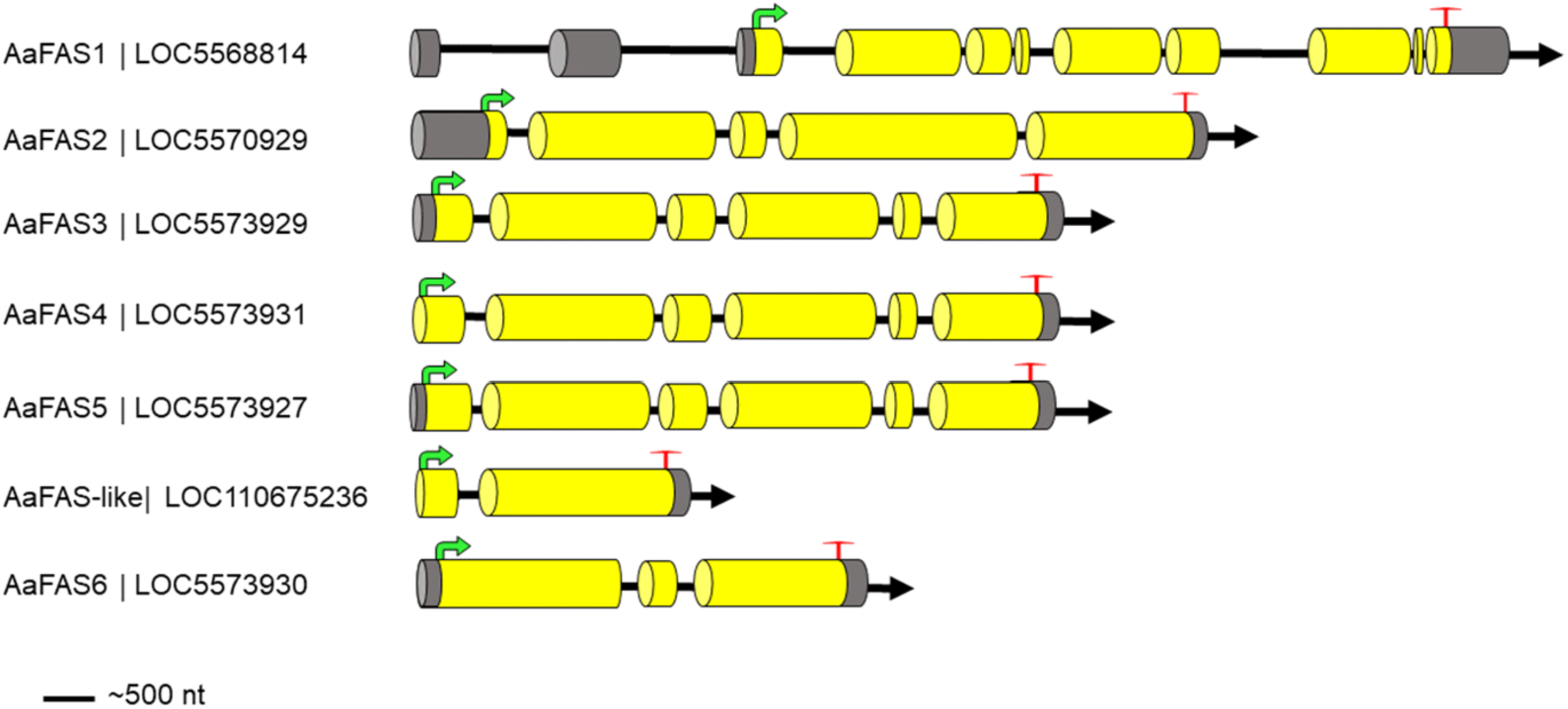
Schematic showing the predicted gene structure of the *Aa*FAS gene family. Exons are indicated by yellow cylindrical bars, 5’ and 3’ non-coding exons by dark grey shading, introns by a black line, start codon by green arrow and stop codon by red T.

The gene models for *Aa*FAS1-5 appear to be full length, with an average gene product length of 2,360 amino acids (Table 1). *Aa*FAS1-5 possessed features associated with functional FAS, including an initiation methionine, a stop codon and the functional catalytic motifs (DTACSS, EAH and GSVKS) important for ketoacyl synthesis as described by Beedessee *et al*. 2015 (26). Additionally, *Aa*FAS1-5 contained the YKELRLRGY motif conserved among the FAS genes of vertebrates and invertebrates, present in the polyketide synthase deshydratase domain (Fig. S1). *Aa*FAS3 lacked 6 amino acid residues in the 3’ terminus of exon 6 and a total of 127 non-synonymous substitutions were identified in this model as compared to its AaegL3 counterpart.

**Table 1.**
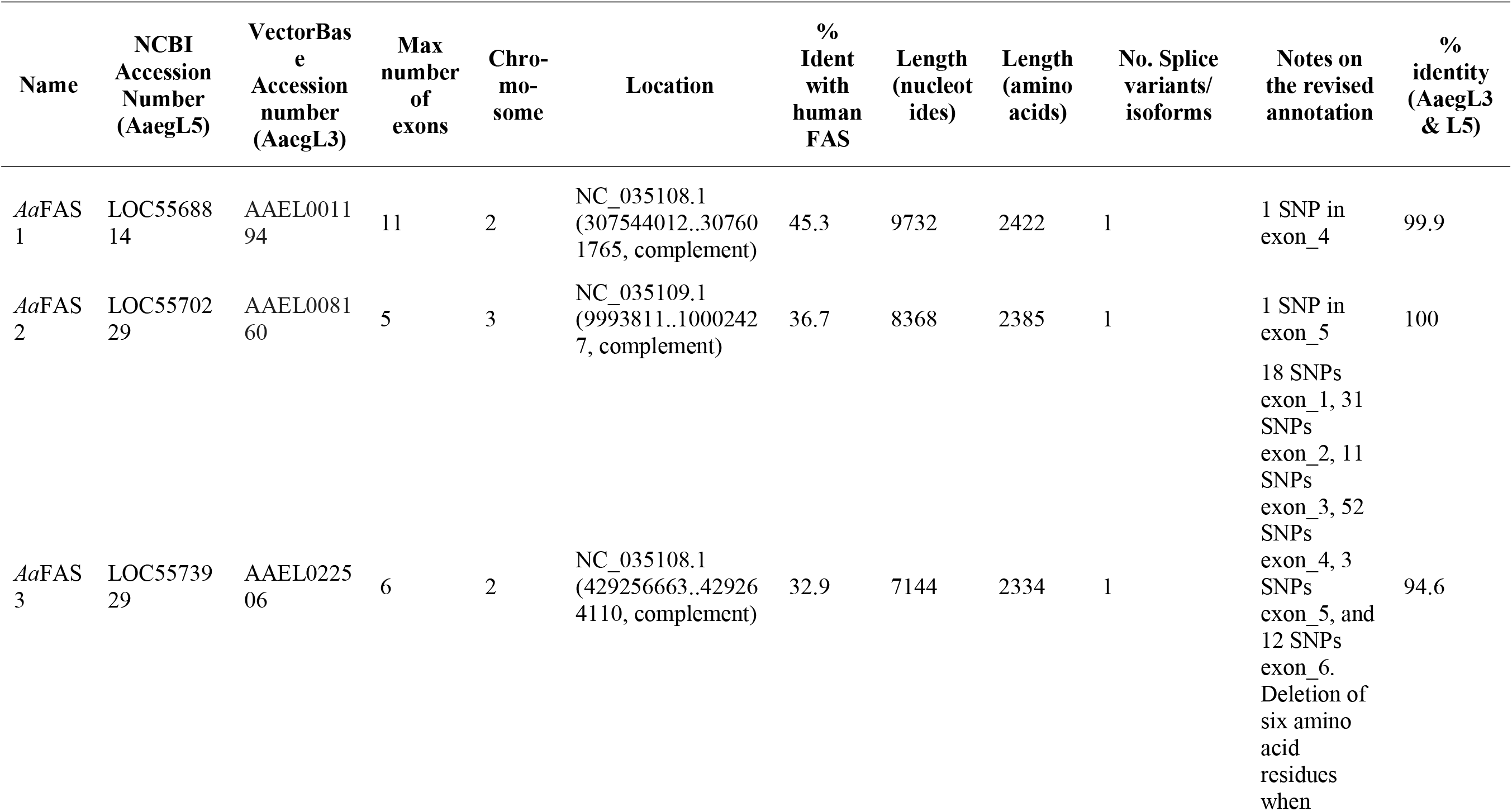
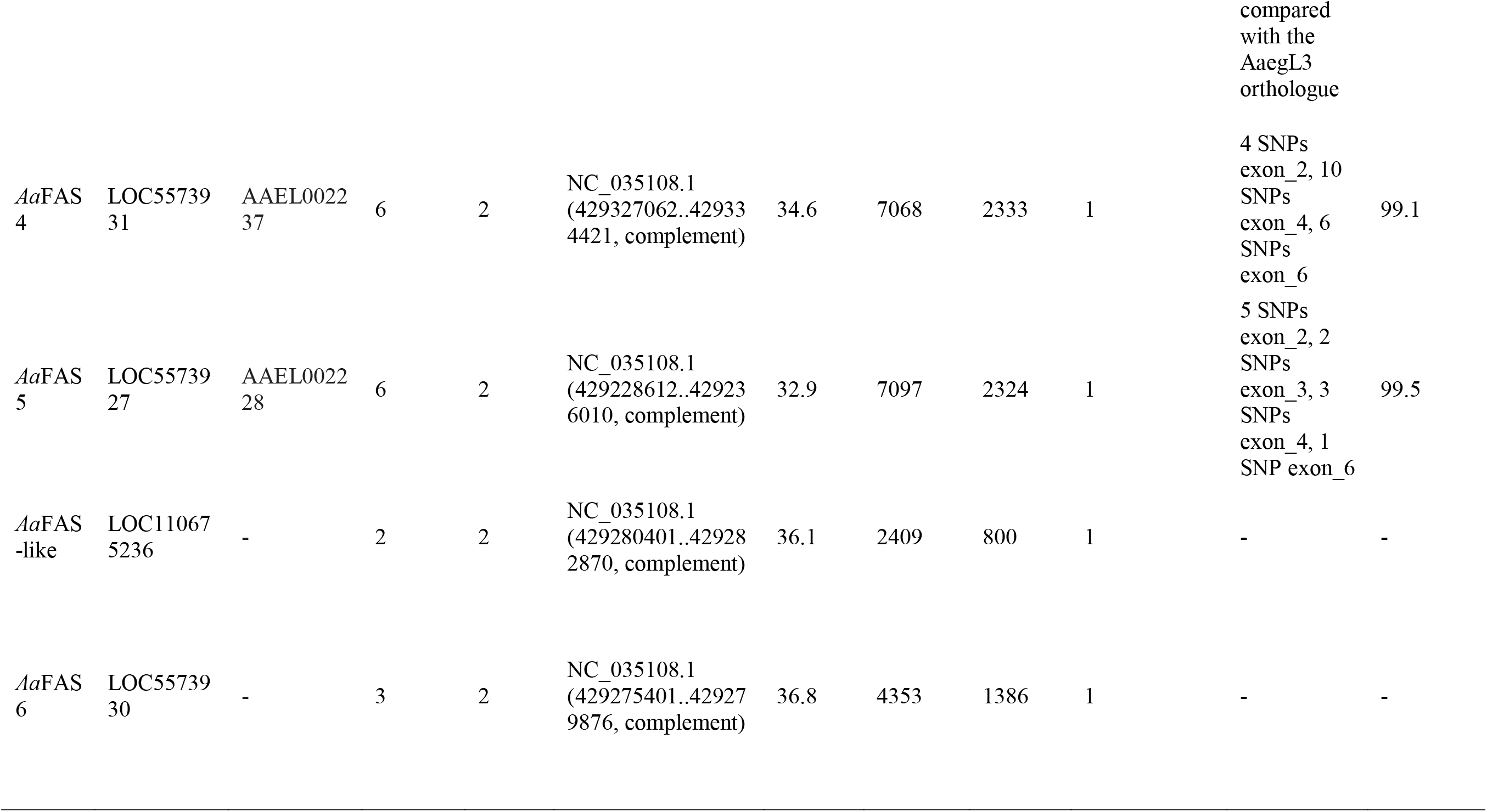
Summary of *Aa*FAS gene family predicted from the *Aedes aegypti* AaegL5 assembly. The AaegL5 annotation is shown in comparison to the AaegL3 gene models reported by Nene et al., 2007 [23].

The Bayesian inference supported *Aa*FAS1-5 as paralogues, and revealed highest percent amino acid similarity between *Aa*FAS1 and the *H. sapiens* FAS (human FAS) (Fig. 2). Notably, *Aa*FAS1 clustered in a clade comprising the *H. sapiens*, *Mus musculus, Apis mellifera* FAS, the *D. melanogaster* FAS1 and 2, and an uncharacterized *Anopheles gambie* FAS (Fig. 2). Similarly, *Aa*FAS2 clustered in a clade with another uncharacterized *Anopheles gambiae* FAS. In contrast, *Aa*FAS3, 4, 5, 6 and -like clustered at the most branched portion of the tree, suggesting a recent diversification event. Phylogenetic analyses and amino acid alignment supported *Aa*FAS1-5 as the counterparts of the AaegL3 genome assembly-derived gene models as follows: LOC5568814-AAEL001194; LOC5570229-AAEL008160; LOC5573929-AAEL022506; LOC5573931-AAEL002237 & LOC5573927-AAEL002228 (Fig. 2, Table 1). *Aa*FAS-like and *Aa*FAS6 (LOC110675236 & LOC5573930) were not identified in the AaegL3 assembly suggesting these models are unique to the AaegL5 assembly.

**Fig. 2.**
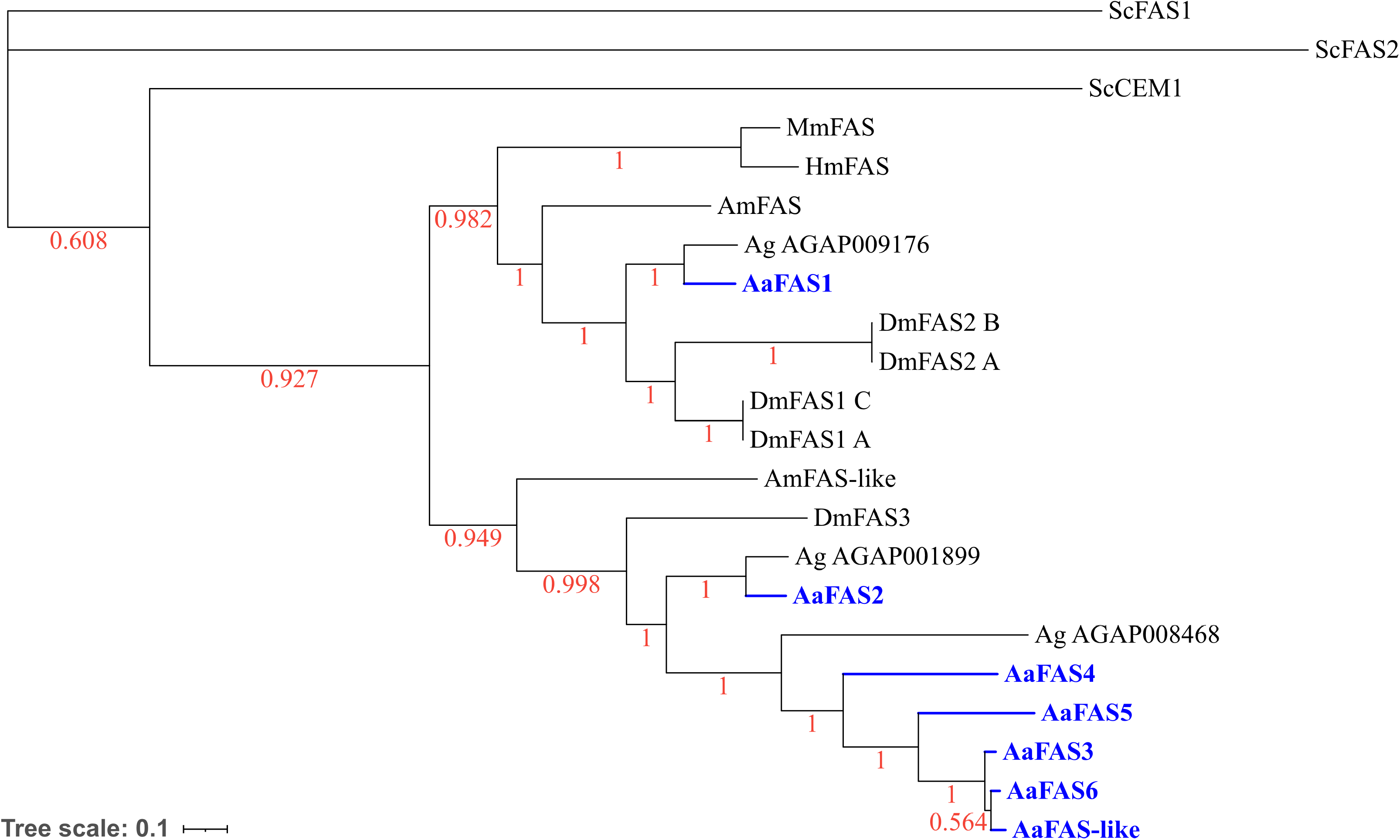
Phylogenetic analysis of *Aa*FAS. Bayesian phylogeny represented with an unrooted tree showing the main relationships between mosquito FAS genes and their counterparts in Ag: *Anopheles gambiae,* Dm: *Drosophila melanogaster,* Am: *Apis mellifera,* Mm: *Mus musculus*, Hm: *Homo sapiens* and Sc: *Saccharomyces cerevisiae*. The branches are supported by posterior probability values. The tree is drawn to scale: amino acid changes per site.

To investigate putative functional domains, *Aa*FAS sequences were aligned to the human FAS using Clustal Omega (27). Human FAS contains seven catalytic domains and three noncatalytic domains (1). Collectively, *Aa*FAS posssesed less than 50% amino acid identity to human FAS, and of the seven gene models, *Aa*FAS1 had the highest amino acid identity (45.3%) (Table 1 and Table S5). Alignment of FAS domains also showed modest sequence identity between human FAS and *Aa*FAS (23.03-63.56%) with greatest similarity for *Aa*FAS1 domains (Table S5). The linear organization of mammalian FAS domains annotated by Maier et. al., 2008, is shown in Fig. 3 (1). Conservation in linear organization of motifs associated with known functional domains identified using Pfam 31.0 software is shown in Fig. 3B. Dotted lines between Fig. 3A and B compare mammalian FAS domains (Fig. 3A) and *Aa*FAS domains (Fig. 3B). Pfam analysis did not show the presence of functional methyltransferase domains in *Aa*FAS (Fig. 3B) and protein sequence alignment using Clustal Omega showed deletion within pseudo-methyltransferase (ΨME) domains of *Aa*FAS compared to human FAS (16.20-23.03% identity; Table S5 and Fig. S2).

**Fig. 3.**
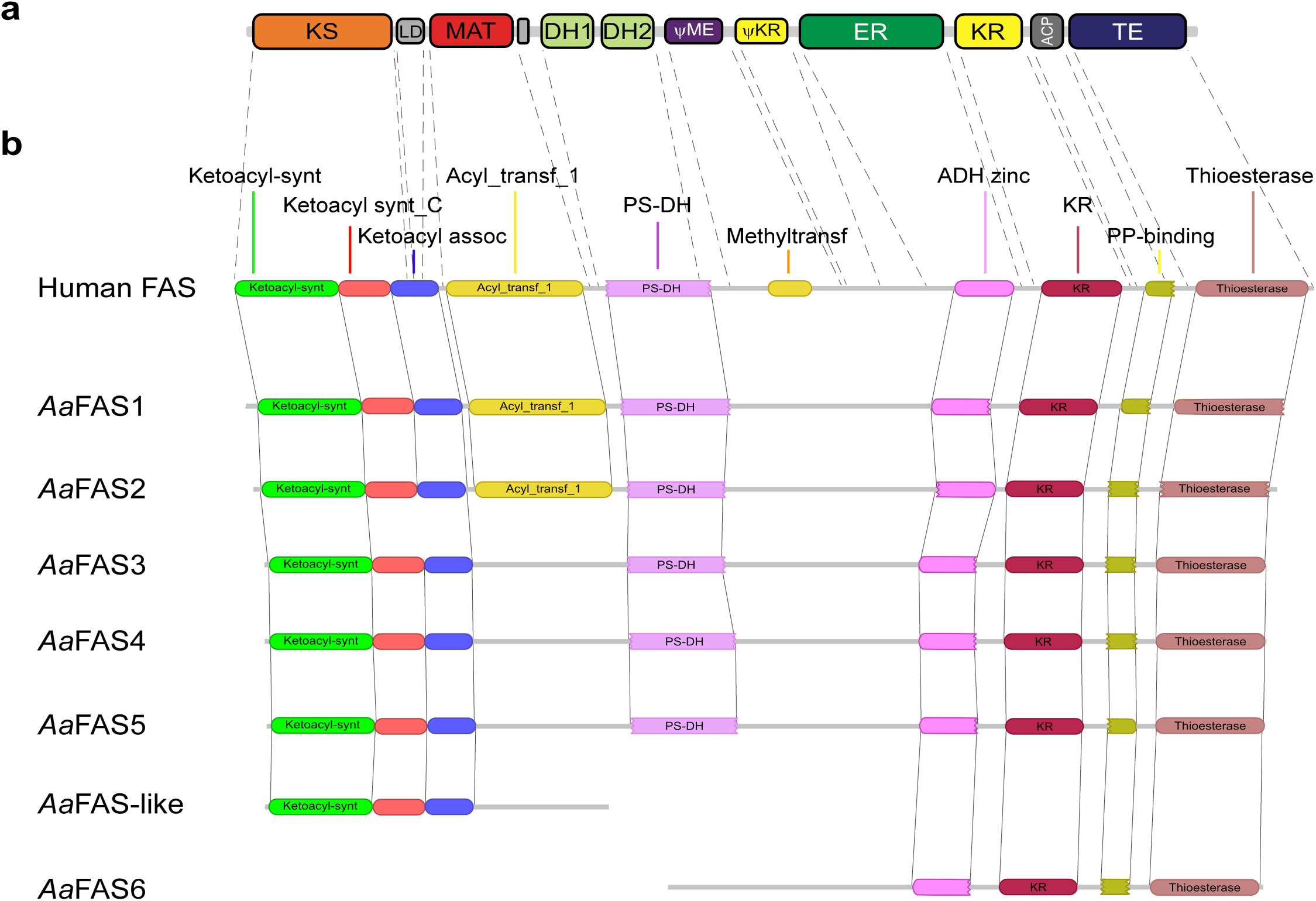
Linear organization of *Aedes aegypti* FAS genes showing functional domains. (**a**) Schematic shows linear organization of 7 catalytic and 3 noncatalytic domains of mammalian FAS annotated by Maier et. al., 2008 (1). Seven catalytic domains are shown in big squares and 3 non-catalytic domains are shown in smaller squares. Abbreviations: KS, β-ketoacyl synthase; LD, linker; MAT, malonyl-acetyl transferase; DH, dehydratase; ΨME, pseudo-methyltransferase; ΨKR, pseudo β-ketoacyl synthase; ER, enoyl reductase; KR, β-ketoacyl synthase; ACP; acyl carrier protein and TE, thioesterase. (**b**) Schematics show conserved domains or motifs of FAS genes and their organization annotated using Pfam 31.0 software. Abbreviations are as follows: ketoacyl_synt, β-ketoacyl synthase; ketoacyl_synt_C, β-ketoacyl-acyl carrier protein synthase; ketoacyl_assoc, ketoacyl-synthase C-terminal extension; acyl_transf_1, acyl transferase domain; PS-DH, polyketide synthase; methyltransf, methyltransferase domain; ADH-zinc, zinc binding dehydrogenase; KR, β-ketoreductase domain; PP-binding, phosphopantetheine attachment site; thioesterase, thioesterase domain.

The gene model of *Aa*FAS-like was 800 amino acids in length and contained all functional catalytic motifs, whereas *Aa*FAS6 was 1,386 amino acids in length, and lacked catalytic motifs but contained the conserved 3’ motif YKELRLRGY conserved in FAS (Fig. S1). *Aa*FAS-like contains ketoacyl synthase, ketoacyl synthase_C and ketoacyl-synthase C-terminal extension domains, the first 5’ domains of *Aa*FAS1-5 and human FAS (Fig. 3B), while *Aa*FAS6 contains ADH zinc, β-ketoreductase, PP binding and thioesterase domains, the last four domains located 3’ in *Aa*FAS1-5 and human FAS (Fig. 3B). In the AaegL5 assembly, *Aa*FAS-like and 6 are located on chromosome 2 at positions 429280401-429282870 and 429275401-429279876, respectively. It is possible that *Aa*FAS-like and -6 reflect an error in genome assembly, or a gene duplication. However, molecular data and the inability to detect transcripts associated with either locus, suggest (Fig. S3) that *Aa*FAS-like and 6 represent pseudogenes (Fig. 3B).

### FAS expression during *Ae. aegypti* development

Mosquitoes undergo four developmental stages: egg, larva, pupa and adult. RT-PCR was used to explore the hypothesis that expression patterns of *Aa*FAS genes vary among these stages. Five individual mosquitoes were collected for each of the 4^th^ larval instar, pupa and adult stages. Expression of each of the *Aa*FAS genes was normalized to the average expression of the β-actin gene (2^-ΔCt^) (Fig. 4).

**Fig. 4.**
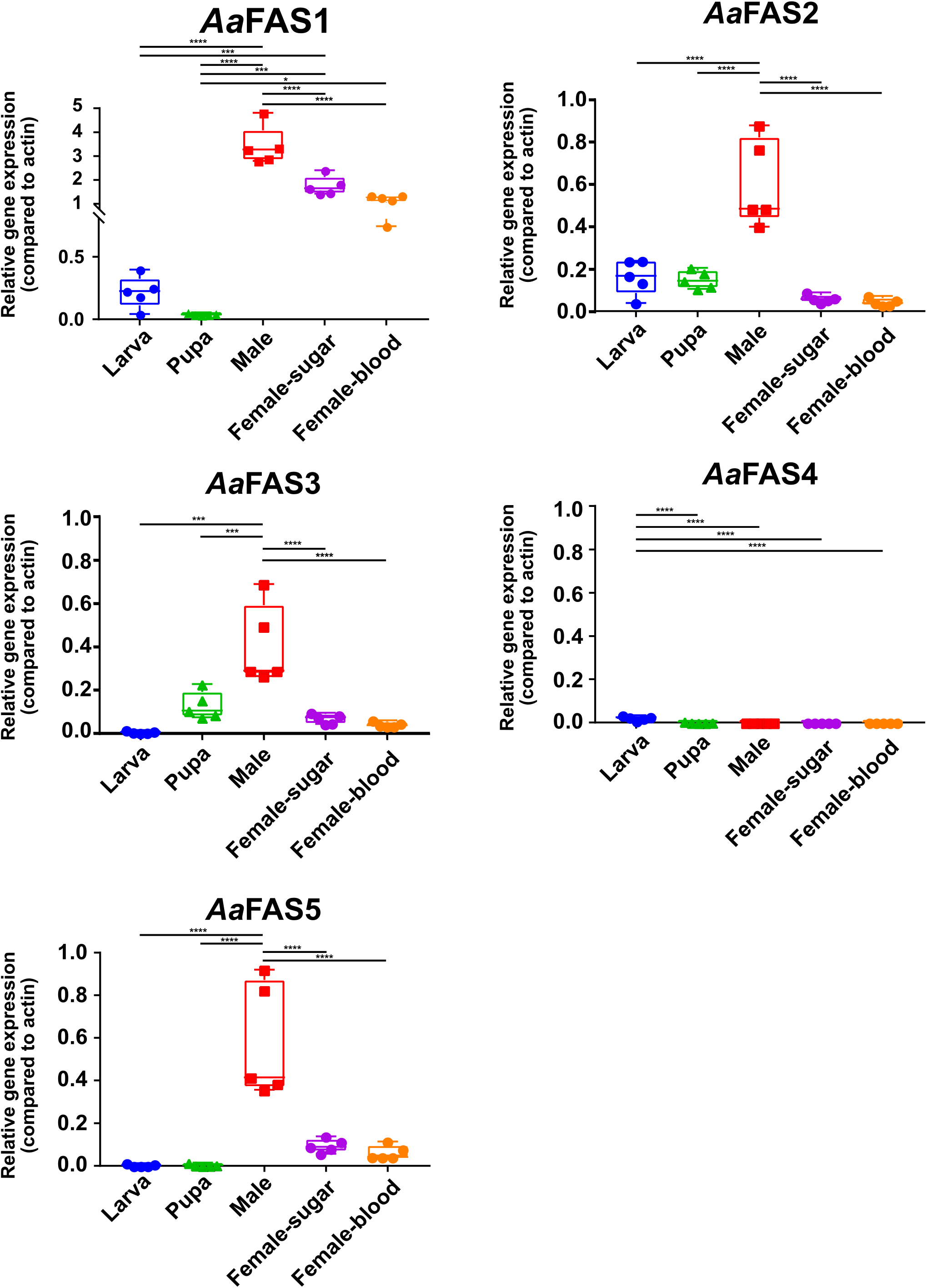
Expression of *Aa*FAS in mosquito developmental stages and sexes. RNA was prepared from 5 specimens of 4^th^ instar larvae, pupae, sugar-fed males, sugar-fed females, and females 3-days pbm (all adult mosquitoes were collected at the same day time; 8-10 days post eclosion). Samples were subjected to RT-qPCR to assess relative expression of *Aa*FAS1-5 and *Aa*FAS-like. RNA levels between samples were normalized to the β-actin gene using 2^-ΔCt^ method. The boxes show the 25^th^ and 75^th^ percentiles, the whiskers show the minimum and maximum values and the midline indicates the median of the relative gene expression value.

Relative expression analyses revealed negligible *Aa*FAS expression in larval and pupal stages, while the highest expression of all genes except *Aa*FAS4 were observed in adult males (Fig. 4). *Aa*FAS1 was the most predominant FAS expressed in adult mosquitoes. Differences in expression levels of any *Aa*FAS were not observed between sugar-fed and 3 days post blood-fed (coinciding with the first gonotrophic cycle) females. The study also revealed negligible *Aa*FAS4 expression in all developmental stages and sexes (Fig. 4).

### Impact of blood feeding on expression of *Aa*FAS1

The diet of the female *Ae. aegypti* typically involves both nectar and blood. The blood meal is rich in proteins and lipids; therefore, this diet may trigger lipolysis, instead of synthesis, to break down lipid molecules. We compared *Aa*FAS1 expression, the predominant *Aa*FAS in adult females, in sugar-fed females versus blood-fed females (feeding once or twice) (Fig. 5). Blood meals were provided only on specific days as shown in Fig. 5A, while mosquitoes from all groups were fed ad lib on 10% sugar diet throughout the experiment. Comparisons of *Aa*FAS1 gene expression from mosquito samples collected on the same day showed no differences among feeding conditions (Fig. 5B). However, when profiled as ratios (Fig. 5C), we observed a slight, but not significant, reduction of *Aa*FAS1 expression in females given a single blood-meal as compared to sugar-fed females on days 1, 3 and 4 post-blood meal (pbm) (Fig. 5B: F vs. B, G vs. C and H vs. D). This data suggests that diet may only play a minor role, if any, in the expression of *Aa*FAS1 gene.

**Fig. 5.**
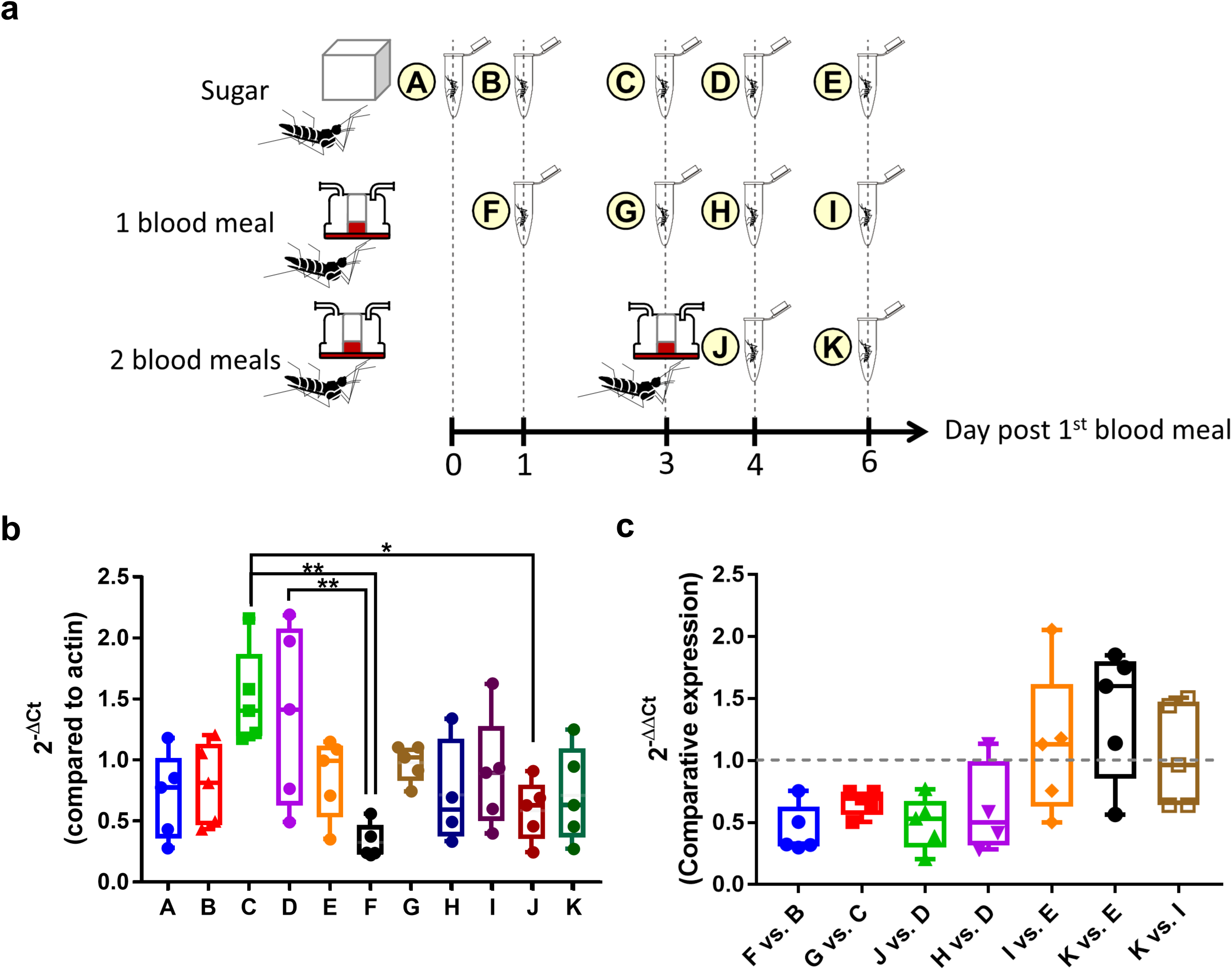
Expression of *Aa*FAS1 in sugar-fed and blood-fed mosquitoes. (**a**) Schematic of the experimental design. Mosquitoes were reared in 3 different conditions: sugar-fed only, one blood meal, which were fed on day zero, and two blood meals, which were again fed on day 3. Mosquitoes in all conditions were also allowed access to additional sugar and water at all time throughout the experiment. Five mosquitoes from each condition were collected on different days post 1^st^ blood meal feeding designated in letters A-K. (**b**) *Aa*FAS1 expression profile (2^-ΔCt^; normalized to actin) of each sampling group were shown. Statistical analysis was performed by one-way ANOVA followed by Tukey’s multiple comparison tests. *, p<0.05 and **, p< 0.01. (**c**) Expression of *Aa*FAS1 was measured by the comparative expression (2^-ΔΔCt^) method. The samples used for comparisons are shown on the X-axis. Boxes show the 25^th^ and 75^th^ percentiles, whiskers show the minimum and maximum values, and the midline shows the median of the relative gene expression value.

### Transient knockdown of *Aa*FAS1 gene causing upregulation of other *Aa*FAS genes

We hypothesized that the redundancy of *Aa*FAS genes may serve as a backup system for the mosquitoes. To test this hypothesis, we employed *Aa*FAS1 loss-of-function studies to investigate the possibility of compensation by other *Aa*FAS genes. Female mosquitoes were IT injected with dsRNA derived from *Aa*FAS1 or GFP (KD control). On day 2 post-dsRNA injection, five mosquitoes were collected for assessment of *Aa*FAS expression (Fig. 6). We observed an approximate 40% reduction in *Aa*FAS1 expression (∼39.3±13.9%) in *Aa*FAS1-KD mosquitoes compared to the GFP-KD control (Fig. 6A). In *Aa*FAS1-KD mosquitoes, expression levels of *Aa*FAS2, 3 and 5 and were 191.7±38.6%, 161.4±21.8%, and 191.1±38.9%, respectively, in comparison to their levels in GFP-KD control, indicating possible compensation for the loss of *Aa*FAS1 transcript. Conversely, the expression of *Aa*FAS4 was 87.71±74.0% compared to *Aa*FAS4 expression in GFP-KD control mosquitoes. To determine whether the upregulation observed in *Aa*FAS2, 3, and 5 could possibly compensate for the loss of *Aa*FAS1 in the *Aa*FAS1-KD mosquitoes, we normalized the level of *Aa*FAS genes to β-actin. We observed modest expression of *Aa*FAS transcripts (5.6±1.44% for *Aa*FAS2, 4.6±0.00% for *Aa*FAS3, and 7.07±0.62% for *Aa*FAS5 compared to β-actin), while these upregulation still did not match the remnent of *Aa*FAS1 expression after the KD effect (36.1±11.6%). This data suggest that other *Aa*FASs may not be able to serve as a backup system for *Aa*FAS1, at least in adult female mosquitoes under transient KD condition.

**Fig. 6.**
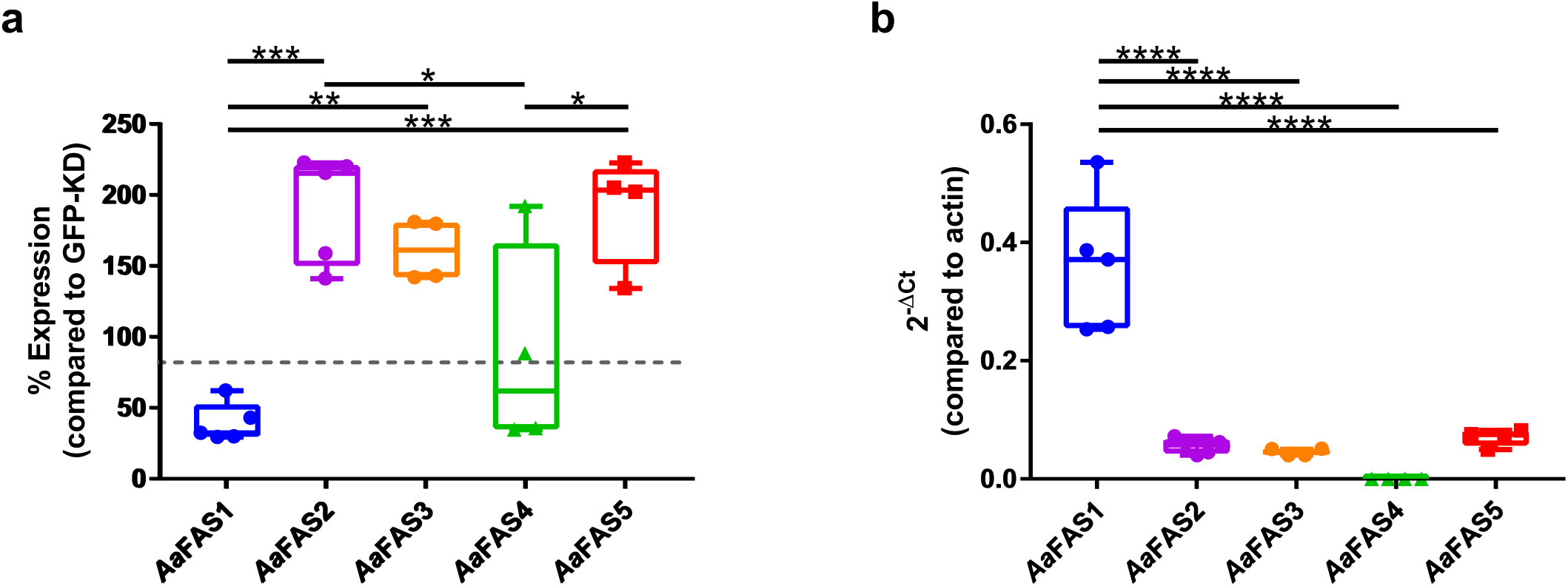
Comparive analyses of *Aa*FAS expression in *Aa*FAS1-KD mosquito. Three-day-old adult female mosquitoes were intrathoracically injected with dsRNA derived from *Aa*FAS1 or GFP mRNA sequence (an irrelevant dsRNA control). Mosquitoes were reared on 10% sugar diet for 2 days post-injection. Three pools of 5 mosquitoes were collected for *Aa*FAS gene expression measurements. (**a**) Percent relative expression of *Aa*FAS genes in *Aa*FAS1-KD mosquitoes was compared to GFP-KD control mosquitoes using the comparative Ct (2^-ΔΔCt^) method. (**b**) Gene expression profiles of *Aa*FAS were measured as normalized to β-actin gene expression (2^-ΔCt^). Boxes show the 25^th^ and 75^th^ percentiles, whiskers show the minimum and maximum values and midline shows median of the relative gene expression value. Statistical analysis was performed by one-way ANOVA followed by Tukey’s multiple comparison tests. **, p< 0.01, ***, p<0.005 and ****, p<0.001.

### Effect of RNAi-induced *Aa*FAS1 knockdown on DENV2 replication in *Ae. aegypti* cells

Studies in cell culture have shown that FAS activity is required for flavivirus genome replication (13, 14, 37). Biochemical inhibition of FAS activity reduced DENV2 replication in both human and mosquito C6/36 cells (13, 15, 16). The lack of functional RNAi machinery in C6/36 cells hindered the use of transient KD strategy in mosquito cells. However, *Ae*. aegypti cells, Aag2, have functional RNAi machinery; therefore, we can investigate the role of *Aa*FAS1, the most abundant transcript in female mosquitoes, in DENV2 replication using dsRNA transient KD in these cells (38). At 48 hours post-*Aa*FAS1-KD (time zero of DENV2 infection), the expression level of *Aa*FAS1 in Aag2 cells was 5.15 ± 6.33 % as compared to *Aa*FAS1 expression in GFP-KD control cells (Fig. 7A). At 24 hours post DENV2 infection (72 hours post-KD), we observed significant reduction (p<0.001) in DENV2 RNA replication in *Aa*FAS1-KD cells as compared to the GFP-KD controls, comparable to replication in DENV2-KD (KD positive control) (Fig. 7B). KD was not associated with detrimental effects to the cells (Fig. 7C), suggesting that *Aa*FAS1 is required for DENV2 replication in mosquito cells.

**Fig. 7.**
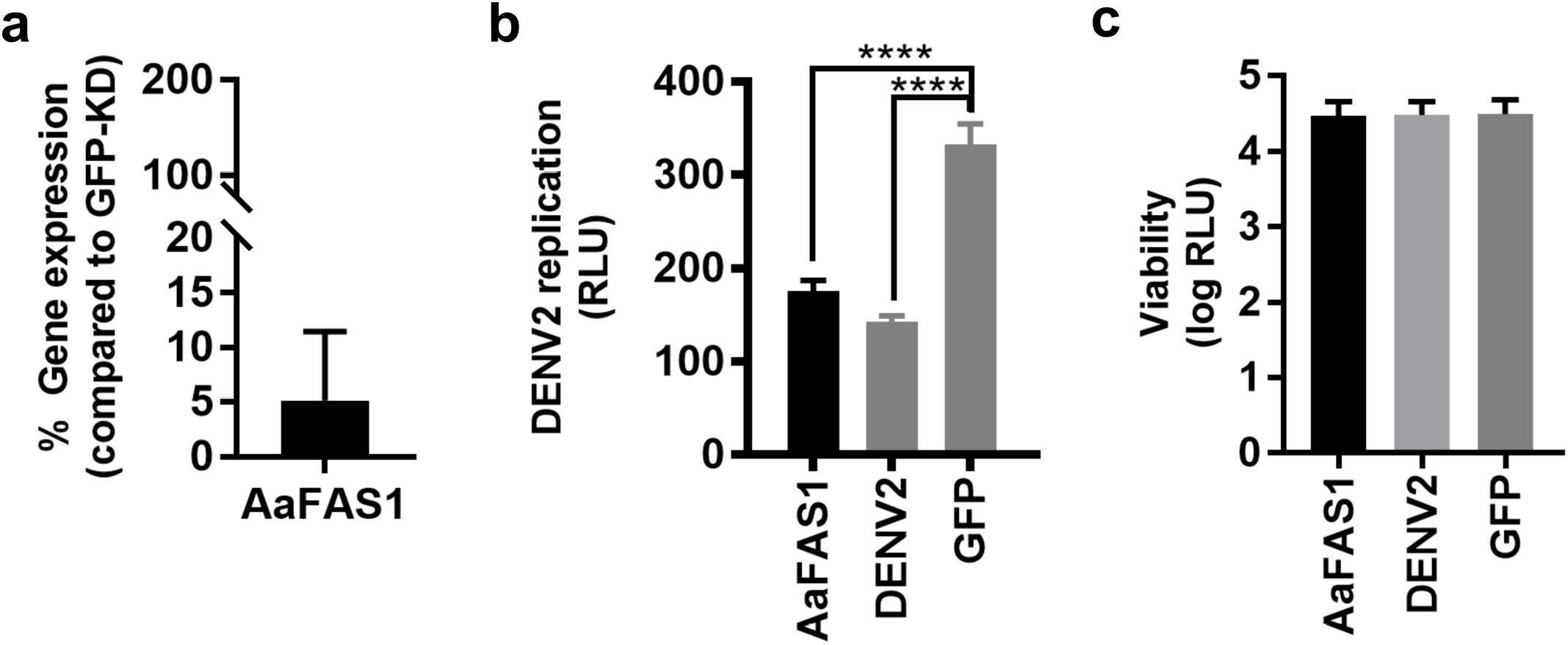
Assessment of RNAi-induced *Aa*FAS1 knockdown on DENV2 replication in Aag2 cells. (**a**) Percent *Aa*FAS1 expression in *Aa*FAS1-KD Aag2 cells compared to the *Aa*FAS1 expression in GFP-KD cells. Expression was measured at two days post dsRNA transfection. (**b**) RNA replication of luciferase-expressing DENV2 in Aag2 cells treated with dsRNA derived from *Aa*FAS1, DENV2 or GFP. Cells were transfected with dsRNA for two days prior to infection with luciferase-tagged DENV2. At 24 hpi, cells were lysed and were assayed for luciferase activity (RLU). One-way ANOVA followed by Tukey’s multiple comparison test were applied for statistical analysis. ****, p < 0.001. (**c**) Viability of Aag2 cells treated with dsRNA derived from *Aa*FAS1, DENV2 (positive DENV2 KD control) and GFP (negative DENV2 KD control) genes assessed by resazurin assay.

### Transient inhibition of *Aa*FAS1 reduced DENV2 infection in the midgut of *Ae. aegypti*

To investigate the role of *Aa*FAS1 in DENV2 replication in vivo, mosquitoes were IT injected with dsRNA derived from *Aa*FAS1 or GFP genes, and subsequently exposed to DENV2 infectious blood meal two-days post-injection (Fig. 8). On days 0, 3 and 7 pbm (corresponding to 2, 5 and 9 days post-dsRNA injection), whole mosquitoes were collected and analyzed for *Aa*FAS1 gene expression (Fig. 8A). On the day of DENV2 infection by blood meal (2 days post-dsRNA injection), the level of *Aa*FAS1 expression was downregulated by 53.73 ± 27.13% relative to GFP-KD group. On day 3 pbm, *Aa*FAS1 expression recovered to 119.82 ± 49.43 % and was comparable to the *Aa*FAS1 expression level in the GFP-KD control. On day 7 pbm, *Aa*FAS1 was upregulated to 191.69 ± 50.17 %, suggesting a possible over-compensation post KD effect (Fig. 8A).

**Fig. 8.**
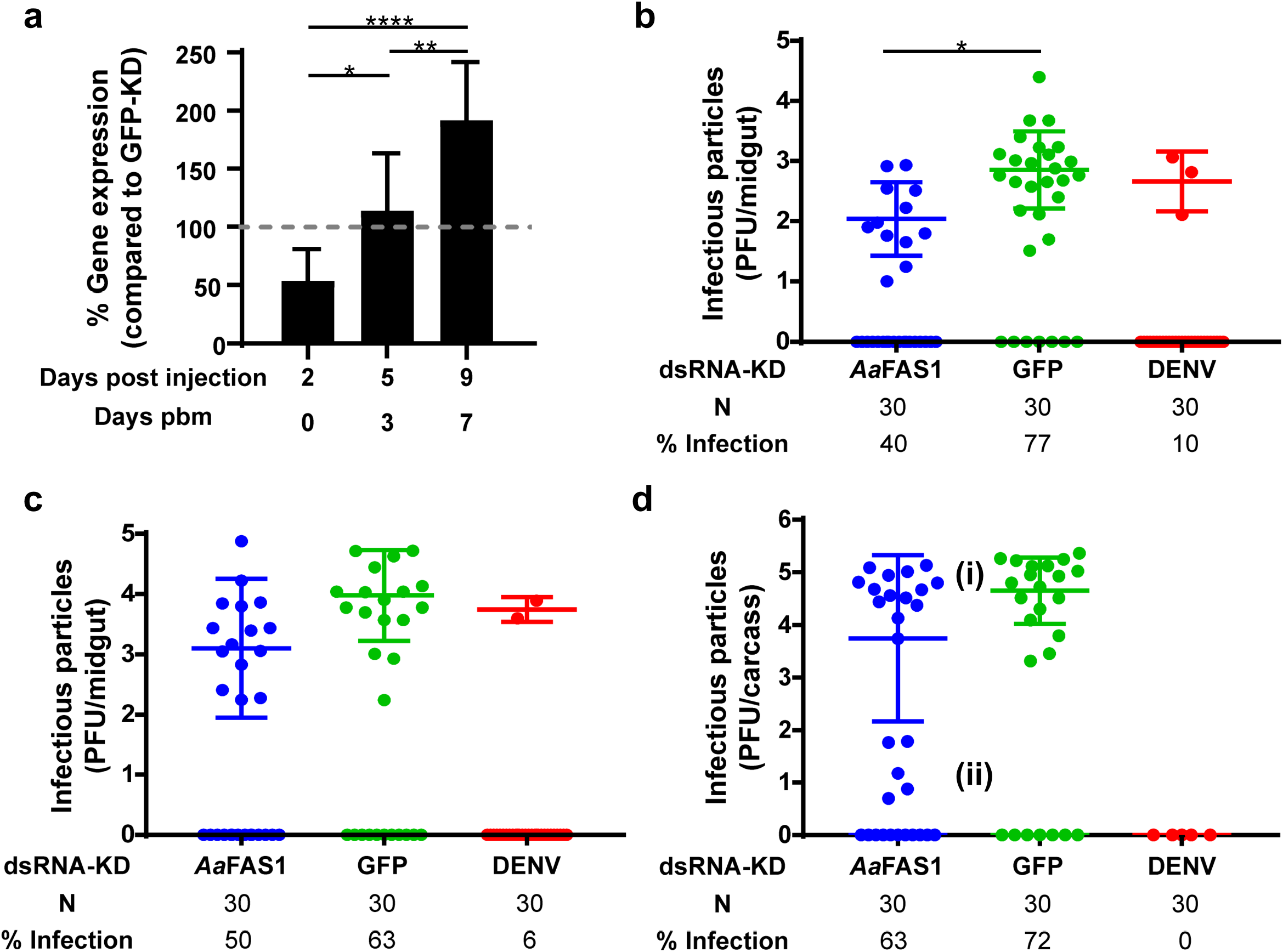
Transient KD of *Aa*FAS1 expression by dsRNA temporarily reduced DENV2 infection in midguts. (**a**) Percent *Aa*FAS1 expression in FAS-KD mosquitoes. Mosquitoes were IT injected with ∼400 ng of dsRNA derived from *Aa*FAS1, DENV2 or GFP two days prior to blood meal infection. On days 0, 3 and 7 pbm (days 2, 5 and 9 days post IT injection), 3 pools of 5 mosquitoes were collected and analyzed for *Aa*FAS1 expression compared to *Aa*FAS1 expression in GFP-KD mosquitoes. (**b-d**) Mosquitoes were IT injected with dsRNAs and infected with DENV2 by infectious blood meal at 2 days post injection. Plaque assay was performed on midguts dissected on (**b**) day 3 and (**c**) day 7 and (**d**) carcasses (whole body without midgut) collected on day 14 pbm. (i) and (ii) indicate the separation of DENV2 titers in the carcass that were produced from the *Aa*FAS1-KD mosquitoes. Mean virus titer (infectious particles) was calculated for infected samples only. One-way ANOVA followed by Dunn’s tests were applied to test the differences in virus titer among samples, *, *p* < 0.05. The odds ratio was applied to test the differences in percent infection. The significant reduction of percent infection in *Aa*FAS1-KD or DENV2-KD compared to GFP-KD are highlighted in bold.

Investigation of DENV2-fed mosquitoes showed that, at day 3 pbm, we observed significant reduction in percent of DENV2 infected midguts compared to the GFP-KD control (Fig. 8B). The odds ratio for *Aa*FAS1 in *Aa*FAS1-KD mosquitoes compared to GFP-KD mosquitoes on day 3 pbm was 0.20 (95% confidence interval (CI): 0.06 - 0.60) and for DENV2 compared to GFP-KD was 0.03 (CI: 0.01 – 0.13) suggesting fewer mosquitoes were infected with DENV2 in the *Aa*FAS1-KD group compared to the GFP-KD group on day 3 pbm. However, no differences in percent infection were observed between *Aa*FAS1-KD and control mosquitoes on days 7 and 14 pbm. Infectious particles produced from midguts (virus titer) from *Aa*FAS1-KD and DENV2-KD groups were significantly different from GFP-KD group when the uninfected samples were included in the analysis using the nonparametric Kruskal-Wallis test followed by Dunn’s test, with p-values adjusted with the Bonferroni method (*p* = 0.0002 and *p* < 0.0001, respectively). However, if the titers of uninfected midguts were excluded, differences in virus titer among different dsRNA treatments were not detected (tested by one-way ANOVA followed by Dunn’s test; virus titer *Aa*FAS1-KD: 2.40×10^2^, GFP-KD: 2.18×10^3^, and DENV2- KD: 6.43×10^2^ plaque forming unit (PFU/midgut).

The inhibitory effect of *Aa*FAS1-KD on DENV2 infection did not persist in the midgut beyond day 3 pbm. The titer and percent infection in the *Aa*FAS1-KD mosquitoes were comparable to the GFP-KD mosquitoes on day 7 pbm (*Aa*FAS1-KD: 2.70×10^3^, GFP-KD: 2.95×10^4^ and DENV2-KD: 5.88×10^3^ PFU/midgut; Fig. 8C). No differences in viral titer and percent infection were observed in midgut on day 7 pbm. To investigate whether transient *Aa*FAS1-KD could disrupt virus dissemination, mosquito carcasses (whole body without midgut) were tested on day 14 pbm for virus infection (*Aa*FAS1-KD: 6.57×10^4^, GFP-KD: 8.47×10^4^ and DENV2-KD: 0.00 PFU/carcass; Fig. 8D). Although we observed no statistical differences in mean titer in *Aa*FAS1-KD as compared to GFP-KD control samples, two distinct populations of mosquitoes with viral titers in *Aa*FAS1-KD carcasses were observed (Fig. 8D); some with viral titers comparable to GFP-KD control (5.77×10^4^ PFU/carcass; i) and some with distinctively lower titers (2.95×10^1^ PFU/carcass, ii). This observation suggsts that transient KD of *Aa*FAS1 had a prolonged effect that can impact dissemination of DENV2 in mosquitoes.

## Discussion

Lipids are essential for a variety of physiological processes in mosquitoes (3, 10, 12, 39, 40). Mosquitoes not only acquire lipids from maternal (i.e. deposition to eggs) and dietary sources, but they also have the ability to synthesize lipids de novo. In this study, we characterized the expression of the *Aa*FAS gene family, the key enzyme in the *de novo* lipid biosynthesis pathway. Additionally, we characterized the response of *Aa*FAS1 expression during infection of mosquitoes and mosquito cell lines with DENV2 to investigate the potential role of this enzyme in supporting virus replication in the mosquito vector.

Seven putative *Aa*FAS genes (*Aa*FAS1-6 and *Aa*FAS-like), were identified in the AaegL5 assembly based on amino acid similarity to FAS from vertebrates, invertebrates and yeast. Amino acid sequence alignments revealed low amino acid similarity (<50%) of *Aa*FAS compared to human FAS (Table 1).

The *Aa*FAS1-5 possess seven catalytic and two of the three noncatalytic domains identified in the human FAS annotated using Pfam (Fig. 3) (1, 29). All *Aa*FAS lack the ΨME domain observed in *Drosophila melanogaster* (fruit fly), *Bombyx mori* (silkworm), *Apis mellifera* (honey bee), *Culex pipiens*, and *Anopheles gambiae* (1). Similarly, mammalian FAS also loses the conservation of the motif involved in S-adenosyl-methionine (SAM)–dependent methyltransferases from the ΨME, resulting in an absencce of methyltransferase activity, while it is highly conserved in bacteria and fungi (1). An absence of ΨME domain in insect FAS may reflect the unnecessity of this domain.

Gene duplication is a hallmark of many mosquito gene families and has been proposed as a source of new evolutionary features (23, 36, 41). Retention of duplicated genes may be indicative of positive/neutral selection and loci associated with a fitness advantage for the mosquito (42). We performed molecular and prelimnary functional characterization of the *Aa*FAS gene family. We detected transcripts for five of the seven *Aa*FAS genes. We were unable to detect transcripts for either *Aa*FAS6 and *Aa*FAS-like, and conclude that these gene models likely represent pseudogenes or may reflect an issue in the assembly.

Since mosquitoes undergo four distinct developmental stages in their life and these stages possess very distinct habitats and food sources, different *Aa*FAS genes may play roles supporting the unique requirements for FAS in these different life stages. Transcription profiles of *Aa*FAS1-5 revealed expression pattern. Notably, we observed low expression levels for all *Aa*FAS in larval and pupal stages, suggesting that these genes may not be constitutively active across the mosquito life-cycle. We speculate that maternal lipids deposited in eggs during oogenesis (these comprise about 35% of dry egg weight (7)) and larvae diets may serve to support the metabolic needs during these stages (8, 9, 43). Thus, they may have minimal requirement for de novo fatty acid biosynthesis.

We observed high levels of expression of all *Aa*FAS, except *Aa*FAS4, in males and to a lesser extent in females. Male *Ae. aegypti* do not blood feed, but solely obtain their diet from plant nectar, honeydew and fruits (44). Since these diets are high in carbohydrate but low in lipid content, high expression of *Aa*FAS in male mosquitoes may reflect a dependency on *Aa*FAS for *de novo* synthesis of lipids.

Expression analyses also revealed *Aa*FAS1 as the dominant *Aa*FAS transcript in both male and female mosquitoes (Fig. 4). The *Aa*FAS1 had the highst amino acid similarity among the *Aa*FAS to the human and mouse FAS genes and is the putative paralogue of these genes. Upon *Aa*FAS1-KD in female mosquitoes, we observed an ∼ 2-fold increase in other *Aa*FAS transcripts indicating the attempt to compensate for the loss of *Aa*FAS1 expression (Fig. 6A). Although we concluded that they fail to compensation for the loss of *Aa*FAS1 since the expression levels of these *Aa*FAS transcripts are still lower than the remaining *Aa*FAS1 expression post-KD, improving the KD efficiency or extending the period of KD of *Aa*FAS1 (such as using CRISPR-CAS9 knockout) may provide further insights into the redundancy of these *Aa*FAS genes.

Since previous studies have demonstrated the importance of FAS activity in flavivirus replication in both human and mosquito cells (13, 15, 16), we wanted to investigate if *Aa*FAS1, also played an important role in DENV2 infection in the mosquito vector. Indeed, KD of *Aa*FAS1 showed significant inhibition of DENV2 infection in midguts of *Aa*FAS1-KD mosquitoes; however, this effect was only observed on day 3 pbm. This phenomenon might be caused by the transient KD of *Aa*FAS expression. Alternative studies with longer suppression of *Aa*FAS1 expression would be required to demonstrate the prominent impact on infection and for further transmission studies.

Nonetheless, we found an upregulation of *Aa*FAS1 expression (200% increase compared to the *Aa*FAS1 levels in the GFP-KD control) on day 9 post-KD. With relevance for strategies aimed at suppression of host factors for disrupting pathogen transmission,further studies are needed to better understand the biological impact of this “rebound” effect.

Additionally, we observed a separation of the virus titers into two groups: high (Fig. 8D, i) and low (Fig. 8D, ii), titers in the carcass of the *Aa*FAS1-KD mosquitoes. Ye et. al. 2015, have shown that when mosquitoes were IT injected with DENV at 10^6^ PFU, they expectorated DENV into the saliva at about 10^2^ PFU, which is about 100-fold less than mosquitoes that were IT injected with DENV at 10^7^ PFU (approximate DENV titer in saliva = 10^4^ PFU) (45). This result suggests that viral titer of DENV in saliva may be dependent on the titer in the body (disseminated titers). In our observations, it is possible that mosquitoes with low body titers (group ii) would inefficienty transmit the virus and the KD of *Aa*FAS1 may result in a reduction of transmission potential. This study suggests that biological relevance of low viral titres in carcasses and its impact on transmission dynamics is worthy of further investigation.

## Conclusions

Here we pesent expression analyses of the *Aa*FAS gene family and a focused study of the *Aa*FAS1 in *Ae. aegypti*. We annotated seven *Aa*FAS genes from the AaegL5 genome assembly, and present evidence to support function of five genes. Expression data revealed complexities of *Aa*FAS expression between stages and sexes, and suggest that *Aa*FAS1 is the dominant transcript in both male and female adult mossquitoes. Sequence homology suggested conservation between mammalian FAS and *Aa*FAS1, and the presence of multiple catalytic domains supports *Aa*FAS1 as a key enzyme in *de novo* lipid biosynthesis. In addition, *Aa*FAS1 was found to facilitate DENV2 replication in both cell culture and in *Ae. aegypti*. In the latter case, it demonstrated the potential to affect vector competency for virus transmission.

## Supporting information

Additional file 1_Chotiwan et al.docx

Additional file 2_Chotiwan et al.docx

Figure S1 _Chotiwan et al

Figure S2_Chotiwan et al

Figure S3 _Chotiwan et al

## Supplementary information

**Additional file 1: Figure S1.** Alignment of the conserved YKELRLRGY motif. **Figure S2.** Amino acid alignment of the pseudo-methyltransferase (ΨME) domain of *H. sapiens* and *Ae. aegypti* FAS. **Figure S3.** RT-PCR assays designed to detect mRNA products of *Aa*FAS-like and *Aa*FAS6. **Table S1.** List of vertebrate, invertebrate and yeast FAS gene models employed in the present study. **Table S2.** Primers for generation of dsRNA for knock-down studies. **Table S3.** Primers for *Aa*FAS expression analyses. **Table S4.** Primers for RT-PCR assay detecting mRNA products *Aa*FAS-like and *Aa*FAS6. **Table S5.** Amino acid similarity of FAS domains between *H. sapiens* and *Ae. aegypti*.

Additional file 2: File S1. mRNA and amino acid sequences of 7 *Aa*FAS genes.

## Abbreviations

ACC: acetyl-CoA carboxylase
Ae.aegypti: Aedes aegypti
AaFAS: Aedes aegypti fatty acid synthase
Bp: base pair
cDNA: complementary deoxyribonucleic acid
DENV2: dengue virus serotype 2
dpi: days post-infection
dsRNA: long double-stranded ribonucleic acid
FAS: fatty acid synthase
FBS: fetal bovine serum
GFP: green fluorescent protein
IT: intrathoracic
KD: knockdown
MEM: Minimum Essential Media
PBS: phosphate-buffered saline
mRNA: messenger ribonucleic acid
PCR: polymerase chain reaction quantitative polymerase chain reaction
pbm: post-blood meal
PFU: plaque forming unit
ΨME: pseudo-methyltransferase
RNA: ribonucleic acid
RNAi: interference ribonucleic acid
RT: reverse transcription
RT-PCR: reverse transcription polymerase chain reaction
SNP: single nucleotide polymorphism

## Declarations

### Acknowledgements

The authors thank Kenneth E. Olson and Irma Sanchez-Vargas for providing mosquito eggs. We also thank Rebecca Gullberg, Laura St. Clair, Jeffrey M. Grabowski and Richard J. Kuhn for advice on experimental design, data analysis and critical evaluation of the manuscript and Tach Costello for clerical and administrative support. Graphical Abstract was created using BioRender.

### Funding

This work was funded by R01AI151166 NIH-NIAID and the Boettcher Foundation Early Career Investigator Awards to RP. GR was funded by the R01AI151166 NIH-NIAID. CBS was supported by Purdue internal monies.

Supplement.

### Availability of Data and Materials

All data and materials were presented in the manuscript and supplementary information.

### Authors’ contribution

Author Contributions: NC, CBS, GR and EL carried out the experiments. NC, CBS, GR, EL, CAH and RP wrote the manuscript.

### Ethics approval and consent to participate

Not applicable

### Consent for publication

All authors consent for publication

### Author details

Not applicable

