## Additional file 1_Chotiwan et al.docx for "Expression of fatty acid synthase genes and their role in development and arboviral infection of *Aedes aegypti*"

**Supplemental Figure**

**Fig. S1. Alignment of the conserved YKELRLRGY motif.** Protein sequences of FAS from *Sacharomyces cereviceae, Homo sapiens, Mus musculus, Drosophila melanogaster, Apis mellifera, Anopheles gambiae* and *Ae. aegypti* were aligned using CLUSTAL OMEGA and curated in Jalview. Conserved residues are highlighted in blue and the consensus sequence is shown at the bottom. Size of the consensus residues correlates with its prevalence among organisms.

**Fig. S2. Amino acid alignment of the pseudo-methyltransferase (ΨME) domain of *H. sapiens* and *Ae. aegypti* FAS**. The amino acid sequence alignment was performed using Clustal Omega and displayed in mView. The alignment of human FAS, ΨME domain of human FAS and *Aa*FAS genes. The alignment revealed incomplete domains ΨME in *Aa*FAS genes, suggesting an issue with loss of function in mosquito genes (shown in yellow).

**Fig. S3.** **RT-PCR assays designed to detect mRNA products** **of *Aa*FAS-like and *Aa*FAS6.** (**a**) A schematic showing designed primers to amplify AaFAS1 (primer A; positive amplification control), combined *Aa*FAS-like and 6 (primerB; to determine if *Aa*FAS-like and 6 are the same gene but split during genome assembly), *Aa*FAS6 (primer C) and *Aa*FAS-like (primer D). (**b**) DNA electrophoresis showing cDNA generated from mosquito total RNA. Lane M; 1kbp marker (Generuler 1kbp DNA ladder, SM1203, Thermo Scientific), lane 1; non-template control (NTC) of primer set A, lane 2-4; primer set A, indicating amplification of *Aa*FAS1 mRNA (expected product 1976 bp), lane 5; NTC of primer set B, lane 6-9; primer set B, no mRNA product. (**c**) DNA electrophoresis of cDNA generated from DNase treated mosquito total RNA. Lane M; 1kbp marker, lane 1; non-template control (NTC) of primer set A, lane 2-5; primer set A, indicating amplification of *Aa*FAS1 mRNA, lane 6; NTC of primer set B, lane 7-9; primer set B, lane 10; NTC primer set C, lane 11-13 ; primer set C, no mRNA product, lane 14; NTC primer set D, lane 15-17 ; primer set C, no mRNA product. Results indicated that we cannot detect mRNA of *Aa*FAS-like, *Aa*FAS6 and combined *Aa*FAS-like and 6. Therefore, they represent pseudogene.

**Supplemental Tables**

**Table S1. List of vertebrate, invertebrate and yeast FAS gene models employed in the present study.**

| Organism | Gene name | NCBI Accession Number |
| --- | --- | --- |
| *Homo sapiens* | HmFAS | NP_004095.4 |
| *Mus musculus* | MmFAS | NP_032014.3 |
| *Drosophila melanogaster* | DmFAS1_A | NP_608748.1 |
|  | DmFAS1_C | NP_001137778.2 |
|  | DmFAS2_A | NP_647613.1 |
|  | DmFAS2_B | NP_001259986.1 |
|  | DmFAS3 | NP_001015405.3 |
| *Apis mellifera* | AmFAS | XP_006567467.1 |
|  | AmFAS-like | XP_395426.4 |
| *Anopheles gambiae* | AgAGAP001899 | XP_321166.4 |
|  | AgAGAP009176 | XP_319941.4 |
|  | AgAGAP008468 | XP_316979.4 |
| *Saccharomyces cerevisiae* | ScFAS1 | NP_012739.1 |
|  | ScFAS2 | NP_015093.1 |
|  | ScCEM1 | NP_010983.2 |

**Table S2. Primers for generation of dsRNA for knock-down studies**

| **Target genes** | **Primers** | **References** |
| --- | --- | --- |
| *Aa*FAS1  (AAEL001194) | F- 5’ GTCCCGTGTGTCTACTCAACAGT 3’  R- 5’ GCTTGTTCCACCTGTTGGTC 3’ | (12) |
| eGFP | F- 5’ GACCACATGAAGCAGCACGA 3’  R- 5’ CGCTTCTCGTTGGGGTCTTT 3’ | (20) |

* * T7 promotor sequence (5’ GAATTAATACGACTCACTATAGGGAGA 3’) was added to the 5’ end of all primers to make dsRNA

**Table S3. Primers for *Aa*FAS expression analyses**

| **Target gene** | **Primers** | **References** |
| --- | --- | --- |
| *Aa*FAS1  (VB: LOC5568814) | F- 5’ GAGGTCGTCCGATTGGTTTC 3’  R- 5’ AGGACAACCTTGCCGATGTG 3’ | (12) |
| *Aa*FAS2  (VB: LOC5570229) | F- 5’ CATTTCAAGCAGGCCCACAC 3’  R- 5’ TCTCAGACTCGGCAAAGCAG 3’ |  |
| *Aa*FAS3  (VB: LOC5573929) | F- 5’ GTGCAGCTTCGAGATGCCATA 3’  R- 5’ GCTCCATCACAGAGTTTGCC 3’ |  |
| *Aa*FAS4  (VB: LOC5573931) | F- 5’ GCTATGCTGGGATGTGCCAA 3’  R- 5’ TTCCTACGAGCTACATCCATAGC 3’ |  |
| *Aa*FAS5  (VB: LOC5573927) | F- 5’ GGGACTGCAAGCCTTCAACA 3’  R- 5’ ATCCTTGGACAACACACCCA 3’ |  |
| β-Actin  (VB: AAEL004616) | F- 5’ GAATGTGCAAGGCCGGATTC 3’  R- 5’ GCTCGATCGGGTACTTCAGG 3’ | (20) |

VB, VectorBase accession number; GB, GenBank accession number

**Table S4. Primers for RT-PCR assay detecting mRNA products *Aa*FAS-like and *Aa*FAS6**

| **Target genes** | **Primer** | **Primer set** | **Expected band size (bp)** |
| --- | --- | --- | --- |
| *Aa*FAS1 (ketoacyl associated domain) | F- TCGGTGGTGCCAACGCTCAC | A | 1976 |
| *Aa*FAS1 (PS-DH domain) | R- GCGGGCCGGGTTGATCACAA |  |  |
| *Aa*FAS-like (ketoacyl associated domain) | F- GATGCAATGCCACTTTCGGG | B | 1379 |
| *Aa*FAS6 (PS-DH domain) | R- TCAGATTTTGCTTCCAACACCG |  |  |
| *Aa*FAS6 | F- TTCCGTGATGTGATGGTTGCT | C | 172 |
|  | R- TCGAATCCGCGTTAACCATCG |  |  |
| *Aa*FAS-like | F- TGAATCCATTGACCCTGCGA | D | 232 |
|  | R- TACGCAACATCCAAGGCGTA |  |  |

**Table S5. Amino acid similarity of FAS domains between *H. sapiens* and *Ae. aegypti***

| Domains | Percent Amino Acid Identity to domains of *H. sapiens* FAS (NP_004095.4) | | | | | | |
| --- | --- | --- | --- | --- | --- | --- | --- |
|  | ***Aa*FAS1** | ***Aa*FAS2** | ***Aa*FAS3** | ***Aa*FAS4** | ***Aa*FAS5** | ***Aa*FAS-like** | ***Aa*FAS6** |
| Human FAS  (full-length) | **45.3** | 36.7 | 35.7 | 32.9 | 34.6 | 36.1 | 36.8 |
| KS | **55.5** | 47.6 | 44.9 | 47.6 | 43.7 | 44.9 | N/A |
| Linker | **31.2** | 27.2 | 23.5 | 24.7 | 23.5 | 23.5 | N/A |
| MAT | **48.1** | 38.8 | 21.2 | 22.7 | 21.2 | 20.1 | N/A |
| DH | **47.5** | 35.3 | 32.1 | 32.8 | 32.1 | N/A | 26.3 |
| ΨME | **23.0** | 23.3 | 21.9 | 19.0 | 16.2 | N/A | 21.2 |
| ΨKR | 27.0 | 30.2 | 34.4 | 32.3 | 34.4 | N/A | **36.5** |
| ER | **59.3** | 45.7 | 44.2 | 47.9 | 47.2 | N/A | 43.3 |
| KR | **63.6** | 54.7 | 51.3 | 55.5 | 50.0 | N/A | 50.4 |
| ACP | 49.1 | 43.9 | 48.2 | 48.2 | **50.0** | N/A | 48.2 |
| TE | **34.5** | 21.6 | 21.4 | 18.2 | 20.8 | N/A | 20.6 |

Note: Bold numbers highlight the *Aa*FAS with highest percent amino acid identity among members of the gene family*.* Abbreviations: KS, β-ketoacyl synthase; MAT, malonyl-acetyl transferase; DH, dehydratase; ΨME, pseudo-methyltransferase; ΨKR, pseudo β-ketoacyl synthase; ER, enoyl reductase; KR, β-ketoacyl synthase; ACP; acyl carrier protein and TE, thioesterase.
