## Additional file 2_Chotiwan et al.docx for "Expression of fatty acid synthase genes and their role in development and arboviral infection of *Aedes aegypti*"

**File S1. mRNA and amino acid sequences of 7 *Aa*FAS genes.** mRNA and amino acid sequences of *Aa*FAS genes are shown and separated into each exon. Start codon is highlighted in turquoise, step codon is highlighted in red, possible SNPs between the AaegL5 and AaegL3 assemblies are highlighted in yellow and the conserve motif DTACSS, important for ketoacyl synthesis is highlighted in olive.

**AaFAS1 LOC5568814 (AAEL001194)**

**XM_001658130.2 PREDICTED: Aedes aegypti fatty acid synthase (LOC5568814), mRNA**

>Exon_1 GGTCACCTTTTCACTACTCAGACATCAGCTTTCGAGTGCGGTGGCGTGTTTTTCCTCCTCGTCGTCAAGTCGCTTTTCCTACACCATTATTGTGTGACATTTCAAGTGCATTTTTAATTGTGCAAGAGTCAAGAGATAACGCAGTGAATATCAGCATCATTTCGTACAATCCAGTCAGTTTATCTGTTCTCCCGGTAGCAACAGGGCAGAAGCCATCCGACGAAAGTGTGAACACATTCCTAACGCAATAGCAGCAGATCAGCGTTTGTTTAAAAAG

>Exon_2

CTGATTCAGAACATTTTGGGGAAAGACTTGCGCGAAGACAGCGGAGACACGCCGGCATACCGCCAGAGGAGTGTGCATAACAACAATACGGACTCGGCTACATAGTCAGCGGCGGTGTTGGGAAGCATCAAGCGGAGGAGTTATCAGTGCGAATTTCACGAGACGGGAGAGTGTCCGCGGAGAAAGAGTGTAATCGAAAAAGAGAGCGCGCGCGAGAGGACCATTCCTCGCGTCGGAAGTAAATAAGTACAAGCCAACGCCGTGAAGAAGGTTTAGTTTGAAGCAGTGCTTCAACGCGTTCAGATCGTGAAGTAATTTCTCAGAAGGAGCTGTAATCTCACAAATCGCGCGGATTGATCCCTTGGCAACCGCTACCAACAGCTTCCCAAACGGACAATCAAAAAAGAAAAAGTGTGTGAATTATTGCTTTATGTGGTTTGATGAGCATCAGGATCAATAAAAGCGATTGAAGCAGCAAAAACAAGTGATAACAGAGTGCACCAGAACAGTGAAGATCAATAACGACAACAGCGAGGTACTGTTCCAGTGCGCACCCTGATAAAAAAAGTGGTGTGATCCAGTCGGAAGCCGAAGTGTTTGTGTGTGCGAGAAGTTTAGGAAGAGAGTCCAGTGTGTAGTTGCTTCGCGAGAAATTGAAGATCCTTCTGCGTTGACCGTTCCCCGTTCAAGAATCTGATCTGAAGCGTTGCAGTCTGCTGAGGAGTTCAGTCGTCTTCTGGACTTACTAGTGAGAAGGTGTTCTTCAAGTCTGCTGACTAGTGAGCGGAGGATCCTTTTTTTGGGATTCCAGTGAAAAGTTTTCATCGCCCAATCGTTAGCTGAATTAACG

>Exon_3

ATAGACTAGACGTCGTCCAGCACTTCGCGGCTTGGTCAGAAAGCAGGACACCCTCTAGCCGATCGTCAGTGCAGCGGACAAGGCCGCCATTCCTTCCCGCGGAAAACGTTCCAAACAGAAGAAAACACGACGCAACAAAGGCGCAACGATGCCAGCCCGCTTCGACGATGTTACTACTGACACCCGCCGCGGTGTCCCCAGGGACCTGGGGGGACACTACGATGGCTGTCAGATTCAGGATGACATCTGCATCACAGGCTTCTCCGGCCGCCTGCCGGAGAGCTCCAACATCGAAGAGTTCAAGCGCAACCTGATGGAGGGCGTCGACATGGTGAACGACGATGACCGCCGGTGGCCCAAGGGATTGTACGACCTCCCGACCAGGATCGGCAAAATTAAGGACGAAGACCTGCAAAACCTGGACGCAGAGTTTTTCAAAATTCATCAAAAACAGGCGGAGTGTATGGACCCGCAGATGAGAATGCTGCTCGAGTGTACATACGAGGCTATTATTGATGCG-G

*Nucleotides G from exon_3 and GC from exon_4 code for amino acid “G”*

>Exon_4

GC-ATCAATCCTCAGGAACTCCGCGGTAGCCGCACCGGCGTGTACATCGGATGCTCCAACTCCGAAACGGAACAGCACTGGTGTGCCGACCCGGACCTGGTAAATGGGTATGGCCTGATCGGATGTGCCCGAGCCATGTTCGCCAACCGGTTGTCCTTCACGTTCGACTTCAAGGGACCCAGCTACGCGGTCGATACGGCCTGTTCCAGTTCGTTGATCGCCATGTCCACGGCCTTTGCCGATATGAAGGCTGGGCGGTGCGACGCGGCCATCGTGGCCGGGTGTGGAATCATTCTGAAGCCCACCATGTCGTTGCAGTTCAAAAGGTTGAACATGTTGGGTAAGGAAGGTATGTGCAAGGTGTTTGACGAATCTGGCAATGGGTACGTGCGGTCAGATGGTTGCGTAGTTACTTTCATGCAACGTGCTTCCGATTCGCGCCGTATCTATGCCAGCGTTCTGAACGTGCGAATCAACACCGATGGGTACAAGGAACAGGGAATCACCTTCCCGAATGGAGCCATGCAAAAGCGCCTGATTCAGGAAACCTATGGCGAAATCAATCTCAACCCAGCGGATGTGGTCTACGTGGAGGCTCACGGTACCGGAACGAAAGTCGGAGACCCGCAGGAGGTGAACAATATTACGGATTTCTTCTGCAAGGACCGGAAGGCGCCCCTGCTGATCGGTTCGGTCAAATCGAACATGGGTCACTCGGAACCTGCCTCCGGAGTTTGTTCCATCGCGAAAATGTTGATCGCTATGGAGGCGGGCATCATTCCCGGAAATCTTCATTTCAAGAACACCAACCCGGACCTTTACGGTATCATGGACGGCAGCGTTAAGGTTGTCGATCGTAACCTGCCATGGAATGGTGGCATAATCGGGCTGAACTCGTTCGGGTTCGGTGGTGCCAACGCTCACGTGATCATGAAGTCATACCCGAAACCGAAACCTGTCAGCCCCAAGGATGGCTTCCCTAAGCTGGTTTTTGCTTCGGGACGTACCGAAGAAGCCGTTGAGGCATTCTTGGATGCCGCAGACACGAACAAGGACGATGAGGAGTTTGTTGGACTGGTGAATGAAGTTCACAGTAAGAACATCCCTCTGCATTACTACCGTGGCTACACGGTATTGGGAGACGGACAAGCTGTACGAGAGGTCAGCGATTTGAATGACGAAAAGCGGCCAATCTGGTTCATCTACTCCGGAATGGGATCGCAGTGGGCCAGTATGGCCAAGGAGATGATGCAGGTGGAAGTGTTCAACAACAGTATCCATCGCTGTGCAGAGGCCTTACGACCGGAGGGAGTCGACCTGATCGATATCTTGACCAAGAGTGACGAATCGCGGTTCGATAACATCTTGAACTCGTTCATTTCGATTGCTGCGGTGCAAGTGGCATTGACCGACGTTCTGAACCATCTGGGCATCACTCCGGACGGTATGGTCGGACACTCCGTTGGTGAGCTGGGCTGTGCCTATGCCGATGGATGCTTCACTCCGGAGCAGACTGTTCTGGCTGCCTACTGGAGAGGACGCAGCATTTTGGATACCCAGCTGATCCCTGGCCTGATGGCTGCTGTGGGCCTCTCGTGGGAGCAATGCAAGGAACGCCTCCCGAAGGACATCATTGCCGCTTGCCACAACAGCAGCGATAGTGTGACG

>Exon_5

ATTTCCGGACCGGTAGATTCTATCAACAAGTTCGTCGCCGAGTTGAACGCCGAAGGCGTCTTTGCCAAGGCAGTCAAATCGTCCGGCATTGCTTTCCACAGTCGTTACATTGCCGACGCCGCACCAAAACTGCGCAAATCGCTAGACAAAATCATTCCGAACCCTAAGAGCCGTTCACCGCGCTGGATCAGTACTAGCATTCCGGAGGAAGCCTGGAACACTCCGTTGGCCCAACAATCGTCCGCCGCCTATCACGTGAACAACCTCCTATCGTCGGTGCTGTTCGCTGAAGGATTGCGGCACGTGCCCAGCAATGCGATTTGCATCGAAATTGCTCCGCACGGTTTGCTGCAGGCCATTTTGAAGCGCGCCCTCGGCAAAGACGCCACTAATTTGAGTCTGATGAAGCGCGGTCACGACAACAATGTGATCTTCATGCTGTCGAACATTGGA-AA

*Nucleotides AA from exon_5 and A from exon_6 code for amino acid “K”*

>Exon_6

A-TTGTACGCCGCCGGAGCTCAACCGCAGGTTCAGAAGCTGTACCGCCCGATTACGTACCCGGTTGGTCGTGGAACGCCCATGTTGAACTCACTGGTCAAATGGGACCATTCGACCAAGTGGTACCTAGCCAGATTCGGTGTTGAA-A

*Nucleotides A from exon_6 and AC from exon_7 code for amino acid “N”*

>Exon_7

AC-AAATCCGGCGAAACGGTCATCGACGTCAACCTGGAGAAACCGGATGATGCCTATCTGGCCGGACACACCATCGATGGACGTGTCCTGTTCCCTGCCACCGGTTACATGACCCTGGCCTGGCGTACATTCGCCAAAATGCGCGGTTCGGACATGGAAAAGACGCCGGTGGTCATCGAAAATGCCGTTTTCCACCGGGCCACCATTCTGCCGAAGGATGGTTCGGTCAAGTTTGGAGTGAATTTCTTCGATGGTACCGGGGCTTTCGAAATCTGCGAAGGTGGATCGTTGGCCGTGTCGGGCAAGATCACCGTTCCGGAAAATATCGACAACGAGGAGTTGCCGTTGTATCCGATCGATGAGGACAAGTCGGGTCTGTCGATGAGTACCGGTGATGCCTACAAGGAGTTGAGATTGAGAGGATACGACTACGGTGGTATGTTCCGTGGAATTACGAAGGCTGATGCCAGCCGAGTTACTGGTGAGCTGCAGTGGAAGGATAACTGGGTTAGCTTTATGGACACCATGCTGCAGTTCAGCATTTTGGGCAAGAACATGAGGGAGTTGTACTTGCCCACTAGAATTGAGAAGATTGTGATCAACCCGGCCCGCCACATGGACATGATGAACACCCTGAAGTCGAAGGATCAAGATGTTCCGGTCGCTGTCTACCGCAACATCGATGTGATCAAGAGTGGAGGTGTGGAGATGCGAGGATTAAAGGCCACACTTGCTCCGAAACGTTCGGGATCACAGGCTCCGCCAACACTTGAGAAGTATGTCTTTGTACCGAACTTCAACGAGAAGGAATTGGCAGAGAGTGGCGAGAAGTCCCAATTCCGGTCAATTTCCACGGCTGTCCACTTGGCTATCGAGAACAGCGCAGGAGCCCTGAAGATCAAGGTTGCTGAAGCATGCTTCCAACGTGCTCCGGAGAATACCATGGCTGCTACCGTGCAGGCCATCATTGAAGGCGAACCTACCCTGGTGAGTGACGTAGCAGTAGTTACGAACTATCAACCAGATACCCTTGTTCAGCACTTTGGAGATAGTGGAGTTCGTGTAGTGGCCAAGGACGCTGCGGCAGGACCTATTGAACAAGGTTGCCATATGGTAGTTTCTTACGATATACTTGGCCGTTCGGACAGCCAAGCTGTTCTGAGCAATCTCCGTGAATCGATCCGTGATGATGGATTCCTGTTCCTCGAAGAATCTCGCACAAACTTTGATGCTACCAAGAAGGGTAAGGCTCTGTTCGATTCGCTAAAACTGACGACCGTGACGTATCAATATTACGACAAAAAGGTCTTTGTACTACTCCGTCCCACCAACAACTACGAGCAACGCAAGTCCACAGTTATTCAGGTCACCGAGAAGAACTTCACCTGGGTGGAAACGTTGAAGGCTGCTTTGGCGAAGGCCGAAGAAACCAATACCTTCGTGTACATTGTATGCCAGGGCGAAGAGCTGTTCGGCGGTCAAGGATTCATCAATTGCATCAAGAACGAGGCTGGCGGTAAGTTCGCCCGTATGGTCTTCATTCAGGACAAGCGAGCGGAGAAATTCTCCCTCACCGGCAAATCGTACGTTGAACAGTTGAAGAAGGACCTCATTTGTAACGTTATGAACAGCAGTGGCGTTTGGGGAAGCTTCCGGCATCTGAGATTGGACAACCAGACCAACGGAACATCGCTGCAGGTCGAGCATGCCTATGTCAATGCCTTGACAAAGGGCGACTTGGCGAGCTTGAAGTGGATCGAAGGACCACTGTCGCGGGACAGGCCAGATCCGAAGGACAAGAAGCAGGAGTTGTGCACCGTTTACTACGCTCCGATCAATTTCCGTGACGTAATGTTGAGCTCTGGCAAGCTTGGAGTTGATGCTCTGCCGGGAGATCTGCCCACTCAGGACTGCATTCTCGGATTGGAG

>Exon_8

TTCTCCGGTCGGGACGCAGCCGGTCGTCGCATCATGGCCATGGTTCCAGCCAAATCGCTGGCAACTACCTGCGTAGCTCACCGGAACATGATGTGGGAAATTCCGGACAACTGGACTATGGAGCAAGCCTCGACAGTCCCGTGTGTCTACTCAACAGTTTACTACGCCCTGGTCGTGCGTGGTCGCATGAAGCGTGGTGAATCTATTCTGATTCACGCCGGATCCGGTGGCGTGGGACAGGCCGCTATCTCGGTTGCCCTTTCCGCTGGACTCACGGTCTTCACCACCGTTGGTAGCAAGGAAAAGCGTGACTTCCTGAAGCGTACCTTCCCACAACTGCAAGATCGCCACATTGGAAACTCCCGAGACTGCTCGTTCGAGCAGATGGTCATGCGTGAAACTCAAGGACGTGGCGTGGACCTGGTACTGAACTCGTTGGCCGATGAGAAACTGCAGGCATCGGTCCGATGCTTGGGATTGAACGGACGGTTCTTGGAAATCGGCAAGTTCGATTTGAGCAACAACAGCCCGCTGGGTATGTCGGTGTTCTTGAAAAACACTTCATTCCACGGCATTCTGGTCGACAGCATTATGGACGGCGATGATGACGTGCTTGCCGAGGTCGTCCGATTGGTTTCGGAAGGTATCAAGAGCGGTGCGGTACGCCCGCTGCCAACTAGCGTGTTTGGCGACCAACAGGTGGAACAAGCTTTC-CG

*Nucleotides CG from exon_8 and A from exon_9 code for amino acid “R”*

>Exon_9

A-TTCATGGCTTCCGGTAAGCACATCGGCAAGGTTGTCCTCAAGATCCGCGACGAAGAGAAGGCCAAGGTGGTCGTCCCGCCCCCAAAACTGATCTCCGCTATCCCTCGTACGTACATGCACAAGGAAAAGAGCTACATCCTGATCGGTGGACTGGGAGGTTTCGGTCTGGAGTTGGCCAACTGGTTGGTCTCGCGCGGAGCTACCAAGATCATTCTGACCTCTCGCAGTGGAGTTCGAACCGGATACCAGGCCCTGATGATCAAGCGTTGGACCGAACGTGGCGTTACGGTTTCGATCGATACGAACGATGTCACTACTTTGAAGGGAGCACAGAAGCTACTGCAGGCCGCCAACAAGCTGGGACCGGTCGGAGGTGTGTTCAACTTGGCGGCGGTGCTGCGGGATGGTCTGTTGGAAAATGCAACAGAAGCGGACTTCAAGACTGTCTGTGTGCCTAAGGTTGACGGAACGAAGAACTTGGATCAGGCCACGCGGGAGCTGTGCCCGGACTTGGATTATTTCGTTGTGTTCTCGAGTGTTTCGTGCGGACGAGGAAACATTGGACAGGTGAACTATGGATTGGCCAATTCGGCCATGGAGCGAATTTGCGAAGCACGTCAAGCCGTCGGTTTGCCTGGAACTGCCATTCAATGGGGCGCCATCGGAGACACCGGTTTGGTGTTGGAGAACTTGGGTGACAACGATACGGTTGTCGGTGGAACTCTCCCGCAAAGAATGCCATCCTGTCTTCAGACAATGGACTTCTTCATGCAGCAGCCTTGTCCAGTTTTGGCTTCGATGGTTGTGGCTGAGAAACGTAAGGCGGAAACCGGAGGAGTTGGATTGGTCAGCTGCGTGGCGAACATTCTTGGTTTGAAGGATACCAAGAACGTGTCCGATTCCGCCACTCTGGCTGATCTCGGAATGGACTCCTTGATGGGCGCTGAAATTAAGCAAACCCTTGAACGAAACTTCGACACTGTTTTAAGTGCAGCGGAAATTCGTATGCTCACATTCGGCCGTCTGAAAGCGCTGGAAAGTGGTGGCGCCGATGCTCCTGGAAGTCCCGTCCCACCGGCATCTGCATCGCCCGAACCAGAGCAGCAAGATCAAAATGGACTGGGAGATGGCACTCAGGTTAGATTCAGTGCAGATCTCATGCCTACGCAGTGCCTGGTTCGATTGGACTCCAAGGCCCCGCCATCCAGCAAGGACCGCCCAGTGTTCATGGTACATGCCATTGAAGGTGTCATCACTTCGCTCATTCCACTAGCACAAACCCTCCCAGTTCCTGTGTACGGTCTCCAGTGTGTAGCCGAAGCACCCCTAGAATCTCTGGAAACGTTGGCCGCGTTCTACATTAAGCATGTCCGATCGGTGCAGCCAAAGGGTCCTTACACCATCGTGGGCTACTCGTTTGGAGCGTCCGTCGCCTATGAAATGGTGGCCCAGTTAGAAAATGCCAAAGAAACCTGTCGATTGCTGATGCTGGACGGTTCGCCACGATACATCGCTGGTTACACGGACGCACAAAAGCAGCGAATCGACAACGGAGAAGCCCTCCAGGCCGAGGATGAAGCCTATGCGCTGGCCTTCTTCGCCATGGTCTGCGGCAATCTGGACTACTCTAAGACTGCCCACGAACTGATTGCACCGAAGACGTGGGAAGCGCGGTTGCAGAAATGCGCCGAAATGGTTCGCACGCGAACGCCACAATACTCTCAGAAACTG

>Exon_10

CTCGAGACGGCAGCGGAATCGTTTGTACATAAAATCGTTGCTGGCCATCTGTATAAACCCACTTCCAAGATCAGTGCCCCGGTGAAACTGGTGAAACCCACCGAAAACTATGCCAAACTGCAAGGCGATTATGGTCTCTCAGAG

>Exon_11

CTCTGCACGAAGGACGTCAAGGTGACGACTGTGAAAGGCGACCATCGATCCATTCTTGTCGGTGATTCGATGCTGGAAATCTCTAAACTTCTCCACGAGCTGCTATAAAGTGAAGCGTATCGTCATCACACCTTCGCACACAGGAAACACCTGCATGGACATTAGAGCATAGAAATCACTGGGGCGTACCGCCTTTTCCCCCAATCTCTCGAAGCGAGTGAGAGTCAGATGGAATGTGTAATTTATTTTCGTTCTTTTGTGTATAGTTCAAATCTACTTCCCGACTTTTCTACATTAATTTTGGTTATAAGATGGTTCCCAGTGGACCTATTGGCACTGTTAACGCAAGCGTGATCTATATAGAACTAGTAGGATGCAGTCCAAAGAGTGCTGGCACTGGACGTCTGATCCAGGTCCGTTTCAGGGGCAGTGTCACCGCCCTTTGCTGTAGTTTCGAAAACATGAAAATGTATAAAAAAGAACGTGATTGCTTATTAGCTGTTAGTTGCCTTCTTTCTGAGAAGCAAAAAAAAAACTTCGACATGAACATTTTCCAAGATCTTGATCTATTTATGAAATGGATTTGAAGAATGTGTTAGTTGCATAGCAGACAACATTCCACTCCTAGAGTAGCTATTTTTTTCCTTTAAAATAATACTTTAATTATAAAAAGTAGATTTTAGTGAGTCGATGGGATTTTATTCCTAAAACAACTTGTAAAAAAAATAGTTATTTTACATCACAGAAATGACAAAAATAAAGAAGGGATCTATTAACCATGTGTACTAAATTTATTCTGTCTTCGACCCAACGAAAAACAATCGGACAAATGATCGAACACACAAACAATTTTACAAAACAGGGGCATCCAGGTTTAGCATGTAAGGACCATTAACTATCATAAGCAGCCATTCAACTTCTATTACCATGTGTGATGCAGAAATCTTACCCGGTACAGGCACATGATCAATCTACTCGAAGTCACACTATCACACGTTTACTGAAGTTGTTGTGTGTAATCACGAATATTTAACAATATTGTAAACTAGCACTACACTTCTGTTGCGGAACACACACACACCCGAAAGCGGATAAGTAAAGCCTATCAAATCTCAAAGCACCAATTGCGTATACCCCCAAAATGGCTTATTGCACTCACCCACACAGAAGTACTGAATGTATGTATTTGAACAGGAACATAGTTATTTGTGCATTTGGAAGAACAAACGGCCAATTATGATGGCAAAACAGGAAACTGTATATTTGAATGCAAATAAAAATATTGCAAATGAAAAA

**LOC5568814_amino_acid_sequence**

>Exon_3

MPARFDDVTTDTRRGVPRDLGGHYDGCQIQDDICITGFSGRLPESSNIEEFKRNLMEGVDMVNDDDRRWPKGLYDLPTRIGKIKDEDLQNLDAEFFKIHQKQAECMDPQMRMLLECTYEAIIDA-G

>Exon_4

INPQELRGSRTGVYIGCSNSETEQHWCADPDLVNGYGLIGCARAMFANRLSFTFDFKGPSYAVDTACSSSLIAMSTAFADMKAGRCDAAIVAGCGIILKPTMSLQFKRLNMLGKEGMCKVFDESGNGYVRSDGCVVTFMQRASDSRRIYASVLNVRINTDGYKEQGITFPNGAMQKRLIQETYGEINLNPADVVYVEAHGTGTKVGDPQEVNNITDFFCKDRKAPLLIGSVKSNMGHSEPASGVCSIAKMLIAMEAGIIPGNLHFKNTNPDLYGIMDGSVKVVDRNLPWNGGIIGLNSFGFGGANAHVIMKSYPKPKPVSPKDGFPKLVFASGRTEEAVEAFLDAADTNKDDEEFVGLVNEVHSKNIPLHYYRGYTVLGDGQAVREVSDLNDEKRPIWFIYSGMGSQWASMAKEMMQVEVFNNSIHRCAEALRPEGVDLIDILTKSDESRFDNILNSFISIAAVQVALTDVLNHLGITPDGMVGHSVGELGCAYADGCFTPEQTVLAAYWRGRSILDTQLIPGLMAAVGLSWEQCKERLPKDIIAACHNSSDSVT

>Exon_5

ISGPVDSINKFVAELNAEGVFAKAVKSSGIAFHSRYIADAAPKLRKSLDKIIPNPKSRSPRWISTSIPEEAWNTPLAQQSSAAYHVNNLLSSVLFAEGLRHVPSNAICIEIAPHGLLQAILKRALGKDATNLSLMKRGHDNNVIFMLSNIG-K

>Exon_6

LYAAGAQPQVQKLYRPITYPVGRGTPMLNSLVKWDHSTKWYLARFGVE-N

>Exon_7

KSGETVIDVNLEKPDDAYLAGHTIDGRVLFPATGYMTLAWRTFAKMRGSDMEKTPVVIENAVFHRATILPKDGSVKFGVNFFDGTGAFEICEGGSLAVSGKITVPENIDNEELPLYPIDEDKSGLSMSTGDAYKELRLRGYDYGGMFRGITKADASRVTGELQWKDNWVSFMDTMLQFSILGKNMRELYLPTRIEKIVINPARHMDMMNTLKSKDQDVPVAVYRNIDVIKSGGVEMRGLKATLAPKRSGSQAPPTLEKYVFVPNFNEKELAESGEKSQFRSISTAVHLAIENSAGALKIKVAEACFQRAPENTMAATVQAIIEGEPTLVSDVAVVTNYQPDTLVQHFGDSGVRVVAKDAAAGPIEQGCHMVVSYDILGRSDSQAVLSNLRESIRDDGFLFLEESRTNFDATKKGKALFDSLKLTTVTYQYYDKKVFVLLRPTNNYEQRKSTVIQVTEKNFTWVETLKAALAKAEETNTFVYIVCQGEELFGGQGFINCIKNEAGGKFARMVFIQDKRAEKFSLTGKSYVEQLKKDLICNVMNSSGVWGSFRHLRLDNQTNGTSLQVEHAYVNALTKGDLASLKWIEGPLSRDRPDPKDKKQELCTVYYAPINFRDVMLSSGKLGVDALPGDLPTQDCILGLE

>Exon_8

FSGRDAAGRRIMAMVPAKSLATTCVAHRNMMWEIPDNWTMEQASTVPCVYSTVYYALVVRGRMKRGESILIHAGSGGVGQAAISVALSAGLTVFTTVGSKEKRDFLKRTFPQLQDRHIGNSRDCSFEQMVMRETQGRGVDLVLNSLADEKLQASVRCLGLNGRFLEIGKFDLSNNSPLGMSVFLKNTSFHGILVDSIMDGDDDVLAEVVRLVSEGIKSGAVRPLPTSVFGDQQVEQAF-R

>Exon_9

FMASGKHIGKVVLKIRDEEKAKVVVPPPKLISAIPRTYMHKEKSYILIGGLGGFGLELANWLVSRGATKIILTSRSGVRTGYQALMIKRWTERGVTVSIDTNDVTTLKGAQKLLQAANKLGPVGGVFNLAAVLRDGLLENATEADFKTVCVPKVDGTKNLDQATRELCPDLDYFVVFSSVSCGRGNIGQVNYGLANSAMERICEARQAVGLPGTAIQWGAIGDTGLVLENLGDNDTVVGGTLPQRMPSCLQTMDFFMQQPCPVLASMVVAEKRKAETGGVGLVSCVANILGLKDTKNVSDSATLADLGMDSLMGAEIKQTLERNFDTVLSAAEIRMLTFGRLKALESGGADAPGSPVPPASASPEPEQQDQNGLGDGTQVRFSADLMPTQCLVRLDSKAPPSSKDRPVFMVHAIEGVITSLIPLAQTLPVPVYGLQCVAEAPLESLETLAAFYIKHVRSVQPKGPYTIVGYSFGASVAYEMVAQLENAKETCRLLMLDGSPRYIAGYTDAQKQRIDNGEALQAEDEAYALAFFAMVCGNLDYSKTAHELIAPKTWEARLQKCAEMVRTRTPQYSQKL

>Exon_10

LETAAESFVHKIVAGHLYKPTSKISAPVKLVKPTENYAKLQGDYGLSE

>Exon_11

LCTKDVKVTTVKGDHRSILVGDSMLEISKLLHELL

**AaFAS2 LOC5570229 (AAEL008160)**

**XM_001658958.2 PREDICTED: Aedes aegypti fatty acid synthase (LOC5570229), mRNA**

>Exon_1

ATTTCACAGCAGTTCATCAAACTTTCATTGTGATTGAAACAAAACGAAAATAACAAACGAAATTTGAATTGAAAACTTCTAACTGGTATAGTCGAGCTAATCTGCATTAAATTCGGTTATTTGCCCGAATGCTTACTGCTGATGGTAGGTAACATGCATTTCTCACCCTTTCCATGTCACTCGAAGTACGTTTATTGCCATTCGGTACATACAAAGCAAGTTTCCATCGTTGATTCTAAGGGGGGTTAGAAAGGTTAAACATGGCTCACGACTTCTTCATTCCACAGTTACGAATTCGGAAAACTGTGTGGTGGCAGCACGTGAAACCCTATTTATAATAATATTTCTGAGTTTTATTATTTCGGACAATATTCGTTTTCGACAAGAATGAAACAAAATGCTGAGGGGTCATTCACAACTTACGCAACGGGATAGTCCCCGGTCGGCGTATTGACGCAGGAAGTTCTTTTCCCAGTGTCGCACGTTCACCGGTCAAAGCTTAATGAAAGTAAAGCGTCGCTAATCAGATGCATGGAATCGTTAATCGAAAGTCGATGCAAGAATCGTAATCGAACCACAGCGTATTGGTCGCACAGGTCAACGCTATAAAACGGCTTCCCACCGTTGGTTCTTCGTTCACAAGCGTATTGGTCGTTGCGCCGCTGAGGTCTTACGTTGGGTTTTTCGTTGAACGGTCATCGCGCGGATTGTGTGCAAATTTCTTTCGGTTGGATAACTTCTGCTGGTTCGGTTTGCAAGTTCAATGTCAATGTCTGCCTAGTGGATTGTCAGCTCAACTAGGACCGGATACGGACGCAGGACTCAGGCAGACGCTTACGGGAACTCCTCTGGCGGAATTTGGAATCTAACGCTTTGTGAAGATCGAGTTGGTGAAGTGACTTTTAAAGGCTTGTATTTGGGAACGGTGCTAAGCATTTGGTGATTTGTGGAACTATTTGTAAGGGATAAAATTGGTGTTTTTGTGAAGTGATTAGTGAACGAACTGATAAACAGTACAAATTGATATAACAATGGGTTCGAGAATTGAAAATCCCGCGATCGAACATCGCGCCAAGCCTCCTATGCCCGGTGAGGAGATCGTGATAGCTGGTATTTCCGGTCGTTTCCCCAATTCGGACAATGTCCGTGAATACGCCCACAACCTGTACAACAAGATCGATATGGTGGACGAACTGGAAACTCGTTGGCGTCACAGCAATGTGGAAATTCCTCGCCGTATGGGAAAGGTCAACAACCTGGAGAAGTTCGACGCTACGTTCTTCGGTGTGCATTTCAAGCAG

>Exon_2

GCCCACACGATGGACCCGCAGTGTCGTATTCTGGTGGAACACGCCTACGAAGCAGTCATCGATGCCGGTGTCAATCCAAAGAGTCTTCGTGGATCGCGTACCGGAGTGTTCGTCGGAGCCTGCTTTGCCGAGTCTGAGAAGACTTGGTTCTACGAGAAGATTTCCTCCGGTGGATTCGGTATTACCGGTTGTTCCCGTGCTATGATGGCGAACCGTATTTCCTACACCATGGGTCTGAACGGTCCATCGTTCTTGTTGGATACCGCTTGCAGTAGTTCCATGTACGCTTTGGATTGCGCTTTCAACGCTATCCGCAACGGAGAATGCGACGCAGCTATTGTTGGCGGATCTAACCTGCTGCTGCATCCTTACGTAACTCTGCAATTTGCCCGTCTTGGAGTCTTGGCGGCCGACGGATATTGTCGCCCGTTCGACAAAGACGCCTCCGGTTATACCCGCTCGGAAGCCATTTGCGTTGTGTTCTTACAAAAGGCTAAGGATGCTAAGAGAATATACGCTGACGTCGTCTACTCGAAGACCAACTGCGATGGGTTCAAGGAGGAAGGCATCACCTACCCGTCAGGCCCGATGCAACGTAAGCTGCTTTCGGAATTCTACCAAGAAATCAACATCGATCCATCGACGGTTGACTACGTAGAAGCTCACAGTACCGGAACAGTTGTTGGTGATCCAGAAGAGTGCAACGGATTGGATAAGGTGTTTTGCCAAGGCCGCAAGAAACCACTGCCAGTCGGATCGGTCAAGTCTAACATCGGTCATTCCGAATCCAGCTCGGGTATCTGCTCTATCATCAAGTCGGTTTTGGCGTTCGAAAATCAACTTATTCCACCAAACATCAACTTCAGCGAAATCAGAGGAGACATTCCATCGCTGGTCGAGGGACGTTTGTCGGTTGTTTCGGAGCCTACACCTTTGGACGGCCCATTGATTGCGGTCAACTCTTTCGGTTTCGGTGGTGCCAATGCTCATGCTCTGCTTAAAGGTAACCCGAAGTTGAAGATTAACGGCGGTATTCCGGAGGATAACCTGCCACGTTTGGTCACATGGTCCGGTAGAACCGAAGAAGCCATCGATGTAATTCTAACTGAGCTTAGCAACAAACATCTGGACGCCGAATACGTCTCCTTGTTGCACAGCGTTCAGAGCGAGTCTATTCCCGGAAATGTGTTCCGTGGCTACGGAGTCTACGCCAAGAACGGAACGGAAAACGCTCTGTGCCTAGGACGTGATTGCCAACATTACACCGGTATGAAGCGCCCGCTGGTTTGGGTGTTCAGCGGTATGGGTTCGCAGTGGACCGAAATGGGTACTTCTCTTCTGGAAATCCCAATCTTCCGGGCTTCTGTTGAAAAATGCCATCAAGTGTTGGAGAAGAAGGGCCTGAATCTGATGGAGATTCTCACCTCTAAGGAATGCAAATATGAAAACATTCTTCACTCATTTGTGGGAATCGCCGCTGTTCAAATCGGTATAGTAGATGTCCTGCGATCGCTCGATATCGAACCAGACTTTGTGATTGGACATTCCGTCGGTGAGCTGGGTTGTGCGTACGCTGATGGATGCTTCACTGCCGAGCAAATGATTCTGTCTGCATACTCGCGAGGTATGGCCAGTTTGGAGACGAAGACCGTGTTTGGTTCGATGGCCGCTGTTGGTATGGGCTATCGCAAGATCCGTACGATGATTCCTCCAGGCATTGAAGTTGCATGCCACAACGGTCCGGATTCGTGTACCATTTCTGGACCCAAGGAAAACGTAGAGGCTTTCGTAAAGGAGTTGACTGGCAAGGGCATATTCGCCAAGGAAGTTCCTTGCTCGAACATTCCTTACCACAGCAAATACATCGCTGAAATGGGCCCAAGGCTGTTGGCACGACTGAACGAAGTCATCCCAGAACCGAAGAAACGCTCACCGAAATGGCTGAGCTCATCCGTACCGAAGATCCGCTGGGATCAACCGGAAAGCCAGTACTCATCGGCTCACTACCATACGAACAATCTGTTGAGCTCGGTACTGTTCGAAGAAACTTCCGCTCTGCTCCCGAACAACGCCATAACCATTGAAGTTGCTCCTCATGGATTGTTGCAAGCCATTTTGAAGAAATCTATGCCGAATGCTATCCATATTGGATTGACCAAGCGCGGTAATAAGGATAACGTGCAGTACATGTTCAACGCTTTGGGA-AA

*Nucleotides AA from exon_2 and A from exon_3 code for amino acid “K”*

>Exon_3

A-TTGTACGTCAATGGATTGGACATCCCGGTTTCGCGTCTCTATCCTCAAGTGGAATATCCCGTCTCCCGAGGAACGCCCCTGATTTCCCACCTGGTCCGTTGGGATCACAGCGAAGATTGGTTCGTGACCAAGTTCGAAATGCAGAAGTCTTCGAAGAGTGGCGAACGTCGTGTCAAGATCAAACTCAGCGATCAGGACTTCAGCTACATCGCTGGACACGTTATCGACGGTCGAGTGCTGTTCCCTGCCACGGCATACCTTCATCTGGCGTGGGAAACTCTTGCCATGGTTAAGGGCCCGATGTACTTCGACTTGGAGGTTGAGTTTGAGGATGTTAAGTTCCTGCGGGCTACGTCGATTGCCAAAGATCAAGAGATCGAGTTCACCGTGATGCTGCAGCCGGGTACCGGCCGATTTGAG

>Exon_4

ATCACCGAAGGGTCGGCAGCAGTCGTTACAGGCTACGTCAAGCACGTTGAAAACATTCAACTGTCGGAAATTCCAGACCCACCAGAAAGCGACTTCCCCATGCTGTCCGCTCGTGACTTCTACAAGGAGCTGCGTCTGCGTGGTTACCATTACAACGGTGCTTTCCGCGCTGTCCAGGAAGCTCGAGCTGATGCCCGTTGTGGAAAGGTCAGCTGGGATCTGAACTGGGTGTCGTTCCTTGACTGCTTGCTGCAGTGCTGTATTGTTGGAAAGGACACTCGTTCTCTGATGATTCCAACCGGAATCGAACGCTTGCGCATCGACCCGAAGATGCACCTGGCCATTGCCAATACCATGGAGGAAGGTGAACAGGTGTTCGACTTCAAGTTCTCCGATGTACTAAACACGATCCGTTGCGGTGGAATCGAAGTGATTGGTCTGCAGGCTAGTACTGTTGGACGCAGGAAGCCACCAGGTTATCCAGTTTTGGAATCGTATCAATTCATCTCGCATCTTCCAGCACCGAAGATGACCAAGGCCGATGCCGTACGAGTGTGCGTCCAACTTGCTCTGGAAAACATCCCAGTACCGAAGGTTAAGGCTGTGGAAACTGACTTCCATAACGACACTCCGATTACGCCGCTGTTTGCTGAAGCACTTGGTGATCTCCCATTGGTAACCGCCGATTTGATGTTCCTTTCTCAGCAACAGCTATCCCTCACTGGAGTACATGTTGAAGACGGTAAACTATCCACTCAATCGAGCTGTCTGTTCTTGATTTCCGTGAACTGTCTCGGCCGTCCGGAATTGATCGAAGAAGCAGCCAACAGCTTGCACGATAGTGGTTTCATCCTCTCGCGTGAAGCCGCAAACCTGGCCTTAGATAGCGTCGTGGCTCCTGCTGGATACCAACTGATTGCCCTAATTCCTGTTGAAGATGAAACCTACGTACTATTGCAGCGTATCAAGCGCAAGTATATGGGAAGTCCAACTATTGTCCCCATTCCTAATGATGACAAGTATGACTGGGTCGCTAAGCTGCGGGATGCGATGAAGCAAGGATCGGTGATCGCTGTGGCCCAAGGAGATAGCATGTCTGGTATTCTCGGACTGGTTAACTGTATTCGTAAGGAGCCCGATGGTGGCATGGTGAGCTGCGTGTTTGTTGATGATCCTAAGGCACCTGCTTTCGAACTGGACAATCCTGTATATAAGTCGCATTTGAAGCTTGGACTTGCATTGAATGTCTACCGTAATGGTCGCTGGGGATCCTACCGTCATCTTCAACTGTTGCAACCGATTGTCACCAAGCCAAGACGTGACCATTGCTATGCTAATGCTCTCACTAAGGGAGATCTCTCATCGATGACCTGGCTCGGCGGACCGTACAATCAGCGTAGACCGACACGTGAACTCGTCAAGGTTTGCTACAGTTCGTTGAACTTCCGTGATGTGATGTTTGCTTCGGGCAAACTTACGGCCGACTTTGCCAATCTGTCACGTCTCGAGCAGCAGTGTGATCTTGGTTTCGAGTATTCCGGAGTTACGGAGAGCGGAAAACGATTGATGGGAATGCTTACGACCGGAGCAATGGCAACTCATGTTGAGAATGATGAGTCGTTGACATGGATTGTTCCCGAACACTGGACTCTGGAGGAGGCGGCTACAGTTCCGGTAGTCTATGGTACAGTGTACACTGCATTGTTCCTCAACGCAAACATTCAGAAAGGACAATCCATCCTTATCCACGCTGGTAGTGGAGGTGTTGGTCTGGCAGCTATCAACGTATGTCAGGCATACGGATTGGAAGTGTACACTACGGTGGGAACACAAGCGAAGCGGGACTTCCTGCTCCAAACCTTCCCGAAACTAAAGCCTGAGAACATCGGAAACTCTCGTGACACTTCATTCGAGAAGATGATCATGCTTCGTACTAACGGCATGGGAGTTGACTACGTACTGAACTCGTTGTCCGAAGATAAACTGCAAGCTTCGTTGCGCTGCTTGGGCAAATACGGGAAATTCCTGGAAATCGGAAAATACGATATGGCCAACGACAGCAAACTGGGCCTGCATAGATTCTTGCGAGCCCTGTCGTTCACGGCCATCTTGGTAGATTTCCTGTTCACAGCTCCATTGGAGGAGAAACAGGTCTTGCGCGATCTGGTCGATAAGGATATCAAGGCCGGAATCGTCAAGCCACTGAAGACTAACGTTTTCCCTGCCAGTGAAATTGAGAATGCCTTCCGATATCTTGCTAGCGGTAAGCACGTCGGTAAGGTTTTGTTGAAGATTCGCGAGAACCCGAACGACGTTGAATCGGTTCCGATTGCTTATCTGCCGAGAATGTACTGCAATCCGGACCATTCGTACATCATCGCTGGAGGTTTGGGAGGATTTGGTTTGGAACTTGCTGACTGGTTGGTATTGCGTGGATGCCGCAAACTAGTGTTGAGCTCCAGCCGAGGTATTACCAAGCCGTACCAAGCCTACAGAATC-AA

*Nucleotides AA from exon_4 and G from exon_5 code for amino acid “K”*

>Exon_5

G-ATGTGGGAACAGTATGGAGTTCAGGTGGTAGTGAATACCGAGGACATCACGACCAAGAAGGGTTGCGAGGCATTGATAATGGCTGCTATGAAGTTGGGTCCAGTTGGTGGTATCTACAATCTGGCCGTACAGTTGCGCGATGGTATCTTTGAAAACCAGAGTGTGCAGAAGTTCATCGAGTGCATGGGTCCAAAGGCCGTTGCGACCAAGCATCTGGATGAGCTCAGCCGTAAGCTGTGTCCCAACCTGTCGTATTTCGTTGTGTTCTCCAGCGTGTCTTGCGGTCGTGGAAACGCTGGACAGAGCAACTACGGTATGGCGAACTCCGTCATGGAACGTATCATTGAACATCGTCATGCGAATGGACTGCCAGCTAAGGCTGTTCAGTGGGGAGCCGTTGGTGAAGTCGGTTTGGTTGCAGACATGCAAGAAGACAAGCTCGACATGGAAATCGGTGGCACTCTACAGCAAAGAATCTCATCTTGTCTTCAGGAGTTGGATCCTCTGCTGACAACTCCAGAACCAATCGTGGCCAGTATGGTTGTTGCTGAAAAGCGTGTGAGGAGCAGCGGCAAGGATAATATCATTGAATCTGTTATGAATATTATGAGCATTCGTGATATTAAGTCGGTATCGATGGACACGACGCTGTCGGAACTGGGAATGGACTCACTGATGGCCGTAGAGATCAAGCAGACCCTTGAACGTGAATACGAACTTTTCCTGACACCGCAGGACCTCCGATCGCTGACTTTCCTGAAACTTCAAGAGCTGACCGAAGCCAAGGCATGTTCCGACGACAACGTTAAACTGAAGCTGGCTAATGAAAAGACTCCAACTGGCGTTGCCATGTTGTTGCGTAACTTGGGCGATGAATTCAACAGCGAACATACGATTCTACGTCTTCAGTCCGAAAACGACTCGAAGAAGTACAATGCTTGCGTGCTCTTTACACCTGGAATTGAAGGCGTTGCCGGTAATGCCTGGCATAGTATCGCTTCGCAGCTGACTCTGCCTACGTTCATCACTCAACTAACCAAGACGATCAACATGAGCAGCATACCGGAGATTTGTCAATTCCTGTCCCAGGATGTGATCGAGAATGTGTTCAAGGGAACGGAACACTTCTATCTCGTCGGTTACTCGTTCGGTGCATTCATTACGCTGGAACTGGCACGATTGCTGGAGGAGACTGGTAAACGAGGAAAAATCCTGCTGATTGACGGAGCACCCAAGTTCTTGCACAAGTTGGCCCTGGATCAAATGTCCGATAACTGGACCGAGGAAACTGTTCAGATTGTTCTGTTCGCTGGCATCCTCAATACCATCTATCCAGATGAGACTACGGATGTGCTACCCATAATCAGCGGCTGCCCTTCGTTCGAGGCACGAATGGACAAACTGCTCGAACTTACCAAGGATCAGAACATCTACTCCGAATCATACCTGAGGATGATGACCAAGGCATTGTTCAACCGTATCAAGATCACCCTGCTCTACGACGTCAACAAGATTACCCCACTGGAGTCCCCGATCACCTTGGTTCGCCCAACGGAAGTGTCCGTCGTGGACATCGAAGAAGACTACGGACTGGCCGAGTACACCAAGGGAACGGTTAGTCTCAAGTTCCTGGAGGGTAACCACATCACCATGTTGGAGAACCCGAAGCTGACACAAATCATCACCGAGTCGGACCCAGTGCTCGAGTCGGATCGCTGCTTCCAGAAGTACCTTCAGAGCAATGTAATTGCCGAATAAGAATCGACCACGCCACGCCGATAACCGATAAAGCAACCCTAACAGCACGACCACTTCACCTTTTGCTATCGATTGTTGTTTGACAACCCAGGCACACACCTCTACTCCCAAAGAAACTATCGAATCCCAAGAAAAATCCAATCCCAAGTAAAGTTAATAAAAGAAAAGTTAATTTATAA

**LOC5570229_amino_acid_sequence**

>Exon_1

MGSRIENPAIEHRAKPPMPGEEIVIAGISGRFPNSDNVREYAHNLYNKIDMVDELETRWRHSNVEIPRRMGKVNNLEKFDATFFGVHFKQ

>Exon_2

AHTMDPQCRILVEHAYEAVIDAGVNPKSLRGSRTGVFVGACFAESEKTWFYEKISSGGFGITGCSRAMMANRISYTMGLNGPSFLLDTACSSSMYALDCAFNAIRNGECDAAIVGGSNLLLHPYVTLQFARLGVLAADGYCRPFDKDASGYTRSEAICVVFLQKAKDAKRIYADVVYSKTNCDGFKEEGITYPSGPMQRKLLSEFYQEINIDPSTVDYVEAHSTGTVVGDPEECNGLDKVFCQGRKKPLPVGSVKSNIGHSESSSGICSIIKSVLAFENQLIPPNINFSEIRGDIPSLVEGRLSVVSEPTPLDGPLIAVNSFGFGGANAHALLKGNPKLKINGGIPEDNLPRLVTWSGRTEEAIDVILTELSNKHLDAEYVSLLHSVQSESIPGNVFRGYGVYAKNGTENALCLGRDCQHYTGMKRPLVWVFSGMGSQWTEMGTSLLEIPIFRASVEKCHQVLEKKGLNLMEILTSKECKYENILHSFVGIAAVQIGIVDVLRSLDIEPDFVIGHSVGELGCAYADGCFTAEQMILSAYSRGMASLETKTVFGSMAAVGMGYRKIRTMIPPGIEVACHNGPDSCTISGPKENVEAFVKELTGKGIFAKEVPCSNIPYHSKYIAEMGPRLLARLNEVIPEPKKRSPKWLSSSVPKIRWDQPESQYSSAHYHTNNLLSSVLFEETSALLPNNAITIEVAPHGLLQAILKKSMPNAIHIGLTKRGNKDNVQYMFNALG-K

>Exon_3

LYVNGLDIPVSRLYPQVEYPVSRGTPLISHLVRWDHSEDWFVTKFEMQKSSKSGERRVKIKLSDQDFSYIAGHVIDGRVLFPATAYLHLAWETLAMVKGPMYFDLEVEFEDVKFLRATSIAKDQEIEFTVMLQPGTGRFE

>Exon_4

ITEGSAAVVTGYVKHVENIQLSEIPDPPESDFPMLSARDFYKELRLRGYHYNGAFRAVQEARADARCGKVSWDLNWVSFLDCLLQCCIVGKDTRSLMIPTGIERLRIDPKMHLAIANTMEEGEQVFDFKFSDVLNTIRCGGIEVIGLQASTVGRRKPPGYPVLESYQFISHLPAPKMTKADAVRVCVQLALENIPVPKVKAVETDFHNDTPITPLFAEALGDLPLVTADLMFLSQQQLSLTGVHVEDGKLSTQSSCLFLISVNCLGRPELIEEAANSLHDSGFILSREAANLALDSVVAPAGYQLIALIPVEDETYVLLQRIKRKYMGSPTIVPIPNDDKYDWVAKLRDAMKQGSVIAVAQGDSMSGILGLVNCIRKEPDGGMVSCVFVDDPKAPAFELDNPVYKSHLKLGLALNVYRNGRWGSYRHLQLLQPIVTKPRRDHCYANALTKGDLSSMTWLGGPYNQRRPTRELVKVCYSSLNFRDVMFASGKLTADFANLSRLEQQCDLGFEYSGVTESGKRLMGMLTTGAMATHVENDESLTWIVPEHWTLEEAATVPVVYGTVYTALFLNANIQKGQSILIHAGSGGVGLAAINVCQAYGLEVYTTVGTQAKRDFLLQTFPKLKPENIGNSRDTSFEKMIMLRTNGMGVDYVLNSLSEDKLQASLRCLGKYGKFLEIGKYDMANDSKLGLHRFLRALSFTAILVDFLFTAPLEEKQVLRDLVDKDIKAGIVKPLKTNVFPASEIENAFRYLASGKHVGKVLLKIRENPNDVESVPIAYLPRMYCNPDHSYIIAGGLGGFGLELADWLVLRGCRKLVLSSSRGITKPYQAYRI-K

>Exon_5

MWEQYGVQVVVNTEDITTKKGCEALIMAAMKLGPVGGIYNLAVQLRDGIFENQSVQKFIECMGPKAVATKHLDELSRKLCPNLSYFVVFSSVSCGRGNAGQSNYGMANSVMERIIEHRHANGLPAKAVQWGAVGEVGLVADMQEDKLDMEIGGTLQQRISSCLQELDPLLTTPEPIVASMVVAEKRVRSSGKDNIIESVMNIMSIRDIKSVSMDTTLSELGMDSLMAVEIKQTLEREYELFLTPQDLRSLTFLKLQELTEAKACSDDNVKLKLANEKTPTGVAMLLRNLGDEFNSEHTILRLQSENDSKKYNACVLFTPGIEGVAGNAWHSIASQLTLPTFITQLTKTINMSSIPEICQFLSQDVIENVFKGTEHFYLVGYSFGAFITLELARLLEETGKRGKILLIDGAPKFLHKLALDQMSDNWTEETVQIVLFAGILNTIYPDETTDVLPIISGCPSFEARMDKLLELTKDQNIYSESYLRMMTKALFNRIKITLLYDVNKITPLESPITLVRPTEVSVVDIEEDYGLAEYTKGTVSLKFLEGNHITMLENPKLTQIITESDPVLESDRCFQKYLQSNVIAE

**AaFAS3 LOC5573929 (AAEL022506)**

**XM_021847676.1 PREDICTED: Aedes aegypti fatty acid synthase (LOC5573929), mRNA**

>Exon_1

GGAAAATATTAGAATCCCTTCAATCCACTTTCGAAGCAGTTTGTGCGAGTTAATCGATCAAAAATGCCGACGCGTTGTATCCAATCTGTAGGAACGGATAAGAGTATCGTGATTAGTGGAATATCCGGCCGTTTTCCGCGGGCTAATAATGTGCGTGAATTTGCGAGCAGCCTGTACGGGAAGCAGGATCTTGTGGACGACCTGGAGACTCGTTGGCAACATACGATGCAGGATGTTCCACGACGTACCGGGAAGGTCGGGAACATGCAAAACTTTGATGCGGACTTCTTCGGAGTAAGTCGAATTGAGCGGGATACGATGGACCCACAACTGCGTATGACAATCGAGCATGTCTACGAAGCTATTCTCGATGCTGGAGTGAATCCTCAAACATTACGGGGCTCACGAACAGGAGTGTTCAGTGGAGTGTGTTTCTCGGAAACTGAAGTATGCATGTACTATAGAGCACGCCCTCCCAAGGGACTCGGTTTGTTG-GG

*Nucleotides GG from exon_1 and G from exon_2 code for amino acid “G”*

>Exon_2

G-TGTGCCAAGTCGCAGATTCCCAATCGTGTTTCGTACTTGCTAGATCTGAGAGGACCTAGCTATGTGCTGGATACTGCTTGTAGTAGCTCAATGTACGCCTTGGACGTTGCTTACCGAAGTATGATGAACGGCGAATGCGATGCGGCAATTGTTACGGGAGCAAATCTAACATTGCATCCCTTCATTACGTACCAGTTTGCAATGTTGGGAGTCTTGGCAAAGGACGGATATTGTCGGCCTTTCGATAAGGACGCAACGGGATACACTCGGTCAGAAGCCGTTTGCGCTGTATTTTTACAGAAAGCAAAAGATGCGAAGCGTGTTTATGGTCACATAATTCACTCAAAGACGAATTGTGATGGATTTAAACCGGAAGGAATAACTTATCCTTCAGGATCAGTACAACAACAGTTGTTAACTGAGTTCTACAACGAGGTAGGGATAAGTCCTACAGAGGTCGACTATGTAGAAGCACATAGTACTGGAACGTTTGTAGGAGATCCAGAAGAATGTGACGCTATAGATAAAGTGTATTGTTCCGGGCGAACCGATCCTCTGCTTGTAGGGTCTGTAAAATCCAGCATTGGCCATACGGAAGCTTCAGCAGGAGTATGCTCCATCACAAAATGCATTATCGCCATGGAGAATGGCCTTATTCCTCCAAACATCAATTATACTGATTACAGGCCTACGATTCCGTCTTTGGTTGAAGGTCGTTTGAAGGTTGTTACAGATGCAATGCCACTTTCGGGCCCATTGGTTGGTATCAACTCGTTTGGATTCGGAGGAGCAAATGCTCATGCTTTGTTATGTCGCAATTTGAAAGAGAAAGTTAAGAATGGCGTTCCAGACGATGATCTTCCAAGGTTGGTCACATGGTCAGGAAGAACCAGAGAATCTATTGAAACTATGCTTCATGACATTGGTCAGCGTCCCTTGGATGTTGAATTCATTGCTCTTCTGTTTAACATTCAGCAGCAGCCTACTCCTGGTCATCGGTATCGAGGATTTGGTATTTATCAGAAGAATGGAGATCGTCCAGCTGTACTACAAACATCATCCATCGATCGAGTGAAGCTTGATGCCATCCCAGTCGTTGCTGTTTTCGGAGGAATTAACACAAACTGGAGAAAAGAATTAGATGCTCTGCGCCAATTCTCCGTAGTAGAAGACACATTTGCTAAATGCAGCGGAATTCTGCGGTCGCTGAAGTTCGATTTGCACAAAAAACCATCAGGAAGAGAAAGCATATTGTACAATATGGTTGGAGCGACCATTCTTCAGTTGTCTATCGTTGATTTGCTGAGCTCGATTGGAGTGAAGTTTGACTTCTATGGAGGTCATTCCATTGGCCAATTCACTTGTGCATACATTGACCACAATTTGAACCTAGAACAAGTTCTTCGCTTAGCTTTCTGGCATGGATTAGTGTTTTCTGATTGTCATGCGGTTTGTGATCGCACTGCGTTTGTACAAATCAATTCAATGCTGAATCAGCTTCCATTGAAAAACTTTTTCAAGGATAGTGCAACCACCTTTGGAATTTTGACCGCAAATGAGAAAATTTTGATGGAACAGGTTCGTCAATTGAAATCGTCCGGCTTTGCAGCTGAAGAGTTGTCGTTTTTGGATGTGCATGCAGACTCAACGAAAAGTTCATCGCTTGCAAATAAACTTCGACAAACCGTCAATACCGTTTTGAGCAGAATCATTTTGCCCAGTGATAAATGGATTACTTCAGCATTGCCTCATACATCTTCTATATTCCATTCATCGAAACTGCACGATGTTACATCCATAGTAAATTTGATCGAGAAAATTCCGCATCATTCACAAGTCGTAGAGTTCGGTAGCTCAAAATCGTGTGAGAATGTACTTCGTTTGTTAAATCATAATTCGAGTTACATTCCTTCAGGATCTACAGCATCTGATACTATTAGTCAATTGTTGTGTCAAATAGGA-CA

*Nucleotides CA from exon_2 and T from exon_3 code for amino acid “H”*

>Exon_3

T-TTATACATGACATCTCAGAACTTGAATATTGCTAAACTCTACCCGGAGGTTCAATTTCCAGTGTCTCGGGGAACTCCGATGATTGCGCCGTTGATCCGTTGGGACCACCGAGAAGATGCTTTTGTAGTTAAATATACTTGGGAAGAGAGTTCGAAGTCCAACATGTTGCGATTCAAGATTTCCCTATCCAGTCAAGAATACAAACATATTGTTGGCCATTGCATTGATGGTCGGATATTGTTTCCCGCTACTGGATATTTACAACTGGTTTGGGAGCTTTTGGCTTACATAGGCAACAGGGATCTTGTTGACTATCCAATTGAATTCGAGGACATTCGATATCTGAGAGCCACCACTCTAACGAAAGGACAAACTGTAGAATTGCTGATCACTATTCAGGAGATTTCTGGTCGTTTTGAG

>Exon_4

ATTTCGGAAGGAGATACCGTAGTTGTGACAGGAATCGCTAGAATGTTGGGTGATGCAAATCATCCAAAAATTCAAGAAGTATCGTCTTCAGCTATAACATTGAAAACAAGAGACTTTTATAAAGAGCTTCGTTTACGAGGATACTACTACACAGGTCTATTTAAATCGGTGATGGAGGCGAAAACTGACGGAACCATGGCAAAAATTCAATGGAAGGGTAATTGGATGGCTTTTCTTGATTGTCTGCTACAGACGGGGATTATTGCGATTGACACTAGATCATTGATGGTTCCTACGGCAATTGAAAAGCTTTCCATTGCACCAAAAGCGCATTTGGCAATGATGGAACGCGAGGGAGAAGATTGTGAGTTCTTCACGATGAAGAGCTGTCCTAAAACCAACGTTTTGGTTTGTGGCGGCATTATGCTTTGCAACCCTCGAGCCAGCAGCGTAGGGCGTAGAAACCCTCCAGGTATTCCAGTTCTAGAAACTTATCAATTTGTACCGTATCACACCGATGATCAGGTTTCGACTCTGGAAGCAATTAGAATTTGCGTTCAGCTTGCGCTAGAAAATGTCCCTACTCTATCAATCAATGTGACAGAAATTCACAGCGAGAGGATTCCGGTTGTCGCACATTTGTTTGGAGAAGCAATTGCCGATCTTCCATTGGTTAAAGCCAATCTGATGGTTCTGGCTAAAACAGAAATTGAGTTGGAATATGTGACTGTCAAAGTGGAAAAGCTATCTGATCAATCAAATCTACTGTTTCTGATCACTGACAGCAATTGGAGTGAGCCGAACTTCCTACAAGACGCTGTTGGTCGTCTCGTAGATGGAGGATTCATCATTGTTCGTGAGAAGTTAACCTTCAAGTTAGATGACCTAGATGTACCAGAAGAACTGAACATGGTGGCCTCATTTAGAGTCGATCAAGAGGAGACATTTATTTGTCTGCAACGTAAAATCAAGGGATTCAATGATACTCCGGCAGTTATCCAGGTTGATTCATCCGACCTTAGTTGGCTAGCTGTTTTAAAACAGGCTGTTAAAGTCCGACCTGTGATCCTATTCTCACAGAATGACTCAGTTTCCGGAGTTATCGGTTTGGTGAATTGTATTCGCAAAGAACCCAAAATGCAAACAGTACGATGCGTACTTATTGATGACCATAATGCTCCAGAATTCTCTCTCAGTGATCCTTTCTACAAGAACCAGTTAGAGCTCGGAATGGCAATTAACGTATTGCGAAACGGAGTGTGGGGTAGCTATCGCCATGCCTTAATATCGAAGAAACCAAAAACGGAGCCTGTATCCAAGCATTGCTACGCAAACAGCCTGACAAAAGGTGATTTGTCTTCGATGATGTGGTTCACGGGAGCCTTCAACGAGTGGGATGTTGTACCAAATAAGGTAAAAGTTTCGTACTGCACTTTAAATTTCCGTGATGTGATGGTTGCTACTGGAAGATTGTCATCGGACGTGAGTTCATTCAGCCGACTGGAAGAAGAATGTGAGCTTGGTTATGAATACGCTGGCGTTACGGAAGATGGAAGACGGGTGATAGGAGTCGTGCCTTCAGGAGCTCTATCTACAATGGTTGATGCTGATCCTTCACTCGCTTGGACTATACCAGACAGTTGGAGTTTACAGGATGCTTGTACTATTCCTATTGTATACAATACGGTTTTAACAGCATTTAATATAAGCGCTAATGTGAAGAAAGGGCAGTCGGTTTTGATCCATGCAGGAAGTGGAGGCATCGGATTGGCTGCAATAAATATTGCACTTGCCTATGGAATGGAAGTATTCACTACTGTTGGATCCGACGAGAAGGTCAGTTATCTGTTGAACGAATATCCATCTCTTAAACGAGAAAACATAGGAAATTCGAGAGATTTGTCTTTCGAACAAATGATCAAGCTGAGAACCAATGGAAGAGGGGTAGACTACGTGTTGAACTCGTTAGCTGAAGAAAAACTGCAAGCATCTGTGAGATGCTTAGCGAAGGGTGGACATTTTCTAGAAATCGGAAAATATGATATGGCAAGAGATTCGCAACTGTCATTGGAGCTCTTTAAAAAGGGAATATCATTCACAAGTGTGATGTTGGATTCGGCAATCAGAGATAAACGTAATCTTAAGTTG

>Exon_5 AGCTTACATAAACTATTGAATGACGCAATCAAGTCTGGTATCGTGAAACCACTGAAAACCAACGTTTTCGATGCTGCTGATTTGGAGAAGGCAATGAGATTTTTGGCGAGTGGAAAACATATGGGCAAAATAGTGATCAAAGTACGAGAGAATGAGAACGACGCTGAAACACTTCCAATCACATACTTTCCACACGTTTTCTGTAATCCAGATCAAGTTTACGTAATCGTTGGAGGCTTAGGAGGATTTGGTTTGGAATTGGCGGATTGGCTTATCCTTCGTGGTTGCAGAAAGTTGGTACTCAGCTCTAGTCGAGGTATCACCAAGCCTTATCAAGAATACAGAATT-AA

*Nucleotides AA from exon_5 and A from exon_6 code for amino acid “K”*

>Exon_6

A-ATATGGAATAGTTATGGCGTTCATACTCATATCTGTACTGCCGATGTTACCACAATGGACGGTTGTCGTGTCCTCTTGAAAGAAGCTTCTCGGTTCGGTTCCGTAACGGCTGTCTACAACTTAGCAGTGCAGCTTCGAGATGCCATATTGGAGAATCAAACTGTGGAGAAGTTCGTGGAGTGTATGGCTCCTAAGGCTACGGCAACTGAATACCTTGACAAGGTCAGTCGTGAGATGTGTCCTCATCTGAAGCACTTCATAGTGTTCTCCAGTGTTTCCTGTGGTCGCGGTAACGCAGGACAGAGCAACTATGGTATGGCAAACTCTGTGATGGAGCGTATAATTGAACGGAGAAACACGGACGGCCTTCCCGCAAAGGCGATTCAATGGGGAGCTATCGGTGAGGTTGGACTTGTAGCAGATATGGCAGAAGATAAGATCGATTTGGAGATCGGAGGAACGTTGCAACAGCGCATATCATCATGTCTTCATGAGATGGATTACTTGCTGACATGTGAAGCTCCTCTTGTGGCCAGTATGGTAGTGGCAGAAAAACGAACTGCAAGCGGATCAAAGAATGTCATCGAAGCTGTCATGAACATAATGAGTATAAGAGATCTGAAATCGGTGTCGATGGAGAGTACTTTGGCCGATATTGGAATGGATTCTTTGATGGCAGTAGAAATCAAACAGGTACTGGAAAGAGACTTCGACATGGTATTGTCACCGCAAGATTTGAGAACATTATCCTTTGCCAAGTTGTTGAAAATGGATGAGGAGAAAAAGCAAGCTGCAAAAGATCAGGAGGAAAAGAAGAGTGAAGGCTTTGTGATTGGAATGCAAATGCTGTTGAGAAATCTTGGCAACGAAGAAACGAGTGAATCGACACTGTTGCATTTACCTTCTGCTGGCCAAGAAGGCCGTCCACTATTGCTCATTCCAGGTGTTGAAGGTGTGGCCGGTAATGTTTGGAAAGCTATTGCTGCACAGATAAAGTCTCCTGTCTACATGCTCCAGCTATCGAGCACTTTAGATTGTGATAGCATCCCAGATATTATAGAACGTGTCATAGATGAGATCTGTGAAACAATGTTCAATGGTTTTGAAGATATTACAATAGTTGCTTATTCATTTGGAGCTCTGATTGCAATCGAAATAGCTCGATATTTACAGGCAAAAGGTATTCGCGGAGAACTTCTACTATTGGATGGTGCACCGAAGTACTTGAAGCAATGGTCACTGAAACAGCTGAATAATAATCCATCGGATGGAGAAATACAGAAGCTCATATTACTTGTCTTGATCGCCATGGTGTTCCCAGATCAGCCACCCGAGAAAGCCATGGCCATATTGGAGATTACGTCATTCGACGACCAAATTGAGAAACTCATTGAATTGGGAGCAGAGCAGAGCGAATATTCCCCAGAGTACACAAGAAAGATGACGAAAGCCCTTTGCAGGAGAATAAAAATGGCAGCTCTGATGTACCTGGACGAAGATCAACCGTTAGACCTTCCAATAACGCTGGTGCGGCCAACCGATGCCGTTTTCTCGGATATTGAGGATGACTACGGGCTTTCTAGTTATACCACAGGAGCTATAACACTGCGAATGGTCGAAGGCAATCATGTATCTATGTTGGAAAATGCTGATCTTGTGGAAATGATCAATAACTTTTGTGTTTAGTTACCGTTAGAATATTTTGAATTCGTCATTAATATTGTTATCTTAAATAAAGCACATTTATGTTTACCGATCTTTA

**LOC5573929_amino_acid_sequence**

>Exon_1

MPTRCIQSVGTDKSIVISGISGRFPRANNVREFASSLYGKQDLVDDLETRWQHTMQDVPRRTGKVGNMQNFDADFFGVSRIERDTMDPQLRMTIEHVYEAILDAGVNPQTLRGSRTGVFSGVCFSETEVCMYYRARPPKGLGLL-G

>Exon_2

CAKSQIPNRVSYLLDLRGPSYVLDTACSSSMYALDVAYRSMMNGECDAAIVTGANLTLHPFITYQFAMLGVLAKDGYCRPFDKDATGYTRSEAVCAVFLQKAKDAKRVYGHIIHSKTNCDGFKPEGITYPSGSVQQQLLTEFYNEVGISPTEVDYVEAHSTGTFVGDPEECDAIDKVYCSGRTDPLLVGSVKSSIGHTEASAGVCSITKCIIAMENGLIPPNINYTDYRPTIPSLVEGRLKVVTDAMPLSGPLVGINSFGFGGANAHALLCRNLKEKVKNGVPDDDLPRLVTWSGRTRESIETMLHDIGQRPLDVEFIALLFNIQQQPTPGHRYRGFGIYQKNGDRPAVLQTSSIDRVKLDAIPVVAVFGGINTNWRKELDALRQFSVVEDTFAKCSGILRSLKFDLHKKPSGRESILYNMVGATILQLSIVDLLSSIGVKFDFYGGHSIGQFTCAYIDHNLNLEQVLRLAFWHGLVFSDCHAVCDRTAFVQINSMLNQLPLKNFFKDSATTFGILTANEKILMEQVRQLKSSGFAAEELSFLDVHADSTKSSSLANKLRQTVNTVLSRIILPSDKWITSALPHTSSIFHSSKLHDVTSIVNLIEKIPHHSQVVEFGSSKSCENVLRLLNHNSSYIPSGSTASDTISQLLCQIG-H

>Exon_3

LYMTSQNLNIAKLYPEVQFPVSRGTPMIAPLIRWDHREDAFVVKYTWEESSKSNMLRFKISLSSQEYKHIVGHCIDGRILFPATGYLQLVWELLAYIGNRDLVDYPIEFEDIRYLRATTLTKGQTVELLITIQEISGRFE

>Exon_4

ISEGDTVVVTGIARMLGDANHPKIQEVSSSAITLKTRDFYKELRLRGYYYTGLFKSVMEAKTDGTMAKIQWKGNWMAFLDCLLQTGIIAIDTRSLMVPTAIEKLSIAPKAHLAMMEREGEDCEFFTMKSCPKTNVLVCGGIMLCNPRASSVGRRNPPGIPVLETYQFVPYHTDDQVSTLEAIRICVQLALENVPTLSINVTEIHSERIPVVAHLFGEAIADLPLVKANLMVLAKTEIELEYVTVKVEKLSDQSNLLFLITDSNWSEPNFLQDAVGRLVDGGFIIVREKLTFKLDDLDVPEELNMVASFRVDQEETFICLQRKIKGFNDTPAVIQVDSSDLSWLAVLKQAVKVRPVILFSQNDSVSGVIGLVNCIRKEPKMQTVRCVLIDDHNAPEFSLSDPFYKNQLELGMAINVLRNGVWGSYRHALISKKPKTEPVSKHCYANSLTKGDLSSMMWFTGAFNEWDVVPNKVKVSYCTLNFRDVMVATGRLSSDVSSFSRLEEECELGYEYAGVTEDGRRVIGVVPSGALSTMVDADPSLAWTIPDSWSLQDACTIPIVYNTVLTAFNISANVKKGQSVLIHAGSGGIGLAAINIALAYGMEVFTTVGSDEKVSYLLNEYPSLKRENIGNSRDLSFEQMIKLRTNGRGVDYVLNSLAEEKLQASVRCLAKGGHFLEIGKYDMARDSQLSLELFKKGISFTSVMLDSAIRDKRNLKL

>Exon_5

SLHKLLNDAIKSGIVKPLKTNVFDAADLEKAMRFLASGKHMGKIVIKVRENENDAETLPITYFPHVFCNPDQVYVIVGGLGGFGLELADWLILRGCRKLVLSSSRGITKPYQEYRI-K

>Exon_6

IWNSYGVHTHICTADVTTMDGCRVLLKEASRFGSVTAVYNLAVQLRDAILENQTVEKFVECMAPKATATEYLDKVSREMCPHLKHFIVFSSVSCGRGNAGQSNYGMANSVMERIIERRNTDGLPAKAIQWGAIGEVGLVADMAEDKIDLEIGGTLQQRISSCLHEMDYLLTCEAPLVASMVVAEKRTASGSKNVIEAVMNIMSIRDLKSVSMESTLADIGMDSLMAVEIKQVLERDFDMVLSPQDLRTLSFAKLLKMDEEKKQAAKDQEEKKSEGFVIGMQMLLRNLGNEETSESTLLHLPSAGQEGRPLLLIPGVEGVAGNVWKAIAAQIKSPVYMLQLSSTLDCDSIPDIIERVIDEICETMFNGFEDITIVAYSFGALIAIEIARYLQAKGIRGELLLLDGAPKYLKQWSLKQLNNNPSDGEIQKLILLVLIAMVFPDQPPEKAMAILEITSFDDQIEKLIELGAEQSEYSPEYTRKMTKALCRRIKMAALMYLDEDQPLDLPITLVRPTDAVFSDIEDDYGLSSYTTGAITLRMVEGNHVSMLENADLVEMINNFCV

**AaFAS4 LOC5573931 (AAEL002237)**

**XM_001654917.2 PREDICTED: Aedes aegypti fatty acid synthase (LOC5573931), mRNA**

>Exon_1

ATGTCGGCCGTCAACTTGGGTTCGGTTCAGCTAGATGAGTGCGTAGTTTTGTCCGGCATTTCGGGCCGCTTCCCGCAGTCGGATAATATGCGTCAGTTTGCTGAGAACCTGTACGACAAGCGTGATCTGGTGGATGATAAGGAAACACGTTGGCCGCATACTATGCCGGAGATTCCGCGTCGTTCCGGTAAGATCAACAATTTGGACAAGTTTGACCGGGAGTTTTTCGGTGTCAGCCGGAATCAGTGCAATGCTATGGATTCGCAATTGCGGCTGTTGCTGGAGCATGCTTATGAAGCGATTGTGGACTCCGGTACCAACCCCGAAACAGTGCGGGGCTCGAAAACCGGAGTGTTTGTCGGAGTTTGCTTCTCGGAAACTGAGGCTAGGTTATTTTTTCAATCGTGCCCTCCGAAAGGATATGCTATGCTA-GG

*Nucleotides GG from exon_1 and A from exon_2 code for amino acid “G”*

>Exon_2

A-TGTGCCAAGTCTCAGATAGCCAACAGAATTTCCTATGTGTTTGACCTTCGTGGACCGTCTCTTGTACTGGATACTGCGTGCAGCAGCAGCATGTATGCTATGGATGTAGCCCGTAGGAAAATATTGAGTGGAGAATGTGATGCTGCTCTTGTTTTGGGAACTAATCTTTGCTTGCATCCATACATTTCGTATCAGTTCTCTCTGCTGGGCGTTCTAGCTAAAGACGGTGTTTGTCGACCGTTCGACGAAAAGGCCAACGGGTATACACGATCTGAAGTGATTTGTGCTGCGTTTTTACAAAAAGCTAAGGACGCCAATCGAATCTATGCGCACATCCTACACTCTAAGACCAATTGCGACGGATTCAAATCCGAGGGAATAACATATCCCTCAGGAGTAATGCAGAAACGGCTCATGAGTGAATTCTACGATGAAATATCAATTGATCCCAGAGACATTGCCTACATAGAAGCTCATAGTACCGGTACTGTTGTAGGAGATCCGGAAGAATGCGATGCAATTGATAAGGTGTTTTGTACGGATCGCCAACGTCCTATGCCGATCGGGTCAGTTAAATCCAATATTGGACACTCGGAAGCCTCAGCGGCATTGTCATCACTTGCCAAGTGTGTCATTGCAATGGAAAGTGACTTGATTCCTCCTAATATCAACTTCACTAAAAATAGAGCTGACGTTCCAGCCTTGAACGCGGGTCGTCTACAGGTCGTCGATCATGCACAACCAATGGAAGGTCCACTTATAGCGATCAACTCTTTTGGATTTGGGGGAGCTAACGCGCACATGTTAATTCAACGTAACACAAGGGAGAAAACTGGACAAGGAATTCCAGATGATGACCTTCCACGTCTAATTACCTGGTCGGGTAGAACCAAAGAAGCAGTGGAATATATGTTTCAGGACATATGCACTCGTCCGCTGGACGTCGATTTCGTCGCTTTGTTGCATAATATGCAACGCACTCGAACTCCAGGACATCGATATCGAGGCTTCGCTGTTTTCGAAAACCGAGGATCACAGCCTACCAGCATGAACGTCTTCAACATAAATCGCGTCAAACTCGACGATTCGCCACTTGTGGCTATCTTTCCCGGTATAACTTTGAGGTGGCGAGAAGATTTGGAAGCCTTAAGTCAACTTCCATGTGTTCAGCAAACGGTGACCGAGTGTTGTACTGTACTGGGAACTTTTGGGTACGATCTTTTCCATAAACCATCGCGGAGAACGGATCTTCGACAGCTCTTGATTGGATCCACTGTACTCCAGCTGGCGTACGCTGATATTCTTACGACAACTGGAGTCCAGCTAAACGCATACGGAGGTCACTCGATCGGGCAGTTTACTTGCGCTTATCTCGATGGCTGCCTAACTTTGGAACAGATAGTCCAAATTGCGAACTATCACGGATCTGTTTTAGCCGAGTACAAAACCGAAATGAACTTCAACGCATTCCTCGAATTAGGCTCGAAACGTCCGGTACCGCCACTTTCTTTCGATGGGTTCATTCAAGACACATTTCTGTCGCGTTTCGGAGTCGTTGGAGGTCCGCTGAATCCCACCTGTGCCGTGGTTGATCAGTTGAAGTCACGAGGCTACGTCGCAGAGCATTTGCCATTTGTGAGTCTGGTCTACGCAAAGGATTCGACAACTGAGCTGTCGGAGAAGCTGAATACCCTTGTTTCGCACATAATTCCACAGCCTCTGACACCTTCGCTTCGCTGGATCAACCTTAAAAACCCATCGGACTTCGCATCGACTTCGGTTCACGATTCGTTCTCTATTGTAAACCTGTTGGAGAAAATCCCGGAGCATTGCATGTTACTGGAACCATTAGCAAAACAATCTATGCAACCTGTTCTGAAAAGTTTGAATCGAAAATCCAAATGCATGCCTTTTGAGAAGGGTACTTCAACTATTAAAGGGCTTCTATCTACGCTAGGA-CA

*Nucleotides CA from exon_2 and C from exon_3 code for amino acid “H”*

>Exon_3

C-CTGTACTTAACTCAGCAGGATCTGAATTTGCTGAACCTCTACCCGCCGGTACAATTTCCAGTTGCTCGAGGAACTCCAATGATTTCGCCGTTGATACGATGGGATCATCGTGATTCGTGGTATGTCGTGCGGTACGAATGGGTGGCGTTCCGAGTATCCAACCAGTTGAACTTCAAGGTTACCCTAGCGGATCAGGACTTCGTTTATGCTGCTGGACATTGCATTGATGGAAGGGTGCTATTTCCGGCAACGGGTTACCTCAGTCTGGTATGGGAGCTGATAGCATACCTGAAGCAACGGGAGCTGTCGGATTGTCCGGTGCAGTTCGATGATGTGCAGTTTCTACGAGCGACAACACTCACCAAGAATAAGGCTGTCAATCTCTTGGTCACATTGCAAAAGGGTACTGGAAGATTCGAG

>Exon_4

ATTGCAGATGGTACAACTCCGGTGGTGACTGGGTTTGCGAGAATGCTCGAAACAAAGCACGAGCAGTATCAGGCTTTCGAAGAGAACTCGTTAGCTCCCCTGCTGACATCCCGGGACTTCTACAAGGAGCTCAGGCTGCGGGGATATCATTACAACGCGTTGTTCAAATCCGTAATGGAAGCTCACAGCGACGGTTCCGCGGCTAAGATTCAATGGAAGGGTAACTGGGTTGCCTTTTTGGACTGCCTCTTACAGGTAGGAATAGTTTCTATCGATACCCGATCCTTGATGGTTCCGACAGCCATTGAAAGTGTCACGATCTACCCGAAACAACACCTGTCGTTGATGTCTCGTGATGATGAAGGTTCTACAGACTACTTCAACGTTTCGAATTGCGCCAAGACTAACGTATGCGTCTCCGGAGGCGTTCGAATAACTGGCCTTCGGGCAAACGTCGTAAGTAGACGAAACCCTCCGGGTGTTCCTGTACTGGAGACATATAGCTTTGTGCCATATAATGCTACCGAGGAAGTGTCTCTCACGGAAGCAGTGAGGATGTGCGTTCAGATAGCTTTGGAAAACGCACCAACTCTGTCAATTCGTGCTACGGAGATATATAACGATGGTACTGAGTTATTATTGCCATACTTTGCAGAGGCCGTGGCTGATTTGCCATTGGTTCAACCAACCTTGACTTTGTTGACGTCGAAAGACATGGAACTTTCAAATATCACAGTCAAAAATGAGAAACTTGCGGACCAAACGGGACTTCAGTTCATAATTAGCCACAACGTGCTAAACGATGACAATTTTCTACGAGACGCTATGACTAGTCTGGACCCTTCAGGTTTCCTAGTAATACGGCACTCGAATACCGATGTTGATATTCCGGCAAAGCTGGTGAAAGTCGCTAGCTTTATCAATCAGAAGAACCAAGCACTTATTCTTCTGCAGAAGACATCCAACAATTTCAAAGAATCACCGGCCGTTATCCGGGTTCGTTCAGACGATCTTAATTTTGAATGGCTTCATGAACTGCAGAACGTGATCAAAATTAAACCGGTTCTGCTGTTTTCGCAAGATGATCCAATCTCTGGCATAATCGGCCTGGTCAACTGCATCAAGAAAGAGCTTAAATCATACCCGGTGCGCTGTGTTTTCATAGACGACCGTAGTGCTCCACCGTTCGCTCAGAGTGAACCGTTCTACAAGAATCAACTCAGCAAGGATCTAACCATCAACGTCTATCGAAATGGAACTTGGGGTAGCTACCGTCATGCGCTCATGGACCTTAAACCGAAAGTAGAGTCCGTACGCAATCATTGCTTTGCCAATTGCTTCACGAAGGGTGATCTTTCGTCGATGACCTGGTTCAGCGGTCCGTTGAACACTTTTTCTTCCGGTGGCGAACTGATACGGGTTGTGTACAGTGCGTTGAACTTCCGCGATGTGATGATCGCTACGAGTAGACTTTCGTCGGATGTGCTGCACGTCAATCGGTTGGAGCAGGAGTGCTTGCTTGGCAATGAATATTCTGGAGTGTCGGTGCGTGGTCGACGGGTTATGGGAGTTCTGCCCAGTGGGGCTATGGCAACCTTGGTAGAGTGCGATCCGCTGATGACTTGGACCATTCCAGATGACTGGAGTTTGGAAGAAGCGGCAACTGTTCCCGTGGTTTACGGAACAGTTTACACGGCACTGTTCGTATGCAGTCGAATCAGGAAGGGCAAGTCGATCTTGATCCATGCTGGCAGCGGAGGCGTTGGATTAGCAGCTATTCAAGTATGCCTGGCATACGGAATGGAAGTGTTCAGTACGGTTAGCAATGAGGAGAAGAAGCAGTTTATTTTGAAACGCTATCCACGCTTAAAGGCAGAGAACATAGGAAACTCCAGGGACACTTCTTTCGAGAACATGATCAAATTGAGAACGAGTGGCCGAGGGGTAGATTTCGTTCTGAACTCTTTGGCTGAAGAAAAGCTACAAGCATCGCTGCGATGTCTCAGTAGAAACGGTCACTTCCTGGAAATAGGAAAGTACGATATGGCAAACAACTCTAAGATCGCCATGGAGCTGTTCCAAAAAGGAATTTCCTTCACGGCTGTTATGTTGGATGCCATGTTTAAGGGATCTTTTGCAGAAAAACAG

>Exon_5

AGACTACATGCATTACTAACGGCAGACATGCTGAGTGGAGTCATCAAGCCATTGAACACCACGGTCTTCCCGGCGATAGAAATAGAAAAGGCCTTCCGGTTTCTTGCCAGCGGGAAACACATTGGCAAAATTCTTCTGGAGATCCGTAGCCGCGAAGACGATGCGCAGACAATGCCTATCAACTATGTTCCAAGAATGTACTGCGATCCAGACTGCTCATACGTGATCGTAGGAGGACTGGGAGGCTTCGGTTTGGAACTGGCAGATTGGTTAATTCTACGTGGATGTCGCAAGCTTATTATCAGTTCTAGCCGAGGAATCACAAAACCCTATCAGGAATATCGAATT-AG

*Nucleotides AG from exon_5 and A from exon_6 code for amino acid “R”*

>Exon_6

A-TTGTGGAATCGATACGGAGTTCGTACTTTGGTGAATACTGCCAATATAACCACCCTGGAAGGATGTCGTGAGCTTCTCCGTGAGGCATCTGTTATAGGGCCAGTTATCGCCATCTTTAATCTAGCGGTCCAATTGCGGGACTCCATCCTGGAAAACCAAACCCCCGGAAAGTTCAGCGAATGTCTAGCACCGAAAGCGTATGCTACCCGCTTTCTGGACGAAGTTAGTCGAAAGTTGTGCCCAAGGTTAACGCACTTCGTGGTGTTCTCAAGTGTATCATGCGGGCGAGGAAACGCAGGCCAGAGCAACTATGGAATGGCCAATTCCATGATGGAGCGGATTGTGGAGAACCGTGTAGCGGAAGGCTTGCCGGGGAAAGCCATTCAATGGGGAGCTATCGGCGAGGTCGGGATTGTTGCAGACATGGCCGAGGATAGACTCGACATGGAGATTGGCGGTACACTACAGCAGCGGATTTCGTCTTGTTTACAGGAACTGGACCGTCTGCTGACGTGTGACGATCCCATTGTGGGCAGTATGGTTGTAGCGGAAAAGCGCATCTGCGGAGGTTCGAGGAACATAATGGAAACAGTTATGAACATCATGAGTATTCGTGATATGAAATCAGTTTCCATGGAAAGTACCCTAGCGGATGTTGGTATGGACTCACTGATGGCAGTAGAAATTAAACAAGTTTTGGAGAGTGACTTCGATTTGGTTCTATCGCCTTTGGAGCTTCGCACGTTAACCTTCATGAAGCTACAGAAGATGGTAGATGAAAAGAGACTCAATGCAGACGCAAAGACCGAAATGTCTGCCCGTCCAATGGTCATCGGTATGGAAATGCTGCTCAGAAACTTGGGTGAAGAGGAGCACAGTGCCATAACGTTACTCCCCTTGCCATCGAAGTCAACTGAAGGACGTCCAATTCTTATCATTCCTGGCTTGGAGGGTGTAGCAGGTCATGTTTGGCGAATATTGGCGGCCGGACTAAATGCGCCTGTTTATATGCTACAAACATTGAACACGTTTGAGCTTAAAAGCATTTCGGAAATTGTGGATCACGTTTTTGATGAGCTAGACAAAGAAGTATTCGCCGGTTTTGATGACTTCTTCATTGTTGGCTACTCCTTCGGAACTTTGATCGCTTATGAAATCGTAAGACGACTGCAGCGCCGGAACCTCCCCGGAAAACTAATACTGCTTGACGGAGCTCCGAAGTTCTTGAAGCAGCTGTCAGCGATGCAGATGGGTAACGGCGTGTCGGACGAGGACATTCAAACCATGCTTATGATTGCAATCATATCGCAAGTGTTCCCGAGTTCTCCCATCGAGAAGAGTTATTCGATTATGCAGGTCGCATCCTTTGATGGGCAAATCGAGAGGCTGATCGAAGTGGCCAAGGGCCAGGCAACTTATTCGCCGGACTATACTCGCAAAATGACCAAGGCACTGTTCAACCGTATCAAGATGGTGCTTGGTTTGAACATTGAGGCAATGGAACCTCTGAAGGTGCCGATTGTGCTGGTGCGCCCGTCGGAAGTGTCTGTGATCGATATAGACGAAGAATATGGATTGGAAGGAAGTACCACTGCAACAATGTCGATGAAAGTAGTTGATGGAAGCCACATGACGATGTTAGACAACAAAGTTTTAGTGGATATTATAAATAGTTTGAATGTGTAGAATGCACAAACTACGTGAGGAAATGAAACTGGCATGACCGGACTACGAACAATATTTTTAGATTAA

**LOC5573931_amino_acid_sequence**

>Exon_1

MSAVNLGSVQLDECVVLSGISGRFPQSDNMRQFAENLYDKRDLVDDKETRWPHTMPEIPRRSGKINNLDKFDREFFGVSRNQCNAMDSQLRLLLEHAYEAIVDSGTNPETVRGSKTGVFVGVCFSETEARLFFQSCPPKGYAML-G

>Exon_2

CAKSQIANRISYVFDLRGPSLVLDTACSSSMYAMDVARRKILSGECDAALVLGTNLCLHPYISYQFSLLGVLAKDGVCRPFDEKANGYTRSEVICAAFLQKAKDANRIYAHILHSKTNCDGFKSEGITYPSGVMQKRLMSEFYDEISIDPRDIAYIEAHSTGTVVGDPEECDAIDKVFCTDRQRPMPIGSVKSNIGHSEASAALSSLAKCVIAMESDLIPPNINFTKNRADVPALNAGRLQVVDHAQPMEGPLIAINSFGFGGANAHMLIQRNTREKTGQGIPDDDLPRLITWSGRTKEAVEYMFQDICTRPLDVDFVALLHNMQRTRTPGHRYRGFAVFENRGSQPTSMNVFNINRVKLDDSPLVAIFPGITLRWREDLEALSQLPCVQQTVTECCTVLGTFGYDLFHKPSRRTDLRQLLIGSTVLQLAYADILTTTGVQLNAYGGHSIGQFTCAYLDGCLTLEQIVQIANYHGSVLAEYKTEMNFNAFLELGSKRPVPPLSFDGFIQDTFLSRFGVVGGPLNPTCAVVDQLKSRGYVAEHLPFVSLVYAKDSTTELSEKLNTLVSHIIPQPLTPSLRWINLKNPSDFASTSVHDSFSIVNLLEKIPEHCMLLEPLAKQSMQPVLKSLNRKSKCMPFEKGTSTIKGLLSTLG-H

>Exon_3

LYLTQQDLNLLNLYPPVQFPVARGTPMISPLIRWDHRDSWYVVRYEWVAFRVSNQLNFKVTLADQDFVYAAGHCIDGRVLFPATGYLSLVWELIAYLKQRELSDCPVQFDDVQFLRATTLTKNKAVNLLVTLQKGTGRFE

>Exon_4

IADGTTPVVTGFARMLETKHEQYQAFEENSLAPLLTSRDFYKELRLRGYHYNALFKSVMEAHSDGSAAKIQWKGNWVAFLDCLLQVGIVSIDTRSLMVPTAIESVTIYPKQHLSLMSRDDEGSTDYFNVSNCAKTNVCVSGGVRITGLRANVVSRRNPPGVPVLETYSFVPYNATEEVSLTEAVRMCVQIALENAPTLSIRATEIYNDGTELLLPYFAEAVADLPLVQPTLTLLTSKDMELSNITVKNEKLADQTGLQFIISHNVLNDDNFLRDAMTSLDPSGFLVIRHSNTDVDIPAKLVKVASFINQKNQALILLQKTSNNFKESPAVIRVRSDDLNFEWLHELQNVIKIKPVLLFSQDDPISGIIGLVNCIKKELKSYPVRCVFIDDRSAPPFAQSEPFYKNQLSKDLTINVYRNGTWGSYRHALMDLKPKVESVRNHCFANCFTKGDLSSMTWFSGPLNTFSSGGELIRVVYSALNFRDVMIATSRLSSDVLHVNRLEQECLLGNEYSGVSVRGRRVMGVLPSGAMATLVECDPLMTWTIPDDWSLEEAATVPVVYGTVYTALFVCSRIRKGKSILIHAGSGGVGLAAIQVCLAYGMEVFSTVSNEEKKQFILKRYPRLKAENIGNSRDTSFENMIKLRTSGRGVDFVLNSLAEEKLQASLRCLSRNGHFLEIGKYDMANNSKIAMELFQKGISFTAVMLDAMFKGSFAEKQ

>Exon_5

RLHALLTADMLSGVIKPLNTTVFPAIEIEKAFRFLASGKHIGKILLEIRSREDDAQTMPINYVPRMYCDPDCSYVIVGGLGGFGLELADWLILRGCRKLIISSSRGITKPYQEYRI-R

>Exon_6

LWNRYGVRTLVNTANITTLEGCRELLREASVIGPVIAIFNLAVQLRDSILENQTPGKFSECLAPKAYATRFLDEVSRKLCPRLTHFVVFSSVSCGRGNAGQSNYGMANSMMERIVENRVAEGLPGKAIQWGAIGEVGIVADMAEDRLDMEIGGTLQQRISSCLQELDRLLTCDDPIVGSMVVAEKRICGGSRNIMETVMNIMSIRDMKSVSMESTLADVGMDSLMAVEIKQVLESDFDLVLSPLELRTLTFMKLQKMVDEKRLNADAKTEMSARPMVIGMEMLLRNLGEEEHSAITLLPLPSKSTEGRPILIIPGLEGVAGHVWRILAAGLNAPVYMLQTLNTFELKSISEIVDHVFDELDKEVFAGFDDFFIVGYSFGTLIAYEIVRRLQRRNLPGKLILLDGAPKFLKQLSAMQMGNGVSDEDIQTMLMIAIISQVFPSSPIEKSYSIMQVASFDGQIERLIEVAKGQATYSPDYTRKMTKALFNRIKMVLGLNIEAMEPLKVPIVLVRPSEVSVIDIDEEYGLEGSTTATMSMKVVDGSHMTMLDNKVLVDIINSLNV

**AaFAS5 LOC5573927 (AAEL002228)**

**XM_001654913.2 PREDICTED: Aedes aegypti fatty acid synthase (LOC5573927), mRNA**

>Exon_1

GGTCGTTTGTAGTGAATCCCTGTGCTAGAATACCGGTCCAAAATGCCATCGCCAAACATTCAATCGGTTGGGTCGGATGAAAGTGTGGTAATATCGGGCATAGCTGGACGTTTTCCGCAGTCGGATGACGTGCGCGAGTTTGCGCGCAATATGTACGAGAAGCGTGATCTGGTAGACGACAAGGAGACTCGGTTTACCCACGCTATGCAGGCCATTCCAAGGCGCTTCGGGAAGATTAACAATCTGCAGAAATTCGATAGGCAATTTTTCGGTTATGATCGTCGGCAGCGGGATACGATGGATCCCCAGCAGTGGATGGCTCTGGAGCATGCTTTCGAGGCTATTTTGGACGCAGGAATCAACCCGGAAAGCTTGCGAGGCACTCGGACGGGTGTTTTCTGCGGAGTTTGTTTCTCGGAGTCTGGAGCACGAATGTACTACAGTGCGCTGGGAAATGCCGAAAAAGGGATGGGACTGCAA-GC

*Nucleotides GC from exon_1 and C from exon_2 code for amino acid “A”*

>Exon_2

C-TTCAACAAGTCATACTTGGCGCTTAGAGTGTCTTACACGTTGGATTTGAACGGACCGAGTGTTGTGGTTGATACGGCCTGCAGCAGTTCGATGTACGCTTTAGACGTTGCTTATCGAAGCATCTTGAATGGTGATTGCGAGGCAGCAATAGTAACTGGGTCAAACTTGACACTGCATCCGTTCATCTCGTATCAGTTTGCACTTTTGGGTGTGTTGTCCAAGGATGGATACTGTCGGCCGTTTGATAAGGAGGCTTCCGGGTACACGAGATCGGAGGCAGTTTGTGCGATATTCCTTCAGAAAAGTAAGAACGCCAAGCGAATATATTGCCATGTTGTGCACTCCAAGACGAACTGTGATGGATTCAAAGCGGAAGGGATAACCTATCCGTCGGGATTGGTTCAGCAACAGTTGTTACGAGAAATCTACGATGAAGTGGGGATCAGTCCGATTCAGGTGGACTACGTTGAAGCTCATAGCACTGGTACTGTAGTGGGTGATCCGGAAGAATGCGATGCAATTGATAAGGTTTATTGTAAGGGACGAGATACAGCTTTGCTCGTGGGGTCGGTTAAGTCCAGCATAGGGCATACCGAAGCAGCTGCGGGATTGTGCTCGATAGCGAAATGTATCATTGCGATGGAATACGACTCGATTCCCCCAAATATAAATTACACTGAGTGTAGAAAAGATATCCCCGCATTGGTTGAAGGCCGATTGAAGGTCGTCACGGAGGCTACGCCTTTACCCGGGCCATTGATTGGGATTAACTCTTTCGGGTTTGGCGGAGCTAACGCCCATACTTTGTTGTGTCGCAACTTGAAGGAGAAGAAGAATAACGGTGTGCCGGAAGACGACCTACCGAGGATAGTTTTGTGGTCCGGAAGTACCCGAGAAGCTGTTGAATGTATGCTTCAGGATATCAGTCAACGACCGTTGGACGCTGAATTCATTGGGCTGATGCACAATATCGAAAAGCAACCGATACAAGGTCATCGTTACCGTGGCTTTGGGGTATACCAGAAAGATGGCGAAAGAGCAAGTATTCTGAAAGAATCGTGTATCGAGCGAGTTAAACTTGAATCCCTTCCAACCGTTGCTATCTTCGGGGGAATTAATGTCGGCTGGAAGCAGGATTTGGACAGTCTGAGAAAGATTTCGTCGGTAGATGCGTTTTACACAATGTGTAAGGAGGTACTCAAAACATTAAAATTTGATCTATACAAGGCCCATCCACACGAGAAGGATCTTCTCTACAATATGGTTGGTTCAACGGTACTGCAATTATCAATCATTGATTTGCTGGTCTCGAGTGGAGTTGCCTTAGATTTCTACGGGGGTCATTCAGTCGGACAGTTCATTTGTGCCTACATTGACCGCTGTTTGACCCTGGAACAAGTTCTGAAGCTCGCTTTCTGGCATGGTCTGGTTTATTCCGACTGCCACGCTCTCTGCGAAAGTTCGGCATTTGTTCGAATGAACGCCAAATTGAATCAACTACCACTGAAAAATATTATAAAAAATTCTGCATCGTCATTCGGTATTCTAACATCCTCGGAGAGAAACATAATCGAGCAGGTTCATCAGCTGAAGTCATCTGGATTTTTGGCGGAAGAGTTATCGTTCTTATGTTTCCACTCGGCGCCCGCTTGCTCTGGAACTTTGAGTAACACACTGCGTCAAACCGTGAATGTCGTTTTATCTAAAATCATTGAACCTTCAAACAAGTGGATCACAGCAAGACTACCCCAAGGATGCTCAATTTTCCACTCACCGGCACTGCATGATGCATCATCGATCGTTTGTATGCTCGACAAGATTCCCAGTCACACGAACGTTTTGCAACTGGATAGCGCGCAGACTTGCGAGAAAGTACTCCAGATGATGCATAGACACTCTCGTTGCTTAACGCCCAGTTCGGAGTCCTCCGATGCCATTTGTCGACTCCTAAGTGTGATTGGC-CA

*Nucleotides CA from exon_2 and A from exon_3 code for amino acid “Q”*

>Exon_3

A-TTACATTTGATTTCCCAAAACTTGGACGTCGCTCATCTGTATCCTCCGGTGCAGTTCCCAGTTTCACGAGGAACGCCAATGATTGCGCCTTTGGTCCACTGGGATCATCGTGAAGACGCCCATATCGTGAAATACACCTGGGATCAAACGGCCAAGTCCCGAGTTGGCACCGTCAACATTTCCGTGTCCAGTCAGGACTGGAAGCACATCGAGGGTCACAACATTGGCGGTAGGGTGCTGTTTCCGGCTACGGGATACCTCAAACTGGTATGGGATCACGTTGCATACCTTGCGCACTGCGATTTGGAGGACTTCGCGATAGAGTTCGAAGATGTGAGGTTCTTGCGGGCAACTAATATCACCAAGGGGCAAACCATTCGGTTCAGGGTTGCCGTAGAGGAGGTATCTGGATTTTTCGAG

>Exon_4

GTGATAGAAGGTGACACGATAGTTGTTACTGGATTTGCTCGACAATGGAACGATAGCGTCCAGGAGAGCTTCAGGGAAGAAAGGAGTTCAGTGGCAGTACTTCGTACGAAAGATTTCTATAAAGAGCTCCGTTTGAGAGGATACTATTATTCGGATCAGTTCAAATCGGTTCTGGAGTCTAAAGCAGACGGATCGCTCGCGAAAATACAGTGGAAAGGCAACTGGGTACCGTTCTTGGATTGCTTGCTGCAAGTTGGAATTATAGCAACTGATAGTCGGCAACTGATGGTACCAACAGCTATTGAGAAGCTGTCTGTGTCGCCGAGAGCTCACATGGAAATGATCGAAAGAAATGAGGACGGCACTGAGTTCTTCACGATGAGGACTTGCTTGCTCACCAATACGTCGATTTGTGGATCGGTGATGATTCGTAATTTGCGAGCTAGCAGCGTGAATCGTAGGAATCCCCCGGGAAATCCAGTCTTGGAGATTTATCAATTCGTTCCTTTCCATTCGGATCATTACATTCCAACAGTGGAAGCCATTAGAATCATCGTCCAGTTGGCTCTGGAGAATGCTCCAACCTTCACTGTTAACGTTACAGAAATCCACCACAAAGATTCTCAGCCTATAGTGCAATTGTTCGGAGATGCTGTGGCTGATTTACCGTTAGTCCAATCCAATCTGACCCTTCTAAGCACTAAAAGTATTGAGTTGGACGACGTCACCGTTCAAGACGAGAAACTTTCAGGCCTAACAAACCAACTGTTTTTGATCACTAATAGCAAATTGGAAGATAACACATTCATTCAGGAAGCTGTTGGATGTTTAGTTGATCGCGGGTTTATAATTGCCCGGAGCACCAATCCTGCAGAGCTAACTGTACCAACCCAGTTCACTTTAGTGGCTTCGTTCCGCGTTGATCAATCCGAAACCCTCATCTGCTTACAGCGAAAAGCCTTCGCAGAATCTCCAATAGTTGTCAAGCTAGATTCAGTTGAAATGAGCTGGCTAACGAATCTCAAACGAGCAGTCAAAAGCCGATCGGTGCTTCTGTATGCACAGAATGACCCGACTTCTGGAATCATCGGTCTGGTGAATTGCATCCGGAAGGAGCCCAAAATGCAAGATGTACGGTGCGTGTTCATCGACGATCCGAAGGCCCCGCCCTTCATGATCAGCGACCCTTTTTACAAAAATCAGCTGGACCTGGGTTTGGCGATCAACGTGCTGCGTAACGGAGTCTGGGGTAGTTATCGTCACCAGTTGTTGCCGAAGCTCCCCAAAACGGAATCTGTAACAAGCCATTGCTTTGCAAACAGTCTTACGAGAGGTGATCTGTCATCGATGATGTGGTTTACAGGAGCGCTCAATGAGTTGAGTGTTGTTCCTTACAAGGTGAAGGTATCGTACTGCACCTTGAACTTCCGCGATGTGATGGTTGCCAGTGGTAGATTATCGTCGGATGTTAGCTCGTCCCATCGATTGGATGAAGATTGTGAACTTGGTTACGAGTATGCTGGAGTGATGGAAGATGGCAGACGGGTTATGGGAGTTGTTCCCTCAGGAGCATTATCTTCAATTGTTAATGCAGATCCGTTATTGCTCTGGTATGTTCCAGACGATTGGAGCCTAGAAGAAGCCTGTACTGTTCCAATTGTTTATTTAACCGTATTGGTGTCATTTTTCTTAAAAGCTCACATACGAAAGGGCCAGTCGGTTTTGATCCACGCCGGAAGCGGTGGTGTTGGATTGGCTGCAATACATATCGCTTTCCGATATGGTTTGGAGGTTTACACCACAGTTGGTACTGCGGAAAAACGCAAATTTCTGTTGTCGATGTTTCCTAAGCTAAAATCGGAAAACATCGGAAATTCGCGAGACAGTTCCTTCGAGCAGATGGTCAAGCTGAGAACCAAAGGCAAGGGCGTAGACTACGTGTTGAACTCGCTGGCTGAAGAGAAGCTGCACGCTTCAGTAAGATGTCTGGCGAAAGGTGGCCACTTTTTGGAAATCGGCAAGTTCGATATGGCGAAGGATTCACAGGTCTCTCTGGATCAATTTAAAAAAGGCATCACATTTTCAAGCGTGTTGTTGGACGCTGCCATTAGAGATACTGATACTGTGAAAAAG

>Exon_5

GGTATGCATGATATGTTTCAGAGGGCAATAGAGGACGGAATGATAAAGCCTCTGAATACAACTGTTTACGAAGCTGGCGAGCTGGAGAAGGCGATGAGATACTTGGGCAGTGGAAAGCATACCGGAAAAGTTGTGATCAAGCTTCGTGAAAATGAAACCGATGTCCAAACATTGCCCATAACCTACTTCCCAAAAGTGTACTGTGATCCAAACCAAGTGTACATAATCATCGGAGGCTTGGGAGGCTTCGGTTTGGAGCTGGTCGATTGGCTGGTGCTCAGAGGTTGTAGAAAGCTGGTTTTCAGTTCAACTCGAGGCATCTCCAAGCCATATCAGGAATACAGAATC-AA

*Nucleotides AA from exon_5 and A from exon_6 code for amino acid “K”*

>Exon_6

A-ATTTGGAACAGCTATGGGGCTCACACGCATATTTGCATGGCAGATGTTACCACAGAAGATGGATGTCGCACACTCATCGAAGAAGCTTCCCAAATGGGCACGATTGCAGCGATCTACAACTTGGCAGTCAAGCTACGGGACGCCATATTGGAGAATCAAACAGAGGAGAAGTTCGCGGAATGTTTAGCACCGAAAGCAAAGGCTACCGAGCACCTGGATCGAATCAGTCGAACAAATTGTCCTGGCTTGAAGCATTTCGTTGTGTTCTCCAGCGTGTCCTGCGGTCGCGGAAATGCAGGTCAAAGCAATTACGGTATGGCAAACTCGGTGATGGAACGGATAATTGAGCGAAGACATGCGGATGGTCTTCCCGGCAAGGCCATTCAATGGGGTGCAATAGGAGAGGTTGGACTGGTGGCAGACATGGCCGAGGATAAGATCGATTTGGAGATCGGAGGAACATTGCAGCAACGTATTTCATCGTGTCTACAGGAGATGGACTACTTGCTGACGTGTGAAGCTCCGCTTGTGGCCAGTATGGTGGTTGCGGAGAAGAGAGGAGGTGGATCGAAGAACGTTGTCGAAGCTATGATGCACATTATGAATATCAGAGATTTGAAATCGTTGTCAATGGAGAGTACTCTGGCCGACATTGGAATGGACTCGTTGATGGCGGTAGAAATCAAACAGGTTTTGGAAAGAGATTTCGACATTGTTCTGTCTCCGCAAGAATTGAGAACACTCACTTTTGCTAAATTAATTAGAATGGTTGATGAGGTGAAGATGGCTGAGAAGAATCGTGAAGCAGATGTTCCAACTCTCGGCTTACATATGTTGATGCGGAACCTCGGCAACGAGGAAACTAGCGAATCTACGCTCCTACGTGTACCATCCGCTGGTGAAAGTGGTTGTCCAATAATTTTTATTCCAGGCATTGAAGGTGTTGCTGGTAATGCTATAAAATCTGTTGCAGAGCAACTGAAGGCTCCAGTATACATGCTTCAGTTGACTAGTACACTGGATTGTGAGTGCGTTTCAGCTATTGCTGATGGCATAATCGATGAGATAAGCTCGTTGGCGCTCAGTGAATATGCCATTGTTGCTTACTCGTTCGGAGCCTTCGTGGGAATCGAGTTAGCGAGGCAATTTCGGAAGCGCGGAATTCGTGGAAAGCTTCTGCTGCTAGACGGTGCTCCAAAGTTCTTGAAATTGCTCGCTCTCAAACAACTAAACTACAACACATCAGACGAAGAAATTCAGAAGCTTATAATTTCCGCAATGATTGCCATGGCGTTCAATCACCAATCGATTGAGCGCATTTCTTCGACCATGCAAGCGCAATCGTTCGATGATCAAATTGAACAGTTGATCGAGCTGGGAGCGGAACAAAGTCCGTACTCGGCGGACTACACGAGGAAAATGACTAAAGCCCTGTTCAGGCGGTTGAAAATGGCAGCCCTGTTCGACCTGGATCAATGCGAGCCATTGGATGTCCCAATCACTTTGGTTCGACCGACGGATGCATCTTTTTCGGACATTGAAGAGGACTACGCCCTTTCGCAGTGCACCACCGGTCCCATTGCGCTTCGGATGGTCGAAGGCACTCACATGTCGATGGTGGAGAATCCGGCTCTGCTCGGGTTGATTAACAGTTGGTATTCGTGATATGGTTTAACCAGTGATAGTAACGTGTAATCATTCATTTCAATAAACGAGTTGGTCTCTTTTGTTTGATTTCTTAGTAA

**LOC5573927_amino_acid_sequence**

>Exon_1

MPSPNIQSVGSDESVVISGIAGRFPQSDDVREFARNMYEKRDLVDDKETRFTHAMQAIPRRFGKINNLQKFDRQFFGYDRRQRDTMDPQQWMALEHAFEAILDAGINPESLRGTRTGVFCGVCFSESGARMYYSALGNAEKGMGLQ-A

>Exon_2

FNKSYLALRVSYTLDLNGPSVVVDTACSSSMYALDVAYRSILNGDCEAAIVTGSNLTLHPFISYQFALLGVLSKDGYCRPFDKEASGYTRSEAVCAIFLQKSKNAKRIYCHVVHSKTNCDGFKAEGITYPSGLVQQQLLREIYDEVGISPIQVDYVEAHSTGTVVGDPEECDAIDKVYCKGRDTALLVGSVKSSIGHTEAAAGLCSIAKCIIAMEYDSIPPNINYTECRKDIPALVEGRLKVVTEATPLPGPLIGINSFGFGGANAHTLLCRNLKEKKNNGVPEDDLPRIVLWSGSTREAVECMLQDISQRPLDAEFIGLMHNIEKQPIQGHRYRGFGVYQKDGERASILKESCIERVKLESLPTVAIFGGINVGWKQDLDSLRKISSVDAFYTMCKEVLKTLKFDLYKAHPHEKDLLYNMVGSTVLQLSIIDLLVSSGVALDFYGGHSVGQFICAYIDRCLTLEQVLKLAFWHGLVYSDCHALCESSAFVRMNAKLNQLPLKNIIKNSASSFGILTSSERNIIEQVHQLKSSGFLAEELSFLCFHSAPACSGTLSNTLRQTVNVVLSKIIEPSNKWITARLPQGCSIFHSPALHDASSIVCMLDKIPSHTNVLQLDSAQTCEKVLQMMHRHSRCLTPSSESSDAICRLLSVIG-Q

>Exon_3

LHLISQNLDVAHLYPPVQFPVSRGTPMIAPLVHWDHREDAHIVKYTWDQTAKSRVGTVNISVSSQDWKHIEGHNIGGRVLFPATGYLKLVWDHVAYLAHCDLEDFAIEFEDVRFLRATNITKGQTIRFRVAVEEVSGFFE

>Exon_4

VIEGDTIVVTGFARQWNDSVQESFREERSSVAVLRTKDFYKELRLRGYYYSDQFKSVLESKADGSLAKIQWKGNWVPFLDCLLQVGIIATDSRQLMVPTAIEKLSVSPRAHMEMIERNEDGTEFFTMRTCLLTNTSICGSVMIRNLRASSVNRRNPPGNPVLEIYQFVPFHSDHYIPTVEAIRIIVQLALENAPTFTVNVTEIHHKDSQPIVQLFGDAVADLPLVQSNLTLLSTKSIELDDVTVQDEKLSGLTNQLFLITNSKLEDNTFIQEAVGCLVDRGFIIARSTNPAELTVPTQFTLVASFRVDQSETLICLQRKAFAESPIVVKLDSVEMSWLTNLKRAVKSRSVLLYAQNDPTSGIIGLVNCIRKEPKMQDVRCVFIDDPKAPPFMISDPFYKNQLDLGLAINVLRNGVWGSYRHQLLPKLPKTESVTSHCFANSLTRGDLSSMMWFTGALNELSVVPYKVKVSYCTLNFRDVMVASGRLSSDVSSSHRLDEDCELGYEYAGVMEDGRRVMGVVPSGALSSIVNADPLLLWYVPDDWSLEEACTVPIVYLTVLVSFFLKAHIRKGQSVLIHAGSGGVGLAAIHIAFRYGLEVYTTVGTAEKRKFLLSMFPKLKSENIGNSRDSSFEQMVKLRTKGKGVDYVLNSLAEEKLHASVRCLAKGGHFLEIGKFDMAKDSQVSLDQFKKGITFSSVLLDAAIRDTDTVKK

>Exon_5

GMHDMFQRAIEDGMIKPLNTTVYEAGELEKAMRYLGSGKHTGKVVIKLRENETDVQTLPITYFPKVYCDPNQVYIIIGGLGGFGLELVDWLVLRGCRKLVFSSTRGISKPYQEYRI-K

>Exon_6

IWNSYGAHTHICMADVTTEDGCRTLIEEASQMGTIAAIYNLAVKLRDAILENQTEEKFAECLAPKAKATEHLDRISRTNCPGLKHFVVFSSVSCGRGNAGQSNYGMANSVMERIIERRHADGLPGKAIQWGAIGEVGLVADMAEDKIDLEIGGTLQQRISSCLQEMDYLLTCEAPLVASMVVAEKRGGGSKNVVEAMMHIMNIRDLKSLSMESTLADIGMDSLMAVEIKQVLERDFDIVLSPQELRTLTFAKLIRMVDEVKMAEKNREADVPTLGLHMLMRNLGNEETSESTLLRVPSAGESGCPIIFIPGIEGVAGNAIKSVAEQLKAPVYMLQLTSTLDCECVSAIADGIIDEISSLALSEYAIVAYSFGAFVGIELARQFRKRGIRGKLLLLDGAPKFLKLLALKQLNYNTSDEEIQKLIISAMIAMAFNHQSIERISSTMQAQSFDDQIEQLIELGAEQSPYSADYTRKMTKALFRRLKMAALFDLDQCEPLDVPITLVRPTDASFSDIEEDYALSQCTTGPIALRMVEGTHMSMVENPALLGLINSWYS

**AaFAS-like LOC110675236**

**XM_021839685.1 PREDICTED: Aedes aegypti fatty acid synthase-like (LOC110675236), mRNA**

>Exon_1

ATGCCGTCGCGTAGTATCCAATCGGTTGGTTCCGATGAGAGTGTCGTGATAACCGGAGTGTCCGGTCGTTTCCCGCGTGCGAACAATGTTGGCGAGTTTGCTCGCAGTTTGTACGGAAAGGAGGACCTCGTGGATGACCTGGAGACCCGTTGGCAACACACGATGCAGGACGTTCCACGGCGTACCGGGAAGATCGGAAACATGGAGAAGTTTGACGCAGACTTCTTCGGAGTCAGTCGGATCGAGCGGGACACGATGGATCCACAGGTGCGCATGACGATCGAGCATGTCTACGAGGCTGTGCTTGACGCTGGATTGAATCCATTGACCCTGCGAGGTTCGCGGACCGGGGTATTCAGTGGAGTGTGTTTCTCAGAAACTGAAGTGTGCATGTACTACAGAGCGTGTCCTCCCAAGGGGCTTGGTTTGTTG-GG

*Nucleotides GG from exon_1 and G from exon_2 code for amino acid “G”*

>Exon_2

G-TGTGCCAAATCGCAGATTCCCAATCGTGTTTCTTACTTGCTAGATCTAAGGGGACCTAGCTATGTGCTGGACACTGCTTGTAGTAGCTCAATGTACGCCTTGGATGTTGCGTACCGAAGTATGATGAACGGCGAATGCGATGCGGCAATTGTAACGGGGGCTAATCTAACACTGCATCCCTTCATTACGTACCAGTTTGCAATGTTGGGAGTCTTGGCAAAGGATGGATATTGTCGGCCTTTCGACAAAGACGCAACGGGATACACGCGTTCAGAAGCCGTCTGTGCTGTATTTCTACAGAAAGCAAAAGATGCGAAGCGTGTTTATGGTCACATAATTCATTCAAAGACCAATTGTGACGGATTCAAACCGGAAGGAATAACTTATCCTTCAGGATCAGTTCAACAACAGTTGCTAACTGAGTTCTACAACGAGGTAGGAATAAGTCCTACAGAGGTCGACTATGTAGAAGCACATAGCACTGGAACGTTTGTAGGAGATCCAGAAGAATGTGACGCTATAGATAAAGTGTATTGCACTGAACGAACCGATCCCCTGCTTGTGGGGTCTGTAAAATCCAGTATTGGCCATACGGAAGCTTCAGCAGGAGTATGCTCCATCACAAAATGCATTATCGCCATGGAGAATGGCCTTATTCCTCCAAACATCAATTATACTGATTACAGGCCTACCATTCCGTCTTTGGTTGAAGGTCGTTTGAAGGTTGTTACAGATGCAATGCCACTTTCGGGCCCATTGGTTGGTATCAACTCGTTTGGATTCGGAGGAGCAAATGCTCATGCTTTGTTATGTCGCAATTTGAAAGAGAAAGTTAAGAATGGCGTTCCAGACGATGATCTTCCAAGGTTGGTCACATGGTCAGGAAGAACCAGAGAATCTATTGAAACTATGCTTTACGACATTGGCCAGCGTCCGTTGGATGTTGAATTCATTGCTCTTCTGTTTAACATTCAGCAGCAGCCTACTCCTGGTCATCGGTATCGAGGATTTGGTATTTATCAGAAGAATGGAGATCGTCCAGCTGTACTACAAACATCATCCATCGATCGAGTGAAGCTTGATGCCATTCCTGTGGTTGCTGTTTTCGGAGGAATTAACATCAACTGGAGAAAAGAATTAGATGCTCTGCGGCAATTCTCTGTAGTAGAAGATACATTTGTTAAATGCAGCGGAATTCTACGATCGCTGAAGTTCGATTTGCACAAAAGATCGTCAGGAAGAGAAAGCATATTGTACAATATGGTTGGATCGACCATTCTTCAGTTGTCTATCGTTGATTTGCTGAGCTCGATTGGAGTGAAGTTTGACTTCTATGGAGGACATTCCATTGGCCAATTCACTTGTGCATACATTGACCACAATTTGAACCTAGAACAAGTTCTTCGTTTAGCTTTCTGGCATGGATTAGTGTTTTCTGACTGTCATGCGGTATGTGATCGTACTGCGTTCGTACAAATCAATTCACAGCTGAAACAGCTTCCTTTGGAGAATCTTATCAAGGATAGTGCAACCAACTTTGGTATTTTGAACGCAAATGAGAAAGTAATGGTGGAACAGATTCGTCAACTGAAGTCGTCCGGTTTGATTGCTGAAGAGTTATCGTTTTTGGATGTGCATGCAGATTCAACGAAAAGTTCATCGCTTGCAAATAAACTACGACAAACCGTGAATACCGTTTTGAGCAGAACCATTTTGCCCAGTGATAAATGGATTACTTCAGCATTGCCTCAGACGTCTTCTATATTCCATTCACCGAAACTGCACGATGTTACATCTATAGTTAATTTGATTGAGAATATTCCGCATCATTCACATATCGTAGAGTTTGGCAGCTCACAATCGTGTGAGAATGTGCTTCGGTTTTTAAATCATAACTCGAGTTACATTCCATCTGGATCTACCGCATCGGATACGATCAGTCAATTGTTGTGTCAAATAGGACAGTAAGTGTTT

**LOC110675236_amino_acid_sequence**

>Exon_1

MPSRSIQSVGSDESVVITGVSGRFPRANNVGEFARSLYGKEDLVDDLETRWQHTMQDVPRRTGKIGNMEKFDADFFGVSRIERDTMDPQVRMTIEHVYEAVLDAGLNPLTLRGSRTGVFSGVCFSETEVCMYYRACPPKGLGLL-G

>Exon_2

CAKSQIPNRVSYLLDLRGPSYVLDTACSSSMYALDVAYRSMMNGECDAAIVTGANLTLHPFITYQFAMLGVLAKDGYCRPFDKDATGYTRSEAVCAVFLQKAKDAKRVYGHIIHSKTNCDGFKPEGITYPSGSVQQQLLTEFYNEVGISPTEVDYVEAHSTGTFVGDPEECDAIDKVYCTERTDPLLVGSVKSSIGHTEASAGVCSITKCIIAMENGLIPPNINYTDYRPTIPSLVEGRLKVVTDAMPLSGPLVGINSFGFGGANAHALLCRNLKEKVKNGVPDDDLPRLVTWSGRTRESIETMLYDIGQRPLDVEFIALLFNIQQQPTPGHRYRGFGIYQKNGDRPAVLQTSSIDRVKLDAIPVVAVFGGININWRKELDALRQFSVVEDTFVKCSGILRSLKFDLHKRSSGRESILYNMVGSTILQLSIVDLLSSIGVKFDFYGGHSIGQFTCAYIDHNLNLEQVLRLAFWHGLVFSDCHAVCDRTAFVQINSQLKQLPLENLIKDSATNFGILNANEKVMVEQIRQLKSSGLIAEELSFLDVHADSTKSSSLANKLRQTVNTVLSRTILPSDKWITSALPQTSSIFHSPKLHDVTSIVNLIENIPHHSHIVEFGSSQSCENVLRFLNHNSSYIPSGSTASDTISQLLCQIGQ

**AaFAS6 LOC5573930**

**XM_021847677.1 PREDICTED: Aedes aegypti fatty acid synthase (LOC5573930), mRNA**

>Exon_1

TTTCTAGATCTATGAAGGAGATTCAGTTGTTGTGACCGGGATTGCAAAAATGCTGGAAAGCGAAAACTTAACTGATATTCAAGAATCCTCAAATCCAGCAGTAACATTGAAATCGAGAGACTTCTATAAAGAGCTACGTTTACGAGGATACCATTACACAGGTCTATTTAAATCGGTGTTGGAAGCAAAATCTGACGGAACCATGGCTAAGGTCCAATGGAAAGGTAATTGGATGGCTTTTCTTGATTGCCTGCTACAAACTGCAATAATTGCGATTGATACCAGATCGTTGATGGTTCCTACGGCAATCGAAAAGCTTTCAATTGCACCGAAAGCCCATTTGGCAATGATGGAACGTGAGGGAGAAGATTGTGAGTTCTTCACGATGAGAAGCTGTCCAAAAACTAACGTTTCGGTTTGTGGCGGTATTATGCTTTGCAACCCTCGGGCCAGCAGCGTAGGACGTAGAAACCCTCCAGGTATTCCAGTTCTAGAGACTTATCAATTTGTTCCGTATCACGCCGATGATCAGGTTCCGACTCTGGAAGCAATTAGAATGTGCGTTCAGCTTGCGCTAGAAAATGTCCCTACTCTATCAATAAATGTGACAGAAATCCACAGTGAGAGGATTCCGGTTGTCGCACATTTGTTCGGAGAAGCAATAGCCGATCTTCCATTGGTTAAAGCCAATCTGACGGTTCTGGCTAAAACAGAAATTGAGTTGGAATACGTGACTGTCAAAGTGGAAAAGCTATCTGATCAATCAAATCAACTCTTTCTGATCACTGACAGCAATTGGAGTGAGCCGAATTTCCTACAAGACGCTGTTGGTCGTCTCGTAGATGGAGGATTCATCATTGTTCGTGAGAAGTTAACCTTCAAGTTAGATGACCTAGATGTACCAGAAGAACTGAACATGGTGGCCTCCTTTAGAGTCGATCAAGAAGAGACATTTATTTGTCTGCAACGCAAAATCAAGGGATTCAATGATACTCCAGCAGTTATCCAGGTTGATTCATCCGACTTCAGTTGGCTAGCTGTTTTGAAACAGGCTGTCAAAGTCCGACCTGTGATCCTATTCTCACAGAATGACTCAGTTTCCGGAGTTATCGGTTTGGTGAACTGTATTCGCAAAGAACCCAAAATGCAAACAGTACGATGCGTACTAATTGATGACCAAAATGCTCCGGAATTCTCTCTCAGTGATCCATTCTACAAGAACCAGTTAGAGCTCGGACTGGCAATTAACGTATTGCGAAACGGAGTGTGGGGTAGCTATCGCCATGCCTTAATATCGAAGAAACCAAAAACGGAACCTGTATCCAAGCATTGCTACGCAAACAGCCTGACAAAAGGTGATTTGTCTTCGATGATGTGGTTCACGGGAGCCTTCAACGAGTGGGATGTTGTACCAAATAAGGTAAAAGTGTCGTACTGCACTTTAAATTTCCGTGATGTGATGGTTGCTACTGGAAGATTGTCATCGGACGTGAGTTCATTCAGTCGACTGGAAGAAGAATGTGAGCTTGGTTATGAATATGCGGGAGTTACGGAAGACGGCAAGCGGGTGATAGGTGTATTGCCATTAGGGGCTCTATCTACGATGGTTAACGCGGATTCGACTCTTATTTGGGTTATACCGGATAGTTGGAATGCACAGGATGCTTGCACAATTCCAATTGTATATACCACGGTTTTGGCAGCATTTAAACTAAACGCTAATGTGAAGAAAGGGCAGTCGGTATTAATCCATGCAGGAAGCGGAGGTGTTGGGTTAGCTGCAATCAACCTTGCGCTTGCATATGGAATGGAGGTATTCACAACTGTTGGATCCGATGAGAAGGTCAATTACCTGTTGAACGAGTTTCCTTCTCTAAAACGAGAAAACATAGGAAATTCGAGAGATTTATCATTTGAACAAATGATCAAATTAAGAACCAATGGAAGAGGAGTGGACTACGTGTTGAACTCGTTAGCTGACGAGAAACTGCAGGCGTCTGTGAGATGCTTGGCGAAAGGCGGACATTTTCTAGAAATCGGAAAATATGATATGGCAAGAGATTCGCAACTATCATTGGAGCTCTTTAAAAAGGGAATATCATTCACAAGTGTGATGTTGGATTCGGCAATCAGAGATAATTATACTCTAAAATTG

>Exon_2

AGCTTACATAAATTATTGGATGAAGCAATCAAGTCTGGTATAGTGAAACCACTGAAAACCAACGTTTTCGATGCTGCTGATTTGGAGAAGGCAATGAGATTTTTGGCGAGTGGAAAACATATGGGCAAAATAGTGATCAAACTGCGTGAACATGAGAATGACGCTGAAACACTTCCAATTACACATTTTCCACACGTCTTCTGTAATCCAGATCAAGTTTACGTAATCGTTGGCGGCTTAGGTGGTTTTGGTCTGGAATTGGCGGATTGGCTTGTCCTTCGTGGGTGCAGGAAGCTGGTACTCAGCTCTAGTCGAGGTATCACCAAGCCATATCAGGAATACAGAATT-AA

*Nucleotides AA from exon_2 and A from exon_3 code for amino acid “K”*

>Exon_3

A-ATATGGAATAGTTATGGCGTTCATACTCATATCTGTACTGCCGATGTTACCACAATGGACGGTTGTCGTGTCCTCTTGAAAGAAGCTTCTCGGTTCGGTTCTGTAACGGCTATCTACAACTTAGCAGTGCAGCTTCGAGATGCCATATTGGAGAATCAGACTGTGGACAAGTTCGTGGAGTGTATGGCACCTAAGGCTACTGCAACTGAATACCTTGACAAGGTCAGTCGTGAGATGTGTCCTCATCTGAAGCACTTCATAGTGTTCTCCAGTGTTTCCTGTGGTCGCGGTAACGCAGGACAGAGCAACTATGGTATGGCAAACTCTGTGATGGAGCGTATAATTGAACGGAGAAACGCGGACGGCCTTCCCGGAAAGGCGATTCAATGGGGAGCTATCGGTGAGGTTGGACTTGTAGCAGATATGGCAGAAGATAAGATCGATTTGGAAATCGGAGGAACGTTGCAACAGCGCATATCATCATGTCTTCATGAGATGGATTACTTGCTGACATGTGAAGCTCCTCTTGTGGCCAGTATGGTGGTTGCGGAAAAACGAACTGCAAGCGGATCAAAGAATGTCATCGAAGCTGTCATGAACATAATGAGTATAAGAGATCTGAAATCGGTGTCGATGGAGAGTACTTTGGCCGATATTGGAATGGACTCTTTGATGGCAGTAGAAATCAAACAGGTACTGGAAAGAGACTTCGACATGGTACTGTCCCCGCAAGATCTGAGAACATTATCCTTTGCTAAGTTGTTGAAAATGGATGAAGAGAAAAAGCAAGCTGCTAAAGATCAGGAGGAAAAGAAGAGTGAAGGATTTGTGATTGGAATGCAAATGTTGTTGAGAAATCTTGGCAACGAAGAAACGAGTGAATCGACGTTATTGCGTTTACCTTCCGCTGATCAAGAAGGCCGTTCACTTTTATTCATTCCAGGTGTTGAAGGCGTGGCCGGTAATGTTTGGAAAGCAATTGCTGCACAGATAAAGTCTCCTGTTTACATGCTACAGCTATCGAACACTTTAGATTGTGATAGCATCCCAGATATCATAGAACGCATCATAGATGAGATCTGTGAAACAATGTTCAATGGTTTTGAAGATATTACAATAGTTGCTTATTCATTTGGAGCTCTGATTGCAATCGAAATAGCTCGATATTTACAGGCAAAAGGTATACGTGGAGAACTTTTACTATTGGATGGTGCACCGAAGTACTTGAAGCAATGGTCACTGAAACAACTGAATAATAATCCATCGGATGGAGAAATACAGAAGCTCATATTACTTGTTTTGATCGCCATGGTGTTCCCAGATCAGCCACCGGAGAAAGCCATGGCTGTATTGGAGATTACGTCATTCGACGACCAAATTGAGAAACTCATCGAATTGGGAGCAGAGCAGAGCGAATATTCACCAGAGTACACAAGAAAGATGACGAAAGCCCTCTGCAGGAGAATCAAAATGGCGGCTCTGATGAACCTGGATGAAGATCAACCGTTGGACCTTCCGATAACGCTGGTTAGACCAACCGATGCCGTTTTCTCGGATATTGAGGATGACTACGGGCTGTCAAGTTATACCACAGGAGCTATAACACTACGAATGGTCGAAGGCAATCATGTATCTATGTTGGAAAATGCCGATCTAATAGGAATGATCAATAATTTCAGTTGTCAGAATAATCGTGCGGGTTAAATGCTAACTACCTTCTGAAGTTGTGTTTTTTATATCTGAAAATGTTGTGCTATACCGTAGGGCTTGGTAGAATAGTATCTCTCTGTAGCGTAAATGTTCCAATAGTGAAAATTCAAGAAGGACAATAAACGTGCTTGTGTTTT

**LOC5573930_amino_acid_sequence**

>Exon_1

MLESENLTDIQESSNPAVTLKSRDFYKELRLRGYHYTGLFKSVLEAKSDGTMAKVQWKGNWMAFLDCLLQTAIIAIDTRSLMVPTAIEKLSIAPKAHLAMMEREGEDCEFFTMRSCPKTNVSVCGGIMLCNPRASSVGRRNPPGIPVLETYQFVPYHADDQVPTLEAIRMCVQLALENVPTLSINVTEIHSERIPVVAHLFGEAIADLPLVKANLTVLAKTEIELEYVTVKVEKLSDQSNQLFLITDSNWSEPNFLQDAVGRLVDGGFIIVREKLTFKLDDLDVPEELNMVASFRVDQEETFICLQRKIKGFNDTPAVIQVDSSDFSWLAVLKQAVKVRPVILFSQNDSVSGVIGLVNCIRKEPKMQTVRCVLIDDQNAPEFSLSDPFYKNQLELGLAINVLRNGVWGSYRHALISKKPKTEPVSKHCYANSLTKGDLSSMMWFTGAFNEWDVVPNKVKVSYCTLNFRDVMVATGRLSSDVSSFSRLEEECELGYEYAGVTEDGKRVIGVLPLGALSTMVNADSTLIWVIPDSWNAQDACTIPIVYTTVLAAFKLNANVKKGQSVLIHAGSGGVGLAAINLALAYGMEVFTTVGSDEKVNYLLNEFPSLKRENIGNSRDLSFEQMIKLRTNGRGVDYVLNSLADEKLQASVRCLAKGGHFLEIGKYDMARDSQLSLELFKKGISFTSVMLDSAIRDNYTLKL

>Exon_2

SLHKLLDEAIKSGIVKPLKTNVFDAADLEKAMRFLASGKHMGKIVIKLREHENDAETLPITHFPHVFCNPDQVYVIVGGLGGFGLELADWLVLRGCRKLVLSSSRGITKPYQEYRI-K

>Exon_3

IWNSYGVHTHICTADVTTMDGCRVLLKEASRFGSVTAIYNLAVQLRDAILENQTVDKFVECMAPKATATEYLDKVSREMCPHLKHFIVFSSVSCGRGNAGQSNYGMANSVMERIIERRNADGLPGKAIQWGAIGEVGLVADMAEDKIDLEIGGTLQQRISSCLHEMDYLLTCEAPLVASMVVAEKRTASGSKNVIEAVMNIMSIRDLKSVSMESTLADIGMDSLMAVEIKQVLERDFDMVLSPQDLRTLSFAKLLKMDEEKKQAAKDQEEKKSEGFVIGMQMLLRNLGNEETSESTLLRLPSADQEGRSLLFIPGVEGVAGNVWKAIAAQIKSPVYMLQLSNTLDCDSIPDIIERIIDEICETMFNGFEDITIVAYSFGALIAIEIARYLQAKGIRGELLLLDGAPKYLKQWSLKQLNNNPSDGEIQKLILLVLIAMVFPDQPPEKAMAVLEITSFDDQIEKLIELGAEQSEYSPEYTRKMTKALCRRIKMAALMNLDEDQPLDLPITLVRPTDAVFSDIEDDYGLSSYTTGAITLRMVEGNHVSMLENADLIGMINNFSCQNNRAG
