## Supplementary figures and images for "Expression of fatty acid synthase genes and their role in development and arboviral infection of *Aedes aegypti*"

### Figure S1 _Chotiwan et al

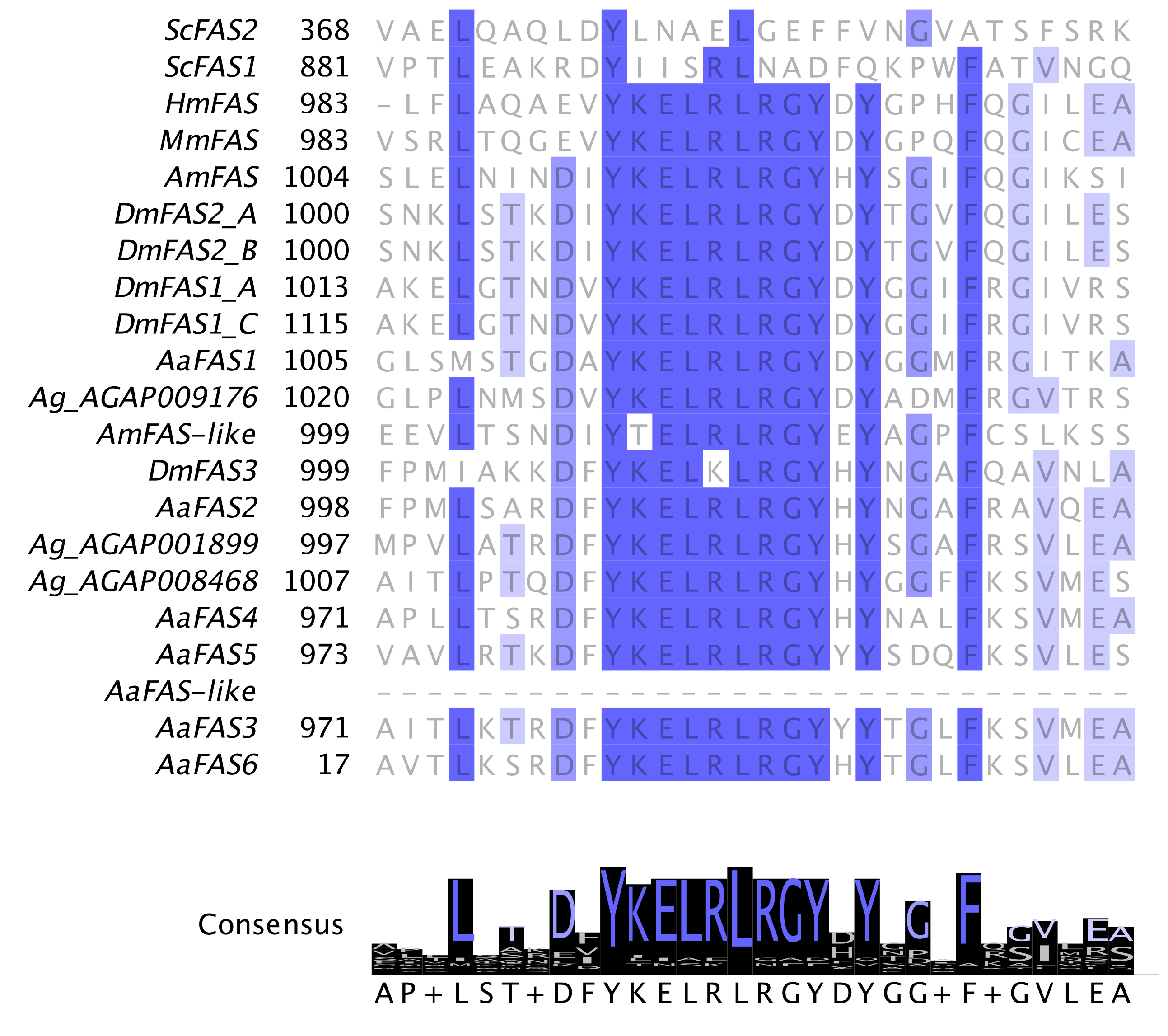

### Figure S2_Chotiwan et al

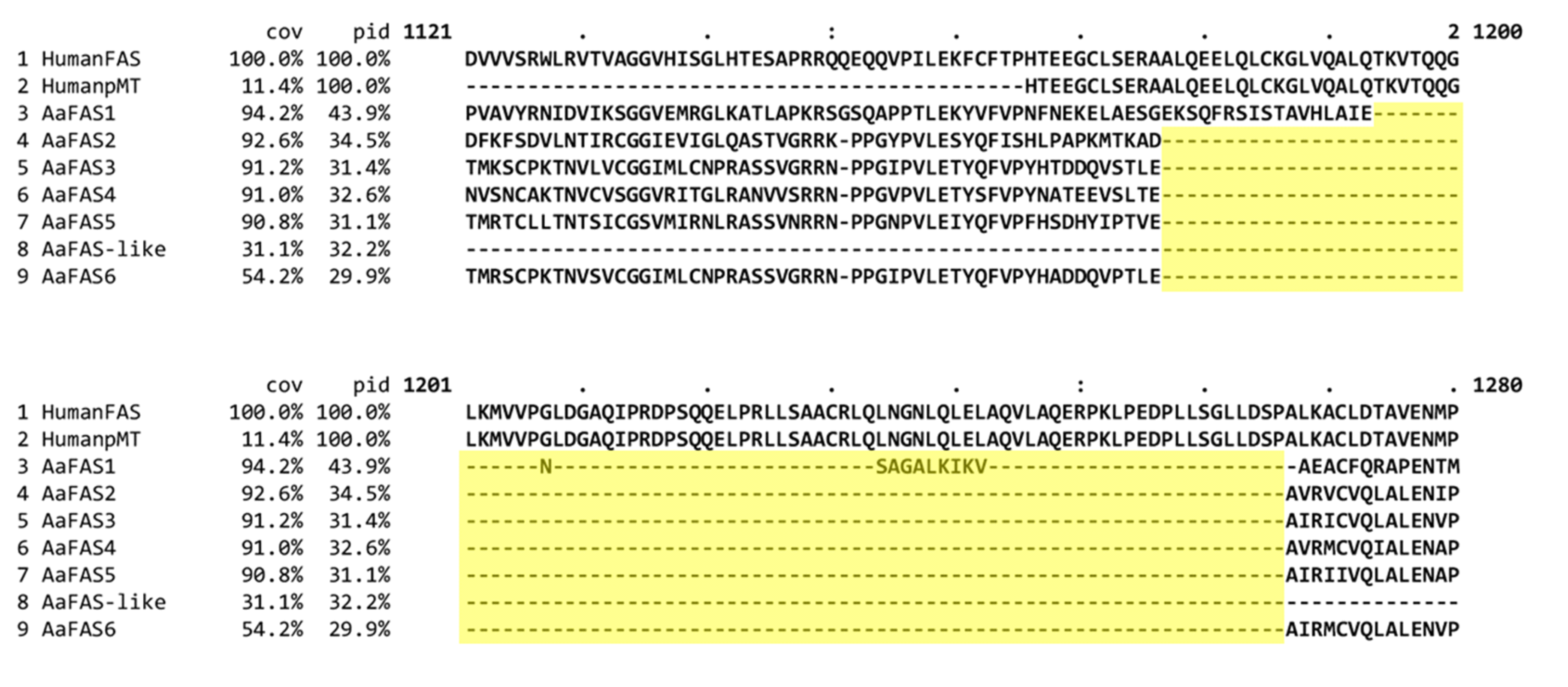

### Figure S3 _Chotiwan et al

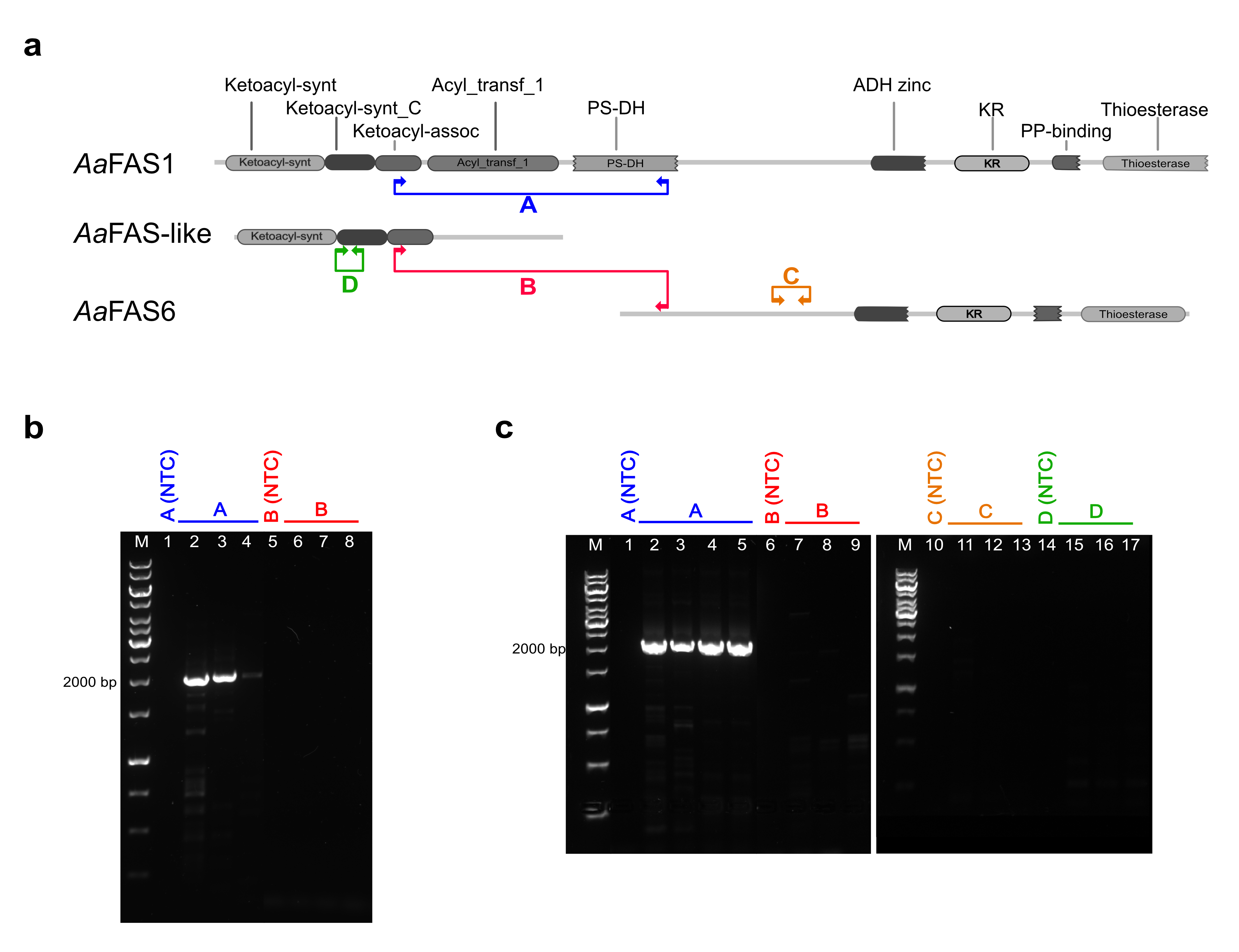
